# Navigating through chemical space and evolutionary time across the Australian continent in plant genus *Eremophila*

**DOI:** 10.1101/2020.11.02.364471

**Authors:** Oliver Gericke, Rachael M. Fowler, Allison M. Heskes, Michael J. Bayly, Susan J. Semple, Chi P. Ndi, Dan Stærk, Claus J. Løland, Daniel J. Murphy, Bevan J. Buirchell, Birger L. Møller

## Abstract

*Eremophila* is the largest genus in the plant tribe Myoporeae (Scrophulariaceae) and exhibits incredible morphological diversity across the Australian continent. The Australian Aboriginal Peoples recognize many *Eremophila* species as important sources of traditional medicine, the most frequently used plant parts being the leaves. Recent phylogenetic studies have revealed complex evolutionary relationships between *Eremophila* and related genera in the tribe. Unique and structurally diverse metabolites, particularly diterpenoids, are also a feature of plants in this group. To assess the full dimension of the chemical space of the tribe Myoporeae, we investigated the metabolite diversity in a chemo-evolutionary framework applying a combination of molecular phylogenetic and state-of-the-art computational metabolomics tools to build a dataset involving leaf samples from a total of 291 specimens of *Eremophila* and allied genera. The chemo-evolutionary relationships are expounded into a systematic context by integration of information about leaf morphology (resin and hairiness), environmental factors (pollination and geographical distribution) and medicinal properties (traditional medicinal uses and antibacterial studies) augmenting our understanding of complex interactions in biological systems.

## Introduction

The large, cosmopolitan plant family Scrophulariaceae *sensu stricto* contains eight tribes including the Myoporeae. Myoporeae contains seven genera, with *Eremophila* being the largest genus in the tribe (∼230 species and 58 subspecies). *Eremophila* exhibits incredible morphological diversity throughout the Eremean (arid) biome, which covers approximately 70% of the Australian continent. Indigenous people of the Australian mainland – the Australian Aboriginal Peoples – recognize *Eremophila* species as important sources of traditional medicine [1–3]. Aboriginal Peoples’ use a range of *Eremophila* species and preparation methods, though use of leaves is documented most commonly. Ethnomedicine can be a key guide to bioactive natural products and potential drug leads [4, 5].

Chemical exploration of *Eremophila* and closely related species was pioneered by Emilio Ghisalberti and colleagues in the 1970s [6]. A plethora of unique and structurally diverse metabolites belonging to the family of terpenoids have subsequently been uncovered [7–14] (Figure 1). Extracts from *Eremophila* species have been tested for anti-viral [15], anti-bacterial [16–18] and anti-cancer [19] activity, and show inhibitory activity of ion channels [20]. Isolated serrulatane- and viscidane-type diterpenoids exhibit anti-malarial [21], anti-bacterial [7, 22–27], anti-diabetic [10, 11], and anti-inflammatory [22, 24] properties. The presence and biosynthesis of interesting metabolites in species of Myoporeae have previously been demonstrated [10–14, 18, 21, 28] and recent phylogenetic studies have revealed complex evolutionary relationships between genera in the tribe [29, 30]. Still only a fraction of species within the Myoporeae have had their metabolites investigated. Based on the rich source of bioactive molecules this group has proven to be, it is expected that further investigation will yield many more novel molecules with potential as pharmaceuticals.

**Figure 1.**
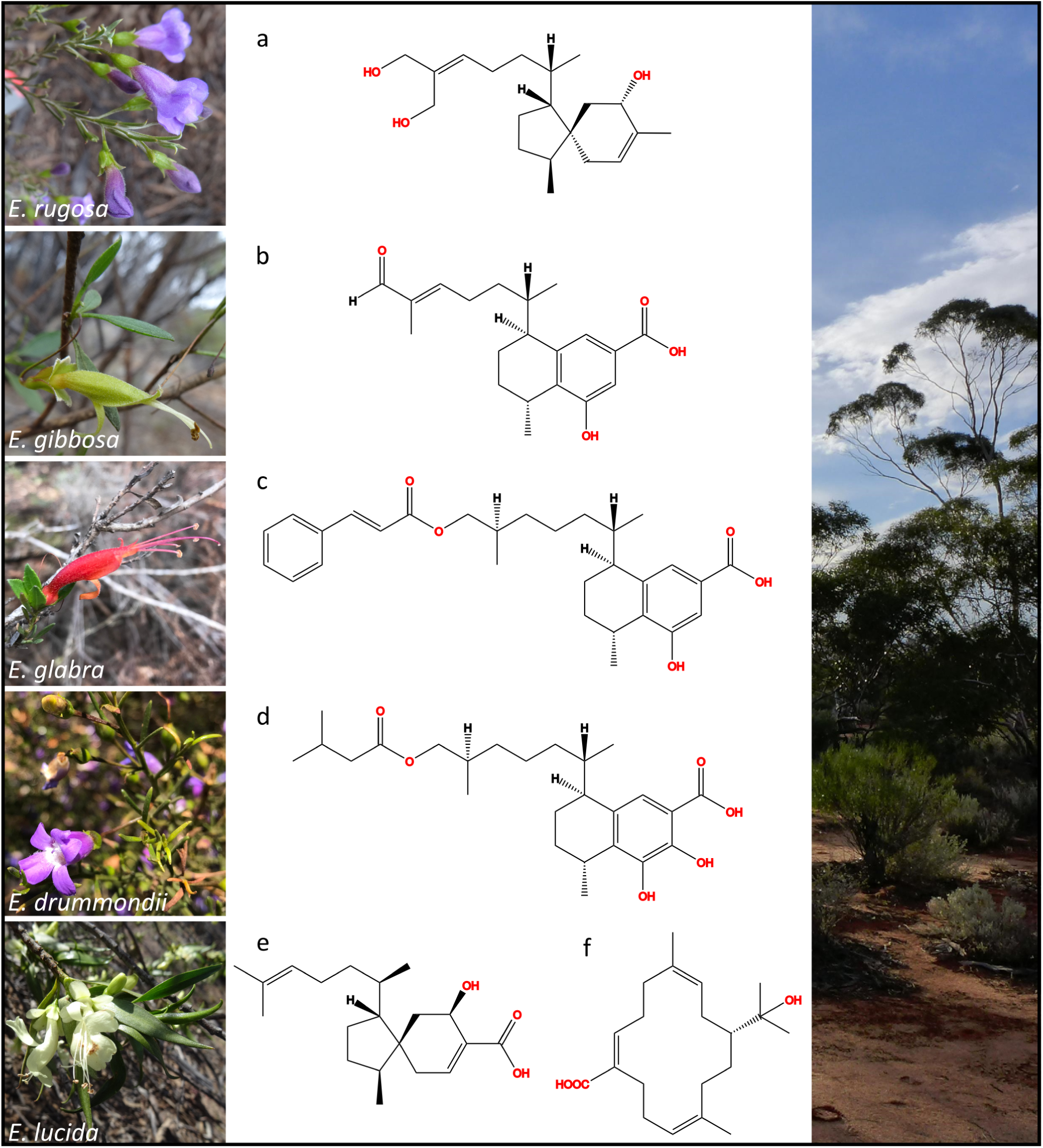
Chemical and morphological representation of selected *Eremophila* species. Images on the left display selected *Eremophila* species. Corresponding diterpenoids isolated in prior studies are displayed in the middle column. The right column shows a typical Eremean habitat in Western Australia. *Eremophila rugosa*, (a) KU036-6-5 [71]; *E. gibbosa*, (b) 8-hydroxy-16-oxoserrulat-14-en-19-oic acid [11]; *E. glabra*, (c) 8-hydroxy-16-cinnamoyloxyserrulat-19-oic acid [11], *E. drummondii*, (d) 7,8-dihydroxy-16-[3-methylbutanoyloxy]serrulat-19-oic acid [11]; *E. lucida*, (e) 5-hydroxyviscida-3,14-dien-20-oic acid and (f) (3Z, 7E, 11Z)-15-hydroxycembra-3,7,11-trien-19-oic acid [10].

Plant diterpenoids represent a rich source of bio-based pharmaceuticals and constitute the foundation of many drug development success stories [31], such as paclitaxel (Taxol) [32], ingenol mebutate (Picato) [33] and forskolin (e.g. ForsLean) [34]. Despite evidence of diterpenoids as high-value compounds, only a few plant species have been thoroughly investigated in drug discovery campaigns [35, 36]. The hunt for novel plant derived pharmaceuticals is not a trivial task given the immense number of unknown metabolites present in any given species. Emerging tools that can help in this challenge are *in silico* dereplication methodologies that can illuminate this chemical “dark matter” found in plants [37]. One such tool is molecular networking, an approach which greatly enhances the dereplication of metabolomics data and allows a streamlined hypothesis-driven targeting of metabolites in contrast to the traditional “grind and find” model [38–40]. Molecular networking further enables integration of functional annotations, such as biological, taxonomic and geographical data [41–43]. This multilayered approach facilitates the dereplication of large metabolite datasets associated with interesting functionalities and thus improves our understanding of biological systems in which they are found.

In this study, we have investigated the chemo-evolutionary relationships in the plant tribe Myoporeae by applying phylogenetics and state-of-the-art computational metabolomics to a dataset of 291 leaf samples of *Eremophila* and allied genera. Information about leaf morphology (resin and hairiness), environmental factors (pollination and geographical distribution) and medicinal properties (traditional medicinal uses and antibacterial studies) is used to bring chemical knowledge gained on the natural products present, qualified by use of a large dataset of 76 reference compounds, into a systematic context that augments our understanding of interactions in biological systems and facilitates targeted drug discovery. This research has been conducted under Australia’s access and benefit-sharing laws which are consistent with obligations under the Nagoya Protocol [44].

## Results

### Chemo-evolutionary relationships in Myoporeae

In this study, molecular phylogenetics using data from high-throughput DNA sequencing was integrated with plant metabolomics utilizing state-of-the-art molecular networking tools to elucidate chemo-evolutionary relationships by exploring metabolite diversity and evolution across the diverse tribe My-oporeae (Fig. S1). Our uniquely comprehensive sampling included 291 specimens, representing six genera and ∼80% of the species in the tribe.

We assessed metabolite diversity in Myoporeae using untargeted high-resolution mass spectrometry (MS^1^ and MS^2^) to generate a molecular network via a feature-based molecular networking pipeline [45]. To focus on the chemically diversified fraction of the Myoporeae metabolome, we used the 100 largest chemical subnetworks with at least 9 nodes; each chemical subnetwork representing a chemical family of structurally related metabolites (Figure 2). Those 100 chemical families altogether comprised 3415 nodes, representing 69% of the full connected network (4936 nodes), excluding singletons (6267 nodes without a structurally similar neighbour node). Within this dataset, the median number of detected chemical families per specimen was 15 and the maximum was 36. Chemical similarity among samples was assessed, based on binary presence/absence information of chemical families, using hierarchical cluster analysis.

**Figure 2.**
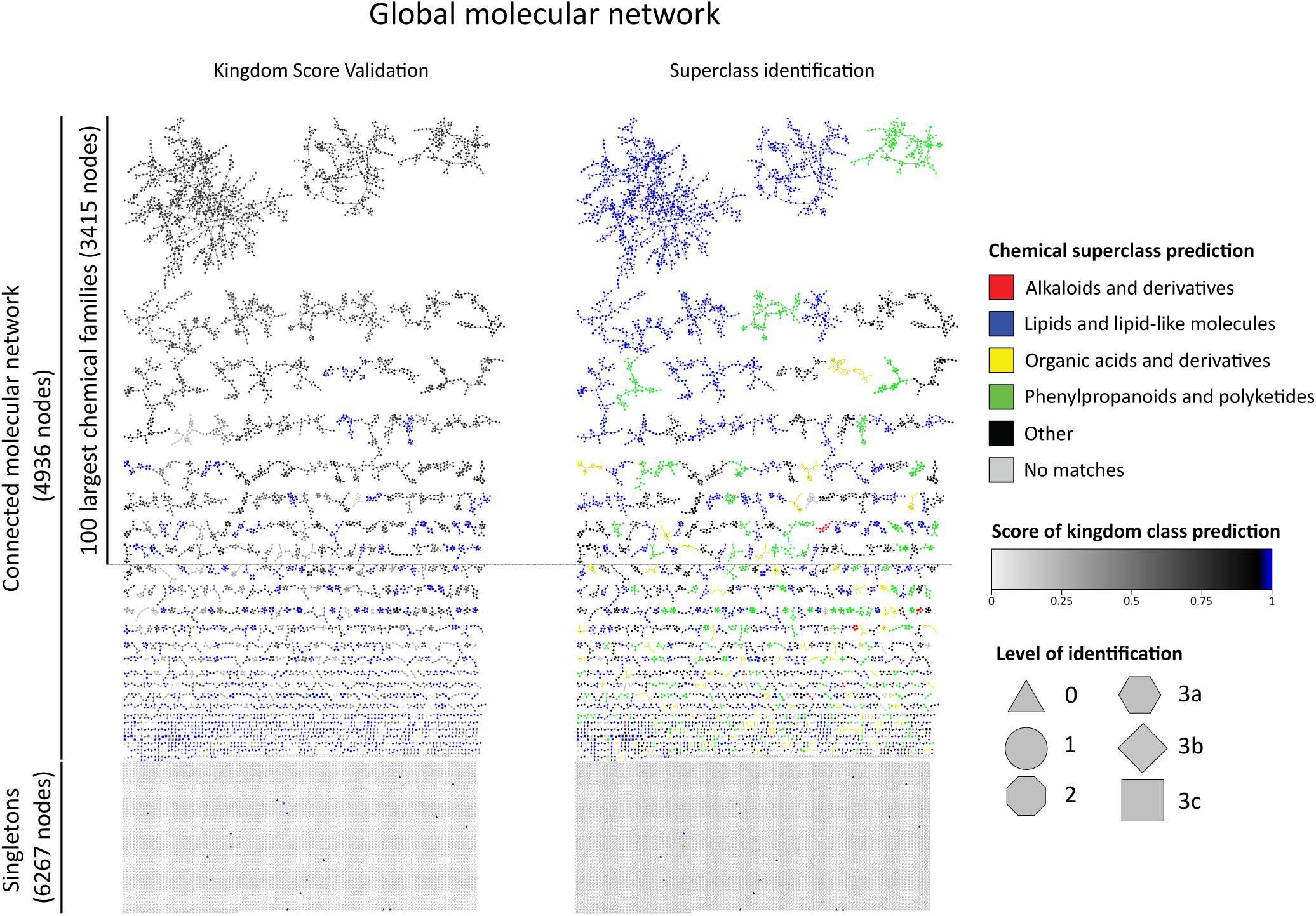
Validation of dereplication in the global molecular network of Myoporeae. The global molecular network of Myoporeae comprises a connected part, with clusters of spectral features that represent structurally-related putative metabolites, as well as singletons, which represent features that did not show spectral similarity to another one. The connected dataset was dereplicated using public, in house and *in silico* spectral libraries and subsequent Network Annotation Propagation (NAP). Identification of nodes took place on six different levels, with level 1: m/z, retention time and MS^2^ match using in house spectral library; level 2: MS^2^ match using in house and public spectral libraries; level 3a-c: Fusion, Consensus or MetFrag algorithm based *in silico* fragmentation database match; level 0: no spectral match. The presented network on the left, displays a kingdom score based assessment with overall high scoring values that support a robust dereplication of the corresponding chemical families. The network on the right side displays NAP-generated chemical superclass prediction for each chemical family, with selected classes highlighted in colour.

Clear phylogenetic patterns, with closely related species having similar chemical profiles, were revealed when metabolic clusters and phylogenetic analyses were compared in the form of a tanglegram (Figure 3). The phylogenetic analysis identified eight major lineages (labelled clades A–H) and the metabolic cluster analysis identified two distinct tanglegram metabolic clusters (TMC) A and B. TMC A includes the majority of *Eremophila* diversity, all allied genera, and both outgroup species (*Leucophyllum*) (Table 1). Morphologically, species producing metabolites characteristic of TMC A are highly diverse, being variously insect or bird pollinated, having hairy or glabrous and resinous or non-resinous leaves. Geographically, species of TMC A are represented across the full distribution of the tribe, throughout Australia’s arid and semi-arid zones, but also including temperate, subtropical and non-Australian members. In contrast, species producing metabolites characterizing TMC B are a subset of *Eremophila* species from three main lineages of phylogenetic clade H (subclades H11, H12, H14). Species producing the metabolites characteristic of TMC B are also morphologically diverse (bird and insect pollinated, hairy and glabrous leaves) and are more likely to have some particular traits: within the TMC B taxa, 83% of the species have resinous leaves whereas this only applies to 31% of the TMC A taxa. Species in TMC B occur largely in Australia’s central arid zone, with particularly high species diversity in the north west (TMC B1) and south west (TMC B2) of Western Australia relative to species associated with TMC A chemistry (Figure S2).

**Figure 3.**
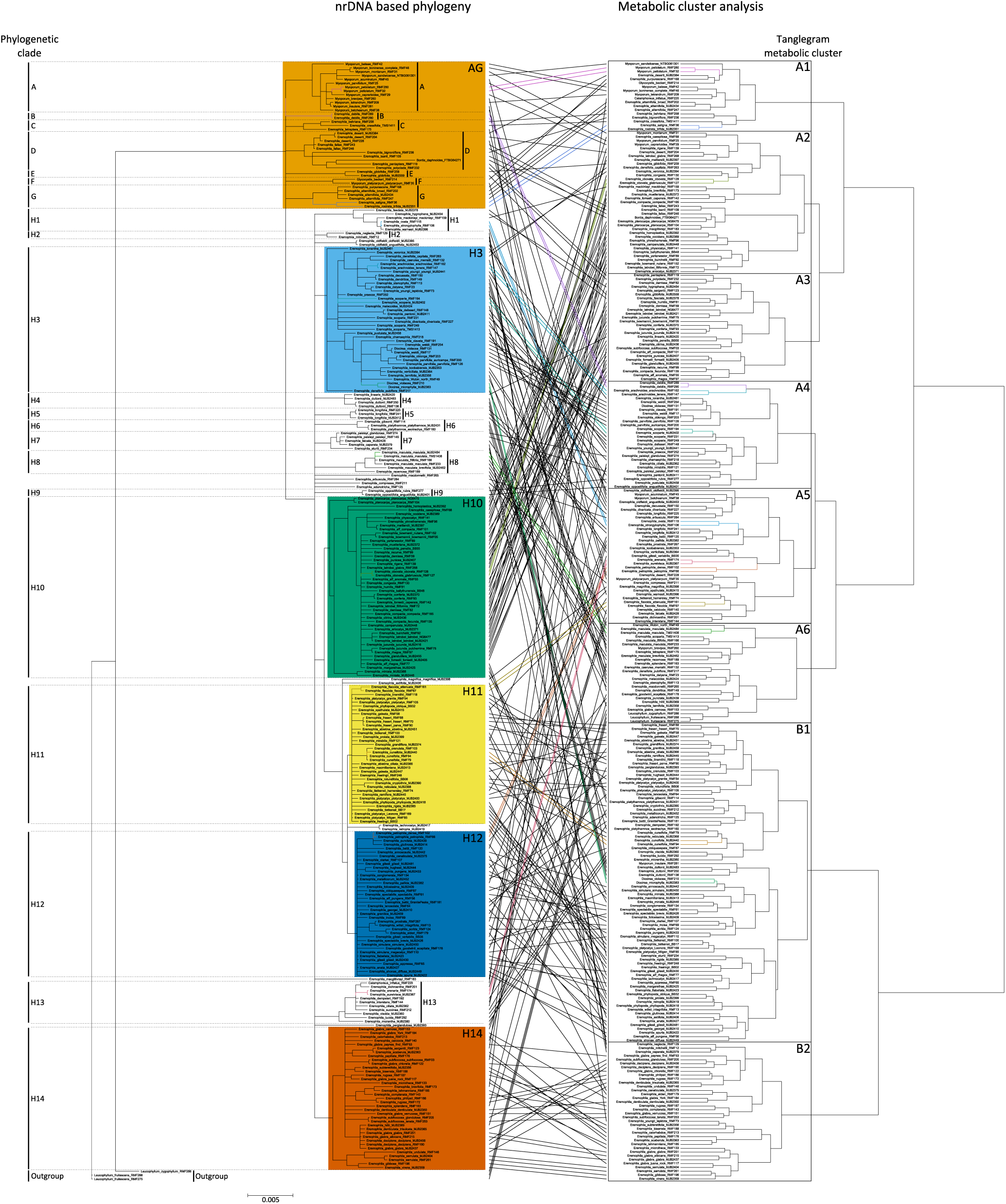
Molecular network-based tanglegram analysis showing chemo-evolutionary relationships in Myoporeae. Tanglegram representation of conjoined phylogenetic and metabolic information from 291 specimens of tribe Myoporeae. Left side displays the Bayesian inference nuclear ribosomal DNA phylogeny. All bifurcating branches have strong support (posterior probability ≥ 0.95 %), all nodes with support <0.95 have been collapsed. All phylogenetic clades are labelled (clades A – H), and major ones are highlighted by color code. The right side displays the metabolic cluster analysis that was conducted based on the presence or absence of the 100 largest chemical families within the generated molecular network of Myoporeae. Therein, tanglegram metabolic cluster (TMC) are indicated. The tanglegram analysis connects same specimens by a line, which is colored when equal clustering of specimens is present in both analyses. _17_

**Table 1.** Summary of morphological, environmental and medicinal attributes between major tanglegram metabolic clusters. Attributes are evaluated between the two dominant tanglegram metabolic clusters (TMC), A and B, which are based on the presence/absence of the 100 largest chemical families found within the investigated Myoporeae associated molecular network.

|  | TMC A | TMC B |
| --- | --- | --- |
| Total no. species | 130 (+ 2 outgroup species) | 85 |
| Genera represented, no. of species in brackets | *Bontia* (1) | *Diocirea* (2) |
|  | *Calamphoreus* (1) | *Eremophila* (82) |
|  | *Diocirea* (1) | *Myoporum* (1) |
|  | *Glycocystis* (1) |  |
|  | *Eremophila* (114) |  |
|  | *Myoporum* (12) |  |
| No. of resinous:non-resinous taxa (duplicate specimens removed) | 45:99 (31% resinous:69% non-resinous) | 86:18 (83% resinous:17% non-resinous) |
| No. of taxa with hairy leaves:glabrous leaves | 90 hairy:54 glabrous (62%:38%) | 76 hairy:28 glabrous (73%:27%) |
| No. of insect:bird pollinated species (duplicate specimens removed) | 111:36 (75% insect:25% bird) | 67:37 (64% insect:36% bird) |
| Documented traditional use | 10 species | 8 species |
| Documented antimicrobial activity | 8 species (of total 51 taxa tested (8+43)) | 25 species (of total 37 taxa tested (25+12)) |

Heatmap analyses integrating the observed chemical space with phylogenetic relationships in Myoporeae were used to explore possible associations between plant chemistry, resin production and geography. This approach shows a clear phylogenetic signal, with specific groups of chemical families (heatmap metabolic clusters, HMC) found to be associated with phylogenetic clades (Figure 4). The HMCs V, VIII, IX and X show strong phylogenetic signatures in the heatmap, and the dereplication approach revealed a dominance of terpenoid-related chemistry in these clusters, typically associated with serrulatane and/or viscidane diterpenoids. The complete lack of these chemical families in outgroup species, yet dominance amongst many clades of *Eremophila* suggests that this chemistry may be unique, and highly specialised within Myoporeae. Specifically, HMC IX chemistry is dominant in phylogenetic subclades H11, H12 and largely absent from other lineages. The HMC VIII chemistry extends to early-diverging lineages (clades A–G) while that of HMC X extends to clade H14. HMC V is largely unique to phylogenetic clade H14.

**Figure 4.**
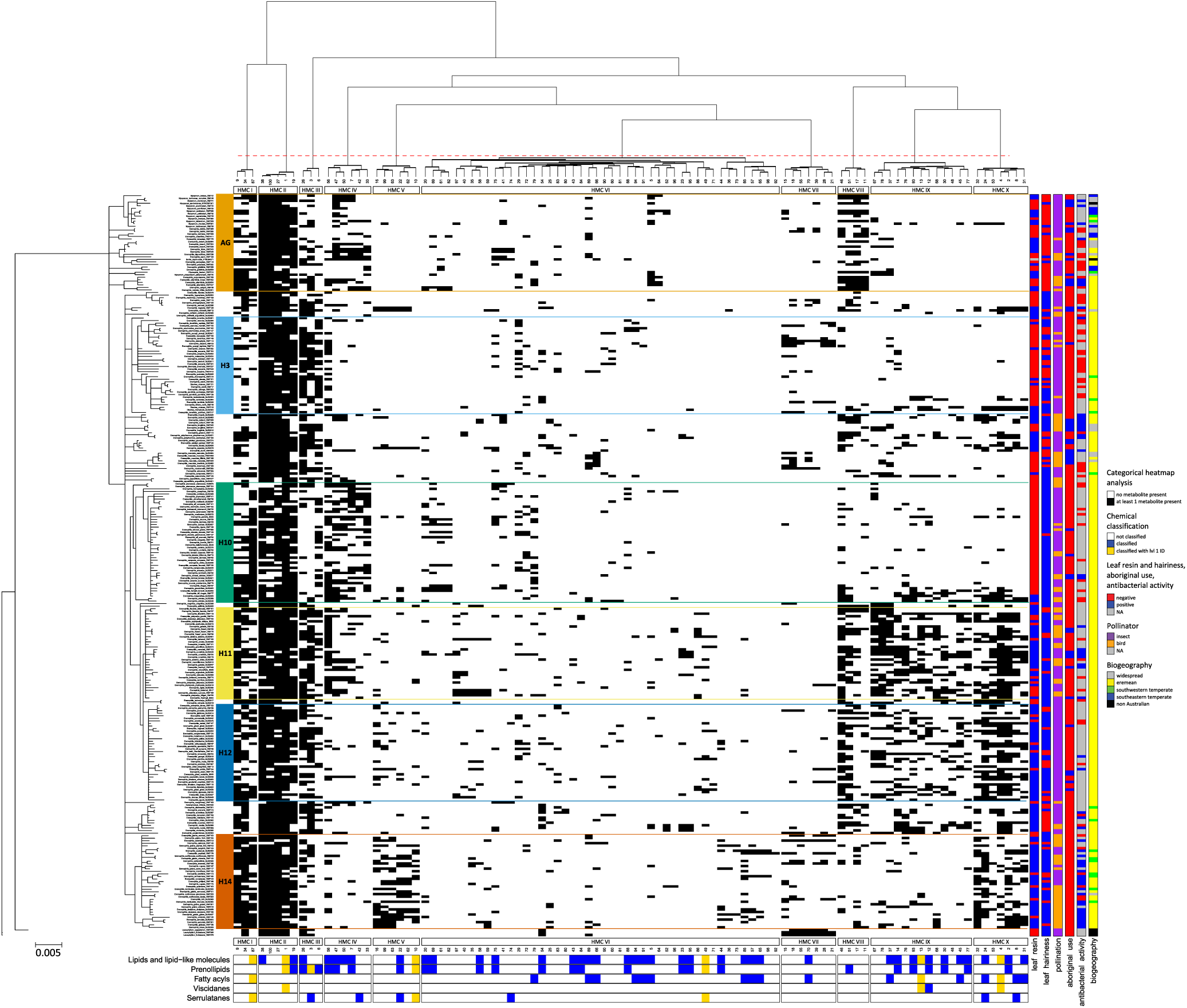
Categorial heatmap analysis displays the chemo-evolutionary framework of Myoporeae. A heatmap analysis was used to evaluate chemical information in an evolutionary context. Information from the 100 largest chemical families was compiled, which derived from the molecular network of Myoporeae. For each specimen in this study, the presence (black) or absence (white) of at least one putative metabolite from a corresponding chemical family is displayed within the heatmap. *LEFT*: The nuclear ribosomal DNA phylogeny on the left side presents the major phylogenetic clades. *TOP*: As depicted on top of the heatmap, the chemical information underwent a hierarchical clustering according to the given phylogeny to reveal chemo-evolutionary patterns among subsets of chemical families, defined as heatmap metabolic clusters (HMC) I – X. *BOTTOM*: Selected information about chemical identity comprise different levels of classification, i.e. superclass (lipids and lipid-like molecules), class (prenol lipids and fatty acyls) as well as annotations on the metabolite level (serrulatane and viscidane-type diterpenoids). Successful dereplication is indicated in blue, while the additional presence of level 1 identification (m/z, retention time and MS^2^ match) in a particular chemical family is highlighted in gold. *RIGHT*: Functional annotations including the presence of leaf resin and hairiness, pollination, antibacterial activity, traditional medicinal usage as well as biogeographical species distribution information are displayed on the right side. A summary of legends are included to the right of the figure.

### Resin chemistry as a key strategy to adapt to the Eremean zone

To relate chemo-evolutionary relationships to specific plant and environmental characteristics, as well as to biological activity, we enhanced the heatmap analysis with metadata on leaf resin and hairiness, pollinator types, geographic distributions, traditional medicinal usage and reported activities against human Gram-positive bacterial pathogens (Table S1). This revealed a strong association between HMC IX and X chemistry and the formation and accumulation of leaf resin in phylogenetic clades H11–H14 (Figure 4). Generalised linear models for each HMC–metadata combination showed that resinous leaves are significantly associated with both HMC IX (manyglm, p = 0.001) and X (manyglm, p = 0.001) (Table S2). The species in these two heatmap metabolic clusters directly correspond to the main members of TMC B with a high frequency of leaf resin relative to TMC A. We hypothesise that HMC IX and X chemistry is a main driver of tanglegram clustering patterns, based on its predominant presence in TMC B (Table S3). PERMANOVA analysis supports a strong overall association of leaf resin on chemical diversity in the dataset, with leaf resin explaining a relatively high degree of variation (8.2%, adonis, F = 25.8, p = 0.001). However, those results could be affected by non-homogeneous dispersion of the data. In fact, specimens without leaf resin are significantly more dispersed (betadisper, p = 0.018), indicating an increased chemical diversity compared to plants with resinous leaves, which are more specialized (Table S4). Notably, no strong associations between leaf chemistry and leaf hairiness nor pollination syndrome are evident (explained variances of 1.4% (adonis, F = 4.16, p = 0.001) and 1.1% (adonis, F = 3.32, p = 0.002), respectively). Univariate testing based on a generalised linear model and subsequent indicator species analysis revealed five chemical families that are significantly associated with bird pollination (Table S5). Those families included a large group of flavonoids (chemical family 9) that comprises the identified compound KU036-6-1. Future work is needed to explore presence of these compounds in other plant organs (i.e. flowers, fruit) and a possible correlation to pollination strategy (i.e. guiding birds through flower colour).

The majority of species in Myoporeae (i.e. phylogenetic clade H) are, with few exceptions including *Bontia*, *Myoporum* and a small number of *Eremophila* species, distributed in arid/semi-arid regions of Australia (Figure S3). The complex evolutionary history and spatially heterogenous nature of Australia’s arid zone is now being recognised [46, 47], although dividing the arid biome into smaller areas for biogeographic analysis remains problematic [48]. In the present study, clear geographic patterns emerge between phylogenetic clades A–H (Figure S3). While phylogenetic clade H is clearly widespread and mainly arid in distribution, the early diverging lineages of the phylogeny (clades A–G) have more coastal/mesic distributions. Within phylogenetic clade H, differences in species distributions between subclades are observed (Figure S4) that can be attributed to the different patterns between the dominant HMC of the heatmap analysis. For example, species in phylogenetic clades H2, H4, H6, H7, H11 and H12 are all distributed in the central arid region of Australia, compared to other subclades that occur in more peripheral southern or eastern arid regions. The central arid phylogenetic clades share the unique HMC IX and X chemistry associated with leaf resin, which is largely absent from other lineages. One exception to this is phylogenetic clade H14 that does show some HMC X chemistry and is also mostly resinous but occurs in the south-west arid region of Western Australia. Unlike all central arid-distributed phylogenetic subclades, H14 also exhibits unique HMC V chemistry not seen anywhere else in the tribe. The exceptional association of HMC V chemistry with clade H14 is considered in greater detail below.

From the observed chemo-evolutionary relationships, we envision a chemistry-based adaptive evolution of species, particularly from the phylogenetic clades H2, H4, H6, H7, H11 and H12, associated with the presence of leaf resin. Further molecular work is needed to resolve relationships between these clades, but they may constitute a single evolutionary lineage adapted for the central/western arid region of Australia. The large number of closely related species in phylogenetic clades H11 and H12 in particular, paired with short branch lengths and a lack of resolution between species relative to other lineages of the phylogeny, suggest that recent and rapid speciation may be responsible for the species diversity. The association between these species and the unique HMC IX and X related serrulatane- and viscidane-type diterpenoids, possibly all localized within the leaf resins [12], may have afforded an essential adaptive advantage in the arms race against arid zone herbivores and pathogens, augmenting the number of specialised diterpenoids formed [49, 50]. A recent chemo-evolutionary adaptation study in the cosmopolitan *Euphorbia* genus advocated that greater herbivory pressure resulted in a highly diversified content of toxic diterpenoids in the Afro-Eurasia geographic region where specialised herbivores co-occur compared to the Americas where specialised herbivores are absent [43]. Evidence suggests that the Australian arid biome is relatively recent in origin and developed within the past 5–10 million years [46, 51]. The contrasting geographic species distributions between the largely non-arid phylogenetic clades A to G at the base of our phylogeny and the mainly arid distribution of phylogenetic clade H (Figure S4) support a hypothesis of a relatively recent diversification of Myoporeae in the arid biome coinciding with the evolution of the unique diterpenoid chemistry represented in HMC IX, X, and V. Following diversification into the arid biome, evolution of a resinous protective layer on the leaf surfaces may offer the additional adaptive advantage of reducing water loss via evapotranspiration and protecting leaves from UV and thermal damage by increased reflection of the solar irradiation [52].

### Chemo-evolutionary relationships of serrulatane and viscidane-type diterpenoids across Myoporeae

We found prenol lipids as the dominant chemical identity within the Myoporeae molecular network, representing 56% of the nodes (1913 nodes) within the 100 largest chemical families. Indeed, the importance of terpenoid-related metabolism in Myoporeae is suggested by the prevalence and patterns of occurrence of chemical families containing serrulatane and viscidane-type diterpenoids in several clades (Figure 4). To further investigate this highly diverse family of diterpenoids [8], we focused on all chemical families within the connected molecular network associated with either serrulatane or viscidane chemistry. Our analysis led to dereplication of 30 distinct chemical families with these diterpenoid scaffolds, involving 1444 nodes (29% of the connected molecular network). Notably, no overlap of these diterpenoid classes within a chemical family was found, despite the structural similarity of the core structures based on a prenyl tail attached to a bicyclic head. Heatmap (Figure S5) and tanglegram analyses (Figure S6) of this chemical space revealed a bloom of metabolite diversity among phylogenetic clades H2, H4, H6, H7 and H11 to H14. This stands in contrast to the less diversified chemistry found in other species of clade H and in the clades A–G. This result supports the observed division of species into TMC A and B, and emphasises their role as important factors in understanding chemo-evolutionary relationships within Myoporeae. The analysis corroborates the co-occurrence of serrulatane and viscidane diterpenoids with the presence of leaf resin, suggesting important physiological roles of the presence of these compounds at the leaf surface. This finding is supported by our recent study, which investigated the underlying biosynthetic pathways in *E. lucida*, *E. drummondii* and *E. denticulata* subsp. *trisulcata* [12], showing that serrulatane and viscidane diterpenoids are concentrated at the resinous leaf surface and synthesized within specialized glandular trichomes embedded within the epidermis. The physiological roles of these diterpenoids are not yet known but could potentially be involved in pathogen defence due to their antibacterial properties [53] or UV protection based either on absorption or resin-mediated increased leaf reflectance [54].

Phylogenetic clade H14 presents a unique chemical signature, which specifically comprises HMC V (Figure 4). Two of the dominant chemical families found in HMC V (10 and 22) were dereplicated as serrulatane-diterpenoid related. Both chemical families show a similar network topology, with a few nodes being found in several species of varying phylogenetic background, besides multiple nodes solely represented by phylogenetic clade H14 (Figure 5). Substructural motif blooms were found within these two chemical families, indicating chemical diversification of serrulatane scaffolds KU006-14 and KU036-12 of chemical families 10 and 22, respectively. Besides some reliable high-level dereplication events, a wide range of metabolite masses, as well as *Eremophila*-specific substructural motifs, indicate a large unknown chemical space of serrulatane chemistry, which has yet to be explored. Most of the species within clade H14 have resinous leaves, as do other species from clades H11 and H12 that also correspond to the diterpenoid enriched TMC B. The majority of species placed in clade H14 are part of TMC B2, with seven species falling in various parts of TMC A as the exceptions. Species within TMC B2 have resinous leaves, while six of the seven species placed in TMC A have non-resinous leaves. The sole exception to this is *E. subfloccosa* subsp. *subfloccosa*, which has resinous leaves and is found in TMC A3. When specifically focusing the metabolic cluster analysis on all chemical families within the connected molecular network associated with serrulatane and viscidane chemistry, *E. subfloccosa* subsp. *subfloccosa* does in fact localize within the resinous members of phylogenetic clade H14 (Figure S6). Furthermore, none of the non-resinous species found in clade H14 share metabolites from the serrulatane-related chemical families in HMC V (10 and 22), underlining the relationship between serrulatane formation and accumulation of leaf resins.

**Figure 5.**
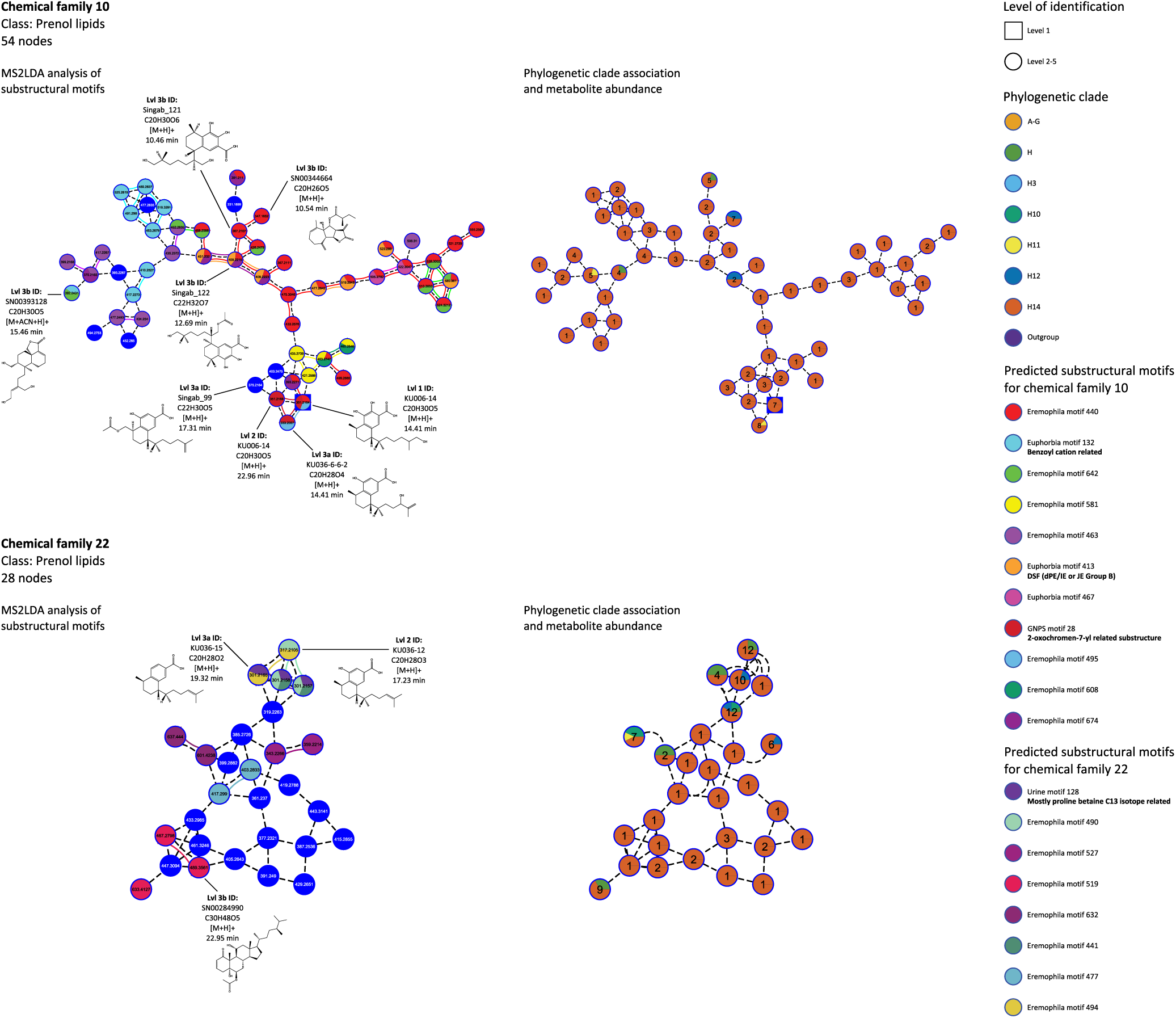
In depth analysis of selected serrulatane diterpenoid-related chemical families. Detailed analysis of serrulatane diterpenoid-related chemical families 10 and 22. On the left, chemical families are displayed with detected metabolite mass (Da) information for each node as well as individual dereplication events, which are highlighted with selected spectrometric and structural information. In addition, MS2LDA-based substructure analysis is shown by color coded pie charts on each node, which points out areas of shared spectral motifs. On the right side, investigated chemical families harboring information about phylogenetic clade association, as displayed for every node with colored pie charts. Also the total amount of occurrences of each putative metabolite in the dataset is presented for every node.

Phylogenetic clade H14 stands out as an interesting group within tribe Myoporeae because it harbours morphologically highly diverse *Eremophila* species, a unique chemical signature and strongly associated antibacterial properties. Notably, there are no published instances of traditional medicinal use for these species. Both phylogenetic and metabolic cluster analyses support the unexpected placement of morphologically diverse species from three different sections of *Eremophila* (*sensu* [3]) in phylogenetic clade H14. Two of these sections (*Stenochilus* and *Virides*) comprise morphologically similar species, while the third section (*Australophilae*) contains species that vary considerably from the other two. Morphologically, species in clade H14 have not previously been considered as closely related, however strong phylogenetic support for this lineage, paired with the unique metabolome identified in the current study, has allowed these relationships to be revealed. This case underlines the utility of combining molecular phylogenetics and metabolite profiling: the observed distribution of particular chemical families supported unexpected phylogenetic relationships and furthermore, raises questions about the potential interplay between specialised metabolism and species diversification for future investigation [55].

### The molecular networking approach facilitates discovery of novel specialised metabolites

Our approach guides drug isolation efforts by narrowing the large number of metabolites down to chemical families and predicting chemical classification. This information can be used to target unknown chemical analogues within chemical families with known bioactivities of interest. In addition, functional annotation with the results of antimicrobial assays of crude plant extracts can guide the selection process to uncover novel chemical families not previously considered. A univariate test based on a generalised linear model including tested specimens, revealed chemical families significantly associated with antibacterial activity that are solely from the terpenoid enriched HMC V and X. A subsequent indicator species analysis further associates these families with positive activity, highlighting them as interesting targets for antibacterial drug discovery (Table S5). When focusing on bioactive serrulatane and viscidane-type diterpenoids, many of the chemical families are not well structurally dereplicated and are thus excellent targets for drug discovery. In fact, 20 out of 30 chemical families that are part of this particular chemical space contain no level 1 identified metabolite, providing a source of unknown serrulatane or viscidane analogues (Figure S5). To maximize investigated chemical diversity while minimizing drug isolation efforts, we can use the heatmap analysis to identify ideal species for sampling. Using this approach, five plant species (*E. galeata*, *E. enata*, *E. spectabilis* subsp. *brevis*, *E. margarethae* and *E. glabra* subsp. Paynes Find) out of a total 206 species were shown to cover up to 18% of the observed chemical space of viscidane and serrulatane-type diterpenoids, accounting for 1444 putative metabolites in total. Other well dereplicated chemical families have been identified and should be explored to further enrich our knowledge of the highly diverse diterpenoid structures present. For example, chemical family 10 was found to contain the highly oxygenated serrulatane KU006-14 besides other identified analogues (Figure 5). Serrulatane KU006-14 was only found in seven of the 23 total species in phylogenetic clade H14, and thus represents a specific branch of the serrulatane-related chemical space. KU006-14 is identical to 7,8,16-trihydroxyserrulat-19-oic acid which we previously isolated from *E. drummondii* and carries a hydroxylation at C-16 as a desired target for substitutions to obtain derivatives with increased Type 2 anti-diabetic activity [11]. Thus the C-16 positioned 3-methylbutanoyl ester drastically decreased the IC_50_ of PTP1B inhibition from 1260 ± 560 µM (KU006-14) to 3.44 ± 0.88 µM [11]. A metabolite with the corresponding mass ([M+H]+ of 435.2739) to this chemical analogue can be found in chemical family 10 in close proximity to KU006-14. Further exploration of chemical family 10-associated diterpenoids, and subsequent testing for their anti-diabetic activities, may reveal the presence of even better drug candidates in other species of phylogenetic clade H14. Additional synthetic functionalization may further extend the potential biological activity of KU006-14-related metabolites. This has been documented for the close analogue 3,7,8-trihydroxyserrulat-14-en-19-oic acid [56]. This compound did not show any antimalarial activity, whereas a number of derived amides did.

Our results show how our molecular networking pipeline can aid in the identification of both plant species and chemical families with potential as sources of new bioactive molecules. Chemical families with low levels of metabolite identification based on *in silico* spectral support are easily determined, and represent a prime source to unfold the rich chemical ”dark matter” in Myoporeae [37]. A single example is chemical family 5, which includes 110 nodes and was dereplicated as fatty acid related. Notably, this prediction is solely based on level 3a identifications, and therefore warrants research to identify the structures of some of the compounds in the molecular network. The median mass of 332.2732 Da and retention time of 17.8 min of this chemical family is comparable with known diterpenoids in this study and further suggests an interesting source for potential drug candidates. Additionally, this chemistry is mainly localized to species of phylogenetic clades A–G, indicating its role in specialized metabolism. Notably, the resinous *Glycocystis beckeri* and *Myoporum bateae* contribute significantly to the chemical space in chemical family 5 and thus represent suitable targets for investigation. Additionally, these two species also harbor flavone related metabolites from chemical family 11, which accumulate mostly in clades A-G.

Australian Aboriginal Peoples use herbal extracts from some *Eremophila* species as integrated components of their traditional medicine [1–3]. These herbal extracts contain a multiplicity of bioactive natural products that individually or in combination could provide the perceived curing effects. In an attempt to begin to associate the presence of specific bioactive natural products or combinations thereof to the beneficial effects, species with documented Aboriginal uses were highlighted in our current study. To underline this approach, measured anti-bacterial effects of herbal extracts against human pathogenic bacterial species and of individual isolated natural products were also integrated into this investigation (Table S1). Traditional medicinal uses of herbal extracts were widespread throughout Myoporeae. Interestingly, no use of *Eremophila* species from phylogenetic clade H14 was found in the literature. This contrasts with the number of reports documenting a high abundance of herbal extracts and isolated diterpenoids possessing anti-bacterial activity within this subclade [18, 57]. This could indicate that the diversity of serrulatanes in clade H14 somehow is associated with negative effects in traditional medicine. However, the use of plants within Aboriginal traditional medicine is likely to be underreported in the literature. It is recognised that much traditional knowledge was lost with the break-up of traditional societies and the denial of access to land and natural resources for Aboriginal Peoples following the British colonisation of Australia [58, 59]. Further, some traditional custodians may have chosen not to have this information recorded in the public domain. Therefore, the absence of recorded traditional use of *Eremophila* species in clade H14 does not mean that some of these species were not used by a specific group at some point in time. Nevertheless, from the records it is clear that *E. alternifolia* had and continues to have a prominent medicinal role [1, 8, 58], and is recognised in our current study for its rich flavone-related chemistry. A second prominent species is *E. cuneifolia* [1, 8] with a distinct elevated compound diversity in terpenoid-related chemical families 23 and 95, which are mostly absent in other species. Thus, successful functional annotations to a specific molecular network could enable assignment of metabolites present to specific biological activities, morphological traits and geographical distributions and enable targeted drug discovery approaches [39, 60].

## Conclusion

The present study highlights the power of combining large-scale molecular networking and phylogenetic analyses to investigate chemo-evolutionary diversification that involves specialized diterpenoid chemistry associated with resin development in Myoporeae. By integrating functional annotations in our analysis, we hypothesize that this specialized chemistry may complement the transpiration barrier function of the resin with pathogen and herbivore defence as well as UV protection. To shed further light on the broader area of arid zone ecosystem evolution, ecological and phylogenetic information for associated insects could be integrated into the chemo-evolutionary framework developed here for Myoporeae. A number of studies have shown several insect groups that occur almost exclusively with *Eremophila* [61, 62]. Further research in this area may help us to understand the links between chemistry and species radiations (both plant and insect) as well as regional and fine scale patterns of diversification.

Despite great progress in illuminating the chemical space of Myoporeae, there are still chemical families of unknown character waiting to be elucidated. Our study shows the state-of-the-art computational tools that can be used to target drug discovery approaches by giving an idea about the identity of the observed chemical space. Additionally, the chemical component of the present study focused solely on leaf chemistry. However, ongoing research in *Eremophila* and other plant groups has revealed that other plant organs may also be rich sources of distinct bioactive metabolites. An example is the antimalarial microthecaline A, a novel quinoline–serrulatane alkaloid, which was found in roots of *E. microtheca* and *M. insulare* [14, 21].

By establishing a chemo-evolutionary framework and enriching it with functional annotations, we present a systematic outline of chemistry and evolution in Myoporeae and lay foundations for further interdisciplinary research. We encourage the use and further extension of this dataset through functional annotations to explore chemo-evolutionary relationships and to select compelling chemical families for natural product isolation to describe the unknown aspects of Myoporeae chemistry.

## Acknowledgements

This work was supported by the VILLUM Center for Plant Plasticity (VKR023054) (BLM); the European Research Council Advanced Grant (ERC-2012-ADG 20120314) (BLM), the Lundbeck Foundation (R223-2016-85, ”Brewing diterpenoids”), the Cybec Foundation (Jim Ross Scholarship), and the Novo Nordisk Foundation Interdisciplinary Synergy (NNF 16OC0021616, “Desert-loving therapeutics”) and Distinguished Investigator 2019 (NNF 0054563, “The Black Holes in the Plant Universe”) programs (BLM). We would like to thank Tanja Schuster for specimen collections, and Bob Chinnock and Ron and Claire Dadd for access to their private garden collections. Australian National Botanic Garden, Canberra, the Australian Arid Lands Botanic Garden, Port Augusta, the National Tropical Botanic Garden, Kauai for collection of garden specimens. Elizabeth H. J. Neilson (UCPH, Denmark) for initiating the chemical profiling work of *Eremophila* species. Madeleine Ernst (SSI, Denmark) for the advice regarding molecular networking tools. Yong Zhao (UCPH, Denmark) and Laura Mikél Mc Nair (UCPH, Denmark) for providing reference compounds to establish the in house spectral library. Dominik Merges (BiK-F, Frankfurt) and the PLEN R-club for additional advice regarding multivariate statistics.

## Author Contributions

OG, RMF and BLM designed the study. RMF, MJB and BJB collected, identified, and sampled *Eremophila* and related species from field and cultivated collections from across Australia. RMF and MJB performed taxonomic description and phylogenetic analyses of these specimens. OG and AMH prepared the plant extracts and performed the LC-qToF-HRMS analysis. OG conducted the mass spectral data processing and quality check as well as subsequent molecular networking analysis. All reference compounds derived from *Eremophila* species were isolated, structurally characterized and provided by DS. OG established the in house reference compound and *in silico* spectral databases and performed dereplication of the molecular network including the MS2LDA substructural analysis. OG established the computational pipeline to establish and analyse the chemo-evolutionary framework in Myoporeae and wrote all scripts used to conduct data processing as well as cluster, tanglegram, heatmap and statistical analysis. Different authors provided information about functional annotations, leaf resin, hairiness and pollination (RMF and BJB), antibacterial activity and traditional usage (RMF, SJS and CPN) and biogeography (RMF and DJM). Species distribution mapping was completed by RMF. OG, RMF and BLM wrote the manuscript. All authors discussed the results and contributed with comments during the writing process.

## Competing Interests statement

The authors declare that there are no competing interests.

## Experimental section

### Collection of plant material

Leaf tissue and herbarium voucher specimens were collected from field and cultivated collections from across Australia. A cultivated specimen of *Bontia daphnoides* was sent from Fairchild Tropical Botanic Garden, Florida. Collection details, herbarium voucher numbers and GenBank accession numbers are provided in Table S1. Leaf material was picked fresh and stored immediately in silica beads. The same silica dried leaf material was used for DNA and phytochemical extractions.

### Phylogenetic analysis

Total genomic DNA was isolated using a modified cetyltrimethylammonium bromide (CTAB) protocol [63]. Library preparation was completed ‘in-house’, using a version of the sample preparation protocol outlined in [64]. Samples were sequenced using both Illumina HiSeq2000 (2 x 125 bp) and Illumina NextSeq500 (2 x 150 bp) sequencing platforms, based at AgriBio, Centre for AgriBioscience (La Trobe University) and the Walter and Eliza Hall Institute (WEHI) in Melbourne, Australia. Raw read data of total genomic DNA was *de-novo* assembled using CLC Genomics Workbench version 8.5.1 (https://www.qiagenbioinformatics.com/) with default settings. Resulting contigs were imported into Geneious version 9.1.8 (http://www.geneious.com [65], where the nuclear ribosomal cistron was constructed for each sample, initially using the partial ITS/ETS sequence available for *Eremophila macdonnellii* on GenBank (DQ444239) as a reference. Contigs were mapped to *E. macdonnellii* using custom sensitivity settings; gaps allowed set to a maximum of 20% per read, maximum gap size of 3000 and 2–3 iterations. Total raw reads were then mapped back to the consensus sequence and a final consensus sequence extracted using a consensus threshold of 75%, to confirm sequence accuracy. Nuclear ribosomal cistron length varied between samples, from 5976–7863 base pairs. Differences in sequence lengths were largely due to variation in DNA read coverage of the external transcribed spacer (ETS)/non-transcribed spacer (NTS) regions. All samples contained complete coverage of 18S, ITS1, 5.8S, ITS2 and 26S nrDNA regions. Sequences were aligned in Geneious using the MAFFT pairwise alignment plug-in (MAFFT version 7.222; [66]) with default settings. The alignment was assessed by eye and any adjustments were made manually. After exclusion of poorly aligned sequence ends, the 291 nuclear ribosomal sequences resulted in an aligned matrix of 6568 bp. This alignment is available in TreeBase (study accession number 26197). Phylogenetic analysis was completed using MrBayes version 2.3 [67]. The alignment was partitioned into six character sets representing coding ribosomal DNA sequence (18S, 5.8S and 28S genes) and intergenic spacers (ITS1, ITS2 and ETS + non-transcribed spacer). Models of evolution for each partition were estimated using the Bayesian information criterion (BIC) in IQ-TREE version 1.6.12 [68]. Where exact models were not available in MrBayes equivalent models were selected (18S: GTR + I + G, 5.8S: K2P + I, 26S: GTR + I + G, ITS1: SYM + G, ITS2: SYM +I + G, non-transcribed spacer: GTR+ I + G). The BI analysis was run for 15 million generations with unlinked partitions and a tree sampling frequency of 1000 to estimate posterior probabilities. The average standard deviation of split frequencies reached a value less than 0.01 during this analysis, with convergence of MCMC chains checked in Tracer version 1.6 [69]. Twenty five percent of the trees were discarded as burn-in, a consensus tree was generated, and Bayesian posterior probabilities estimated for nodes from the consensus tree (Figure S7).

### Metabolite analysis

Ground and dried leaf tissues were extracted with 50% acetonitrile (supplemented with 2ppm forskolin), while shaking incubation at 25 °C for 2 h. Acetonitrile extracts were filtered using a 0.22 µm 96-well filter plate (Merck Millipore, Darmstadt, Germany) and analysed using an Ultimate 3000 UHPLC+ Focused system (Dionex Corporation, Sunnyvale, CA) coupled to a Bruker Compact ESI-QTOF-MS (Bruker) system. Samples were separated on a Kinetex XB-C18 column (100 × 2.1 mm ID, 1.7 µm particle size, 100 Å pore size; Phenomenex Inc., Torrance, CA) maintained at 40 °C with a flow rate of 0.3 mL min-1 and mobile phase consisting of 0.05% (v/v) formic acid in water (solvent A) and 0.05% (v/v) formic acid in acetonitrile (solvent B). The LC method was as follows: 0-1 min, 10% B; 1-23 min, linear increase from 10 to 100% B; 23-25 min, 100% B; 25 to 25.5 min, 100-20%; 25.5-30.5 min, linear decrease to 10% B. Mass spectra were acquired in positive ion mode over a scan range of m/z 50-1200 with the following ESI and MS settings: capillary voltage, 4000 V; end plate offset, 500 V; dry gas temperature, 220 °C; dry gas flow of 8 L min^-1^; nebulizer pressure, 2 bar; in source CID energy, 0 eV; hexapole RF, 50 Vpp; quadrupole ion energy, 4 eV; collision cell energy, 7 eV. Samples were further subjected to untargeted LC-MS/MS using a collision cell energy of 27 eV. Quality control samples were prepared from a pool of different extract samples and run after a sequence of 22 samples as well as before and after the total run. Raw chromatogram data was calibrated using an internal sodium formate standard and subsequently exported as .mzML format using DataAnalysis 4.1 (Bruker). Further processing of the raw chromatogram data was conducted using MZmine2 (v.2.5.3) [70]. At first, a signal intensity noise cutoff of 1000 and 100 was applied to MS^1^ and MS^2^ data, respectively. In addition, only scans between 0.5 and 24 min retention time were considered. Chromatograms have been built using the ‘ADAP chromatogram builder’ with m/z tolerance of 0.003 Da (5 ppm) and a signal intensity threshold of 20000. The generated extracted ion chromatograms were deconvoluted into individual peaks using the ‘local minimum search’ module. After isotopes were removed, the feature list was aligned using the ‘joint aligner’ tool using m/z tolerance of 0.006 Da (10 ppm) and retention time tolerance of 0.1 min. Subsequently, individual metabolites have been identified by comparing it to MS^2^ data of an in house spectral database (level 1 identification includes m/z, RT and MS^2^ similarity). This library includes 76 reference compounds commercially sourced as well as isolated from different *Eremophila* species during prior studies ([71], Table S6), which were analysed with the same MS system as described above. Missing data points have been added using the ‘gap filler’ tool and afterwards intensities below 1000 set to 0, by applying a peak filter. The resulting peak list was further used to undergo a quality control that was conducted by a principal component analysis comprising all species and quality control samples on top of the gap-filled (but not peak filtered) peak list. In this projection, all quality control samples are found in close proximity, indicating reliable data acquisition over time (Figure S8). Finally, background features that were shared between the samples and quality controls were removed manually, which includes the internal standard forskolin as well (data kept for normalization). The generated peak list was exported twice, as csv-file (containing peak signal intensity) for further processing in R script and via the ‘GNPS-FBMN export’ module for subsequent upload to the GNPS server. A detailed description of the MZmine2 processing parameters used can be found in Table S7. A molecular network was created with the Feature-Based Molecular Networking workflow [45] on GNPS (https://gnps.ucsd.edu) [38]. The data was filtered by removing all MS^2^ fragment ions within +/- 17 Da of the precursor m/z. MS^2^ spectra were window filtered by choosing only the top 6 fragment ions in the +/- 50 Da window throughout the spectrum. The precursor ion mass tolerance was set to 0.02 Da and the MS^2^ fragment ion tolerance to 0.02 Da. A molecular network was then built where edges were filtered to have a cosine score above 0.8 and more than 8 matched peaks. Further, edges between two nodes were kept in the network if and only if each of the nodes appeared in each others respective top 5 most similar nodes. Finally, the maximum size of a chemical family was set to 0 (no size limitation), and the lowest scoring edges were removed from chemical families until the chemical family size was below this threshold. The spectra in the network were then searched against GNPS spectral libraries [38, 72] as well as an in house library including 76 reference spectra (level 2 identification includes MS^2^ similarity). The library spectra were filtered in the same manner as the input data. All matches kept between network spectra and library spectra were required to have a score above 0.7 and at least 6 matched peaks. The molecular networks were visualized using Cytoscape software (v3.7.2) [73]. To predict a consensus of chemical classification for individual chemical families, the network annotation propagation tool was applied to the generated molecular network. Thereby, multiple public *in silico* spectral databases were used, i.e. GNPS, SUPNAT, NPAtlas, CHEBI and DRUGBANK. Additionally, an *Eremophila* specific in house *in silico* fragmentation database was used that contains 293 metabolite structures that have been characterized in *Eremophila* species during prior studies [8, 71]. The *in silico* fragmentation based dereplication results were categorized by their reliability, with SMILES from Fusion, Consensus and MetFrag algorithm corresponding to level 3a, 3b and 3c identification, respectively. Additionally, insight into substructural information was achieved using the MS2LDA motif search via the GNPS server. MS2LDA allows annotation of smaller substructures shared by metabolites of a chemical family within a molecular network [74]. Eventually, the results from the different molecular network based analyses were joined via the MolNetEnhancer (v15) tool [75]. A detailed description of the molecular networking pipeline presented in this study can be found in Table S8. The MZmine2 pre-processed chromatogram data was further processed for downstream analysis using R script (v3.6.1). Thereby, the data was normalized to the median of the internal standard and sample weight. Subsequently, a signal intensity cutoff of 10000 was applied and resulting features that had no occurrence anymore were removed, resulting in a normalized dataset of 10696 features. Eventually, every intensity above 0 was assigned the value ‘1’ to create a binary matrix. To add the information gained from the molecular network analysis, the 100 largest chemical families (9 – 810 nodes per chemical family, Table S9) have been isolated and combined with the processed chromatogram data to generate a new data matrix. In this way, a ‘continous’ dataset that contains the number of features that are shared of a particular chemical family for each species was generated. Additionally, a ‘categorical’ dataset was generated that contains solely the value ‘1’, for a species that shares at least one node of a chemical family, and ‘0’ for not having a single node (Figure 4). Based on the categorical dataset, a distance matrix was generated using Jaccard distance, which was subjected to a hierarchical cluster analysis using the ‘ward.d2’ agglomeration method. Using the ‘ape’ package (v5.3), this analysis was converted to a dendrogram, which was then compared to the dendrogram derived from the molecular phylogeny via a tanglegram analysis (‘dendextend’ package v1.12.0). The resulting tanglegram was untangled using the ‘step2side’ algorithm and the rearranged molecular phylogeny used for further heatmap analyses. Using the R package ‘ComplexHeatmap’, a heatmap was generated based on the ’categorical’ dataset using Jaccard distance and ‘ward.d2’ agglomeration method (km=10000), which clusters the presence/absence of chemical families according to phylogenetic similarity, while the order of specimen in the phylogeny is fixed. A ’continous’ heatmap analysis is based on the cluster generated by the ’categorical’ analysis and highlights the total number of putative metabolites from a chemical family for each specimen (Figure S9). Combined detailed information of tanglegram and heatmap analysis are found in Table S10. Additional information about chemical classification (Table S11, Table S12) and leaf morphology (resin and hairiness), environmental factors (pollination and geographical distribution) and medicinal properties (traditional medicinal uses and antibacterial studies) were aligned to the heatmap analyses. Another heatmap analysis focused solely on serrulatane and viscidane diterpenoid related chemical families. In that regard, all chemical families (down to 2 nodes) were selected manually from the global molecular network that contain at least one corresponding dereplication hit of level 1-3c (Table S13, Table S14). These 30 chemical families were subjected to a tanglegram analysis as described above (Figure S6) as well as a heatmap analysis that clustered the ’continous’ chemical information using Euclidean distance and ‘complete’ agglomeration method (Figure S5). Another tanglegram was conducted based on the binary dataset including all metabolites after normalization (Figure S10).

Raw spectral data available at: (MassIVE link will be provided upon journal publication).

The molecular networking job can be publicly accessed at https://gnps.ucsd.edu/ProteoSAFe/status.jsp?task=f74d3374a7ba43eb8d58e5d2a312bf33.

The network annotation propagation job can be publicly accessed at https://proteomics2.ucsd.edu/ProteoSAFe/status.jsp?task=5980fea229b9427bb610a8e7d6c25827.

The MS2LDA job can be publicly accessed at https://gnps.ucsd.edu/ProteoSAFe/status.jsp?task=8082c8133d1843c39af42164157ab043.

### Statistical analysis

To test for overall significant differences of functional annotations among the investigated specimens, we used the non-parametric permutational multivariate statistical tests: permutational multivariate analysis of variance (PERMANOVA) [76]. This method use a dissimilarity metric to calculate for differences between annotation object classes. For PERMANOVA, the null hypothesis is that the metric centroid does not differ between groups. PERMANOVA was calculated with the adonis2() function in the vegan package (http://cran.r-project.org/package=vegan) [77]. This analysis was based on the categorical dataset, for which dissimilarity metric was generated by the vegdist() function using Jaccard distance. PERMANOVA is sensitive to differences in data dispersion within groups and may therefore confuse within-group variation with among-group variation [78]. To test if groups differed in their dispersion, we used the betadisper() function with the null hypothesis that the average within-group dispersion is the same in all groups. In each test, the number of permutations was set to 999. For all analyses, we considered a p-value threshold of 0.05 significant. We conducted these tests on the full dataset (100 chemical families), to test the significance of functional annotations of leaf resin, leaf hairiness, antibacterial activity, pollination, biogeography and general clades (“O”: outgroup, “AG”: clade A to G, “H3”: clade H3, “H10”: clade H10, “H11”: clade H11, “H12”: clade H12, “H14”: clade H14 and “H”: remaining clade H associated taxa), whereby specimens with missing values were excluded. Since records for aboriginal usage are scarce, we excluded it from all statistical analyses. We used generalised linear models to test for significant differences of functional annotations among the investigated specimens within a HMC background, [79]. Models were generated for each HMC–annotation combination using the clogloc model, while excluding specimens with missing functional annotation. Each test was conducted using the anova.manyglm() function from the mvabund package [80]. In each test, the number of bootstrap iterations was set to 999 and used the montecarlo resampling method. For all analyses, we considered a p-value threshold of 0.05 significant. To access individual chemical families as overall significant drivers for a functional annotation, a subsequent univariate test was conducted for the generated generalised linear models to receive individual p-values. An additional indicator species analysis using the indval() function (labdsv package [81]) was applied to indicate the direction of annotation (level) for each chemical family (Table S5).

## Supplementary figures and tables

### Supplementary tables

**Table S1.** Specimen details for all samples included in this study. Specimen information include collecting number, species name (authority), *Eremophila* section, collection location, herbarium voucher number and GenBank accession number. Collecting numbers followed by * denote specimens grown in cultivation, where provenance information was known this was recorded in latitude/longitude. Collection initials refer to: RMF (Rachael Fowler), MJB (Michael Bayly), BB (Bevan Buirchell), TMS (Tanja Schuster), FTBG (Fairchild Tropical Botanic Garden), NTBG (National Tropical Botanical Garden). Published records of traditional medicinal use for each species is recorded where available [1, 18, 22, 58, 82–108]. Evidence of antimicrobial activity against human bacterial pathogens (gram positive bacteria) for each species is recorded where available [7, 9, 18, 22, 23, 25–27, 57, 97, 109–112].

| Collection number | Taxon name and authority | *Eremophila* section (*sensu* Chinnock 2007) | Latitude | Longitude | Herbarium voucher number | GenBank accession number | Published traditional medicinal use for this species | Evidence available of antimicrobial activity for this species |
| --- | --- | --- | --- | --- | --- | --- | --- | --- |
| RMF211 | *Eremophila compressa* Chinnock | *Platycarpus* | -32.90429 | 121.60685 | MELUD119198a | MN411553 |  |  |
| RMF36* | *Eremophila saligna* (S. Moore) C. Gardner | *Platycarpus* | -31.2314 | 121.4603 | - | MN486179 |  |  |
| MJB2384* | *Eremophila deserti* (Cunn. ex Benth.) | *Pholidiopsis* | -30.82 | 117.48 | MELUD119043a | MN411456 |  |  |
| RMF204 | *Eremophila deserti* (Cunn. ex Benth.) | *Pholidiopsis* | -32.94995 | 123.20631 | MELUD119194a | MN411550 |  |  |
| RMF228 | *Eremophila deserti* (Cunn. ex Benth.) | *Pholidiopsis* | -34.29655 | 142.2312 | MELUD119206a | MN411558 |  |  |
| RMF243 | *Eremophila fallax* Chinnock | *Pholidiopsis* | -31.27039 | 138.87086 | MELUD119213a | MN411564 |  |  |
| RMF246 | *Eremophila fallax* Chinnock | *Pholidiopsis* | -31.27932 | 138.57864 | - | MN486169 |  |  |
| MJB2380* | *Eremophila micrantha* Chinnock | *Crustaceae* | -25.48 | 119.65 | MELUD119040a | MN411453 |  |  |
| RMF212 | *Eremophila succinea* Chinnock | *Crustaceae* | -32.82656 | 121.19814 | MELUD119199a | MN411554 |  |  |
| RMF192 | *Eremophila dempsteri* F. Muell. | *Crustaceae* | -32.15891 | 120.58533 | MELUD119186a | MN411545 |  |  |
| MJB2362* | *Eremophila ciliata* Chinnock | *Crustaceae* | -33.41 | 123.4 | MELUD119021a | MN411437 |  |  |
| RMF201 | *Eremophila dichroantha* Diels | *Crustaceae* | -32.06511 | 122.37669 | MELUD119191a | MN411548 |  |  |
| RMF144* | *Eremophila interstans* subsp. *virgata* (W.V. Fitzg.) Chinnock | *Crustaceae* | -30.99 | 121.13 | MELUD119150a | MN411382 |  |  |
| RMF234 | *Eremophila sturtii* R. Br. | *Crustaceae* | -34.14759 | 141.30539 | MELUD119210a | MN411561 | [22, 58, 82, 83] | [22, 97] |
| RMF12* | *Eremophila mitchellii* Benth. | *Crustaceae* | -32.75 | 145.5833 | CBG7910121 | MN411422 | [1] |  |
| RMF145* | *Eremophila paisleyi* subsp. *paisleyi* F. Muell. | *Crustaceae* | -31.04 | 123.56 | MELUD119151a | MN411383 | [82, 84] |  |
| RMF274* | *Eremophila paisleyi* subsp. *glandulosa* Chinnock | *Crustaceae* | - | - | MELUD119230a | MN411406 |  |  |
| MJB2428 | *Eremophila falcata* Chinnock | *Crustaceae* | -26.56599 | 120.00422 | MELUD119081a | MN411470 |  |  |
| MJB2379* | *Eremophila caperata* Chinnock | *Crustaceae* | -31.32 | 119.13 | MELUD119039a | MN411452 |  |  |
| RMF289 | *Eremophila debilis* (Andrews) | *Chamaepogonia* | -36.1082 | 145.8249 | MELUD119238a | MN411583 | [85] |  |
| RMF290 | *Eremophila debilis* (Andrews) | *Chamaepogonia* | -36.1082 | 145.8249 | MELUD119239a | MN411584 | [85] |  |
| RMF218 | *Eremophila chamaephila* Diels | *Australophilae* | -33.5168 | 120.58369 | MELUD119203a | MN411556 |  |  |
| MJB2393* | *Eremophila perglandulosa* Chinnock | *Australophilae* | -30.79 | 123.46 | MELUD119049a | MN411462 |  |  |
| RMF17* | *Eremophila weldii* F. Muell. | *Australophilae* | -31.6667 | 128.9167 | CBG59633 | MN411418 |  |  |
| RMF254 | *Eremophila weldii* F. Muell. | *Australophilae* | -33.52283 | 135.72063 | MELUD119221a | MN411570 |  |  |
| RMF126* | *Eremophila parvifolia* subsp. *parvifolia* J. Black | *Australophilae* | - | - | MELUD119135a | MN411376 |  |  |
| RMF200 | *Eremophila parvifolia* subsp. *auricampa* Chinnock | *Australophilae* | -32.1581 | 121.76448 | MELUD119190a | MN411396 |  |  |
| RMF203 | *Eremophila oblonga* Chinnock | *Australophilae* | -32.0843 | 123.0882 | MELUD119193a | MN411549 |  |  |
| MJB2461 | *Eremophila ionantha* Diels | *Australophilae* | -30.95319 | 121.1434 | MELUD119107a | MN411501 |  | [18] |
| RMF252 | *Eremophila praecox* Chinnock | *Australophilae* | -33.12447 | 136.24911 | MELUD119219a | MN411569 |  |  |
| MJB2458 | *Eremophila pustulata* S. Moore | *Australophilae* | -30.17 | 120.60627 | MELUD119105a | MN411499 |  |  |
| MJB2392* | *Eremophila homoplastica* (S. Moore) C. Gardner | *Australophilae* | -28.18 | 119.29 | MELUD119048a | MN411461 |  |  |
| MJB2364* | *Eremophila verticillata* Chinnock | *Australophilae* | -33.1 | 119.02 | MELUD119027a | MN411443 |  |  |
| MJB2358* | *Eremophila ternifolia* Chinnock | *Australophilae* | -30.89 | 116.72 | MELUD119018a | MN411434 |  |  |
| MJB2394* | *Eremophila veronica* (S. Moore) C. Gardner | *Australophilae* | -30.99 | 121.13 | MELUD119050a | MN411463 |  |  |
| RMF132* | *Eremophila caerulea* subsp. *merralli* (F. Muell. ex Ewart, Jean White & B. Wood) | *Australophilae* | -31.88 | 118.15 | MELUD119140a | MN411379 |  |  |
| RMF191 | *Eremophila clavata* Chinnock | *Australophilae* | -32.15881 | 120.58519 | MELUD119185a | MN411544 |  |  |
| RMF217 | *Eremophila densifolia* subsp. *pubiflora* Chinnock | *Australophilae* | -32.62364 | 119.7429 | MELUD119202a | MN411399 |  |  |
| RMF263* | *Eremophila densifolia* subsp. *capitata* Chinnock | *Australophilae* | - | - | MELUD119226a | MN411404 |  |  |
| RMF133* | *Eremophila microtheca* (F. Muell. ex Benth.) | *Australophilae* | - | - | MELUD119141a | MN411520 |  | [18, 23, 57] |
| RMF186 | *Eremophila phillipsii* F. Muell. | *Australophilae* | -32.54483 | 118.60204 | MELUD119180a | MN411540 |  | [18] |
| RMF143* | *Eremophila complanata* Chinnock | *Australophilae* | -30.9 | 118.66 | MELUD119149a | MN411526 |  | [18, 57] |
| RMF125* | *Eremophila adenotricha* (F. Muell. ex Benth.) | *Australophilae* | -31.93 | 119.1 | MELUD119134a | MN411517 |  |  |
| RMF172* | *Eremophila rugosa* Chinnock | *Australophilae* | -30.98 | 121.14 | MELUD119170a | MN411534 |  | [18], Semple *et al.* (unpub.) |
| RMF187 | *Eremophila rugosa* Chinnock | *Australophilae* | -32.41673 | 119.71435 | MELUD119181a | MN411541 |  |  |
| RMF176* | *Eremophila papillata* Chinnock | *Australophilae* | -30.82 | 116.68 | MELUD119174a | MN411538 |  |  |
| MJB2363* | *Eremophila scaberula* W.V. Fitzg. | *Australophilae* | -30.63 | 116 | MELUD119026a | MN411442 |  | [57] |
| RMF123* | *Eremophila sargentii* (S. Moore) Chinnock | *Australophilae* | -30.82 | 116.68 | MELUD119132a | MN411515 |  |  |
| RMF173* | *Eremophila brevifolia* (Bartling) | *Australophilae* | -28.399 | 115.01 | MELUD119171a | MN411535 |  |  |
| RMF185 | *Eremophila lehmanniana* (Sonder ex Lehm.) Chinnock | *Australophilae* | -32.328 | 117.79542 | MELUD119179a | MN411539 |  | [57] |
| MJB2389* | *Eremophila occidens* Chinnock, G. Paczkowska & A.R. Chapman | *Australophilae* | -22.1 | 114 | MELUD119046a | MN411459 |  |  |
| MJB2353* | *Eremophila koobabbiensis* Chinnock | *Australophilae* | -30.48 | 116.27 | MELUD119016a | MN411432 |  |  |
| RMF258 | *Eremophila behriana* (F. Muell.) F. Muell. | *Duttonia* | -33.91742 | 136.19766 | MELUD119222a | MN411571 |  |  |
| TMS14-11 | *Eremophila crassifolia* (F. Muell.) F. Muell. | *Duttonia* | -34.52 | 141.49 | - | MN486165 |  |  |
| RMF259 | *Eremophila gibbifolia* (F. Muell.) F. Muell. | *Duttonia* | -33.63334 | 136.54605 | MELUD119223a | MN411572 |  |  |
| MJB2559 | *Eremophila gibbifolia* (F. Muell.) F. Muell. | *Duttonia* | -36.62289 | 141.74581 | MELUD119112a | MN411503 |  |  |
| MJB2422 | *Eremophila spuria* Chinnock | *Arenariae* | -26.58504 | 118.60338 | MELUD119075a | MN411482 |  |  |
| MJB2431 | *Eremophila platythamnos* subsp. *platythamnos* Diels | *Arenariae* | -26.28864 | 121.08012 | MELUD119083a | MN411358 |  | [18] |
| RMF160* | *Eremophila platythamnos* subsp. *exotrachys* (Kraenzlin) Chinnock | *Arenariae* | -25.81 | 123.14 | MELUD119161a | MN411389 |  |  |
| RMF114* | *Eremophila gibsonii* F. Muell. | *Arenariae* | - | - | MELUD119124a | MN411509 |  |  |
| RMF174* | *Eremophila arenaria* Chinnock | *Arenariae* | - | - | MELUD119172a | MN411536 |  |  |
| MJB2367* | *Eremophila aureivisca* Chinnock | *Arenariae* | -28.76 | 124.33 | MELUD119030a | MN411444 |  | [57] |
| MJB2427 | *Eremophila enata* Chinnock | *Eremaeae* | -26.54832 | 119.95721 | MELUD119080a | MN411486 |  |  |
| MJB2430 | *Eremophila gilesii* subsp. *gilesii* F. Muell. (broad leaf form) | *Eremaeae* | -26.55436 | 120.66073 | MELUD119082a | MN411357 | [84, 86, 87, 88] |  |
| BB35 | *Eremophila gilesii* subsp. *variabilis* Chinnock | *Eremaeae* | -24.01503 | 117.28156 | - | MN486171 |  |  |
| MJB2481 | *Eremophila gilesii* F. Muell. | *Eremaeae* | -23.50827 | 133.85552 | MELUD119109a | MN411369 |  |  |
| MJB2409 | *Eremophila foliosissima* Kraenzlin | *Eremaeae* | -27.82393 | 117.92946 | MELUD119062a | MN411472 |  |  |
| RMF64 | *Eremophila lanceolata* Chinnock | *Eremaeae* | -24.44214 | 119.67825 | MELUD119254a | MN411592 |  |  |
| RMF118* | *Eremophila linsmithii* R.D. Henderson | *Eremaeae* | - | - | MELUD119127a | MN411511 |  |  |
| MJB2399* | *Eremophila prolata* Chinnock | *Eremaeae* | -25.69 | 118.07 | MELUD119055a | MN411466 |  |  |
| BB02 | *Eremophila freelingii* F. Muell. | *Eremaeae* | -31.21825 | 139.64822 | MELUD118892a | MN411429 | [58, 82, 84, 86, 89-91] | [97] |
| RMF248 | *Eremophila freelingii* F. Muell. | *Eremaeae* | -31.34296 | 138.56223 | MELUD119215a | MN411566 |  |  |
| RMF61 | *Eremophila spectabilis* subsp. *spectabilis* C. Gardner | *Eremaeae* | -24.9467 | 119.82532 | MELUD119253a | MN411408 |  |  |
| MJB2426 | *Eremophila spectabilis* subsp. *brevis* Chinnock | *Eremaeae* | -26.50627 | 119.86596 | MELUD119079a | MN411356 |  |  |
| MJB2418 | *Eremophila phyllopoda* subsp. *phyllopoda* Chinnock | *Eremaeae* | -26.58322 | 118.46572 | MELUD119071a | MN411354 |  |  |
| BB32 | *Eremophila phyllopoda* subsp. *obliqua* Chinnock | *Eremaeae* | -23.45561 | 117.13253 | - | MN486178 |  |  |
| RMF139 | *Eremophila rigens* Chinnock | *Eremaeae* | -24.2 | 116.87 | MELUD119145a | MN411523 |  |  |
| RMF103 | *Eremophila crenulata* Chinnock | *Eremaeae* | -25.48907 | 114.3763 | - | MN486166 |  | [18] |
| MJB2440 | *Eremophila cuneifolia* Kraenzlin | *Eremaeae* | -26.25232 | 121.68553 | MELUD119089a | MN411490 | [90] | [18] |
| RMF79 | *Eremophila cuneifolia* Kraenzlin | *Eremaeae* | -24.32425 | 119.54748 | MELUD119266a | MN411595 |  |  |
| RMF94 | *Eremophila cuneifolia* 'Big leaf' Kraenzlin | *Eremaeae* | -24.21885 | 116.53811 | MELUD119276a | MN411337 |  |  |
| RMF178* | *Eremophila goodwinii* subsp. *ecapitata* Chinnock | *Eremaeae* | - | - | MELUD119175a | MN411394 | [84] |  |
| RMF13* | *Eremophila willsii* subsp. *integrifolia* (Ewart) Chinnock | *Eremaeae* | -25.3464 | 131.0636 | CANB801460 | MN380716 |  |  |
| RMF267* | *Eremophila prostrata* Chinnock | *Eremaeae* | - | - | MELUD119228a | MN411576 |  |  |
| RMF179* | *Eremophila elderii* F. Muell. | *Eremaeae* | -24.859 | 128.02 | MELUD119176a | MN486168 | [83] |  |
| MJB2375* | *Eremophila canaliculata* Chinnock | *Eremaeae* | -25.08 | 118.28 | MELUD119037a | MN411450 |  |  |
| RMF124* | *Eremophila acrida* Chinnock | *Eremaeae* | -24.859 | 128.02 | MELUD119133a | MN411516 |  | [18] |
| MJB2410 | *Eremophila georgei* Diels | *Eremaeae* | -27.40926 | 117.90519 | MELUD119063a | MN411473 |  | [18] |
| RMF107 | *Eremophila clarkeii* Oldfield & F. Muell. | *Eremaeae* | -26.64231 | 114.61106 | MELUD119120a | MN411506 |  |  |
| MJB2452 | *Eremophila metallicorum* S. Moore | *Eremaeae* | -28.69941 | 122.07719 | MELUD119100a | MN411496 |  |  |
| MJB2459 | *Eremophila granitica* S. Moore | *Eremaeae* | -30.43796 | 120.81703 | MELUD119106a | MN411500 |  |  |
| RMF65 | *Eremophila appressa* Chinnock | *Eremaeae* | -24.34813 | 119.6922 | MELUD119255a | MN411593 |  |  |
| MJB2439 | *Eremophila punctata* Chinnock | *Eremaeae* | -26.25219 | 121.6821 | MELUD119088a | MN411327 |  |  |
| RMF87 | *Eremophila obliquepala* Chinnock | *Eremaeae* | -24.97115 | 118.42377 | MELUD119270a | MN411329 |  |  |
| MJB2450 | *Eremophila simulans* subsp. *simulans* Chinnock | *Eremaeae* | -27.80841 | 122.28447 | MELUD119098a | MN411363 |  |  |
| RMF110 | *Eremophila simulans* subsp. *megacalyx* Chinnock | *Eremaeae* | -27.21885 | 116.3052 | MELUD119122a | MN411374 |  |  |
| RMF134* | *Eremophila conglomerata* Chinnock | *Eremaeae* | -27.98 | 119.29 | MELUD119142a | MN411521 |  |  |
| MJB2433 | *Eremophila pungens* Chinnock | *Eremaeae* | -26.14454 | 121.14867 | MELUD119084a | MN411487 |  |  |
| RMF56 | *Eremophila* aff. *pungens* 'Meekatharra' Chinnock | *Eremaeae* | -25.97086 | 118.9006 | MELUD119250a | MN486194 |  |  |
| RMF69 | *Eremophila incisa* Chinnock | *Eremaeae* | -24.22421 | 119.78342 | MELUD119259a | MN411594 |  |  |
| MJB2423 | *Eremophila flabellata* Chinnock | *Eremaeae* | -26.58504 | 118.60338 | MELUD119076a | MN411483 |  |  |
| MJB2414 | *Eremophila glutinosa* Chinnock | *Eremaeae* | -27.20193 | 118.01662 | MELUD119067a | MN411477 |  |  |
| RMF66 | *Eremophila petrophila* subsp. *petrophila* Chinnock | *Eremaeae* | -24.34786 | 119.6922 | MELUD119256a | MN411409 |  |  |
| RMF102 | *Eremophila petrophila* subsp. *densa* Chinnock | *Eremaeae* | -25.46398 | 115.18572 | MELUD119116a | MN411371 |  |  |
| MJB2444 | *Eremophila hughesii* | *Eremaeae* | -26.02358 | 121.96667 | MELUD119092a | MN411361 |  |  |
| MJB2406 | *Eremophila exilifolia* F. Muell. | *Eremaeae* | -28.26294 | 117.85965 | MELUD119060a | MN411469 |  |  |
| MJB2449 | *Eremophila shonae* subsp. *diffusa* Chinnock | *Eremaeae* | -27.64173 | 121.63891 | MELUD119097a | MN411362 |  |  |
| MJB2442 | *Eremophila annosocaulis* Chinnock | *Eremaeae* | -26.27867 | 121.83372 | MELUD119091a | MN411491 |  |  |
| RMF120* | *Eremophila battii* F. Muell. | *Eremaeae* | -26.59 | 120.22 | MELUD119129a | MN411513 |  |  |
| RMF181* | *Eremophila battii* 'Granite Peaks' F. Muell. | *Eremaeae* | -25.67 | 121.3 | MELUD119177a | MN411331 |  |  |
| MJB2398* | *Eremophila magnifica* subsp. *magnifica* Chinnock | *Eremaeae* | -22.25 | 117.98 | MELUD119054a | MN411350 |  |  |
| MJB2382* | *Eremophila pallida* Chinnock | *Eremaeae* | -23.5 | 126.37 | MELUD119041a | MN411454 |  |  |
| RMF119* | *Eremophila pentaptera* J. Black | *Carnosae* | - | - | MELUD119128a | MN411512 |  |  |
| RMF265* | *Eremophila macdonnellii* (broad leaf) F. Muell. | *Platycalyx* | - | - | MELUD119227a | MN411575 |  |  |
| RMF284* | *Eremophila arbuscula* Chinnock | *Eremophila* | -26.5 | 143.81 | MELUD119236a | MN411581 |  |  |
| MJB2401 | *Eremophila oppositifolia* subsp. *angustifolia* (S. Moore) Chinnock | *Eremophila* | -29.70754 | 117.0847 | MELUD118629a | MN411343 |  |  |
| RMF277* | *Eremophila oppositifolia* subsp. *rubra* (White & Francis) Chinnock | *Eremophila* | - | - | MELUD119232a | MN411407 |  |  |
| MJB2368* | *Eremophila reticulata* Chinnock | *Eremophila* | -24.32 | 117.23 | MELUD119031a | MN411445 |  |  |
| MJB2390* | *Eremophila cryptothrix* Chinnock | *Eremophila* | -23.16 | 117.63 | MELUD119047a | MN411460 |  |  |
| BB06 | *Eremophila rotundifolia* F. Muell. | *Eremophila* | -29.57689 | 135.07367 | MELUD119011a | MN411430 |  |  |
| MJB2415 | *Eremophila spathulata* W.V. Fitzg. | *Eremophila* | -27.04921 | 118.17319 | MELUD119068a | MN411478 |  |  |
| MJB2385* | *Eremophila rigida* Chinnock | *Eremophila* | -24.94 | 118.7 | MELUD119044a | MN411457 |  |  |
| RMF100 | *Eremophila tietkensisii* F. Muell. | *Eremophila* | -24.678 | 115.25732 | MELUD119114a | MN411504 |  |  |
| RMF74 | *Eremophila tietkensisii* 'Hamersby' F. Muell. | *Eremophila* | -23.04108 | 118.85095 | MELUD119263a | MN411336 |  |  |
| BB17 | *Eremophila tietkensisii* F. Muell. | *Eremophila* | -23.91603 | 128.74172 | - | MN486181 |  |  |
| RMF121* | *Eremophila mirabilis* Chinnock | *Eremophila* | -29.35 | 121.29 | MELUD119130a | MN411514 |  |  |
| MJB2400* | *Eremophila platycalyx* subsp. *platycalyx* F. Muell. | *Eremophila* | -26.51 | 118.5 | MELUD119056a | MN411351 |  |  |
| RMF105 | *Eremophila platycalyx* subsp. *platycalyx* F. Muell. | *Eremophila* | -26.28749 | 114.41776 | MELUD119118a | MN411373 |  |  |
| RMF169* | *Eremophila platycalyx* 'Leonora' F. Muell. | *Eremophila* | -28.85 | 121.64 | MELUD119169a | MN411330 |  |  |
| RMF85 | *Eremophila platycalyx* 'Milgen' F. Muell. | *Eremophila* | -24.94533 | 118.45538 | MELUD119269a | MN411597 |  |  |
| RMF54 | *Eremophila platycalyx* 'Granite' F. Muell. | *Eremophila* | -26.57933 | 118.47971 | MELUD119248a | MN411335 |  |  |
| MJB2413 | *Eremophila macmillaniana* C. Gardner | *Eremophila* | -27.20172 | 118.01705 | MELUD119066a | MN411476 |  |  |
| RMF232 | *Eremophila polyclada* (F. Muell.) F. Muell. | *Platychilus* | -34.13445 | 141.31102 | MELUD119208a | MN411560 |  |  |
| RMF236 | *Eremophila bignoniiflora* (Benth.) F. Muell. | *Platychilus* | -34.13842 | 141.33371 | MELUD119211a | MN411562 | [92] |  |
| RMF162* | *Eremophila arachnoides* subsp. *arachnoides* Chinnock | *Pholidia* | - | - | MELUD119163a | MN411391 |  |  |
| RMF147* | *Eremophila arachnoides* subsp. *tenera* Chinnock | *Pholidia* | - | - | MELUD119153a | MN411384 |  |  |
| RMF23* | *Eremophila dalyana* F. Muell. | *Pholidia* | -27.7322 | 142.9833 | CANB602768 | MN380715 | [84, 88, 93] |  |
| RMF113* | *Eremophila stenophylla* Chinnock | *Pholidia* | - | - | MELUD119123a | MN411508 |  |  |
| MJB2411 | *Eremophila pantonii* F. Muell. | *Pholidia* | -27.27829 | 117.97772 | MELUD119064a | MN411474 |  |  |
| MJB2402 | *Eremophila scoparia* (R. Br.) | *Pholidia* | -29.52997 | 117.16625 | MELUD119057a | MN411467 |  |  |
| RMF194 | *Eremophila scoparia* (R. Br.) | *Pholidia* | -32.03243 | 120.74231 | MELUD119187a | MN411546 |  |  |
| TMS14-13 | *Eremophila scoparia* (R. Br.) | *Pholidia* | -34.79 | 141.63 | - | MN486180 |  |  |
| RMF231 | *Eremophila scoparia* (R. Br.) | *Pholidia* | -34.38153 | 141.33385 | MELUD119207a | MN411559 |  |  |
| RMF249 | *Eremophila scoparia* (R. Br.) | *Pholidia* | -31.33969 | 138.55145 | MELUD119216a | MN411567 |  |  |
| MJB2441 | *Eremophila youngii* subsp. *youngii* F. Muell. | *Pholidia* | -26.25773 | 121.75411 | MELUD119090a | MN411360 |  |  |
| RMF73 | *Eremophila youngii* subsp. *lepidota* Chinnock | *Pholidia* | -22.69531 | 119.94966 | MELUD119262a | MN411413 |  |  |
| RMF227 | *Eremophila divaricata* subsp. *divaricata* (F. Muell.) F. Muell. | *Sentis* | -34.28471 | 142.21725 | MELUD119205a | MN411400 |  |  |
| RMF150* | *Eremophila decussata* Chinnock | *Decussatae* | - | - | MELUD119156a | MN411530 |  |  |
| MJB2424 | *Eremophila malacoides* Chinnock | *Decussatae* | -26.49111 | 119.78154 | MELUD119077a | MN411484 |  |  |
| RMF148* | *Eremophila delisseri* F. Muell. | *Decussatae* | - | - | MELUD119154a | MN411528 |  |  |
| RMF149* | *Eremophila dendritica* Chinnock | *Decussatae* | -30.79 | 125.37 | MELUD119155a | MN411529 |  |  |
| RMF159* | *Eremophila mackinlayi* subsp. *mackinlayi* F. Muell. | *Hygrophanae* | -28.07 | 117.75 | MELUD119160a | MN411388 |  |  |
| RMF106 | *Eremophila stronglylophylla* F. Muell. | *Hygrophanae* | -26.54748 | 114.51396 | MELUD119119a | MN411505 |  |  |
| MJB2454 | *Eremophila hygrophana* Chinnock | *Hygrophanae* | -28.88918 | 121.37331 | MELUD119102a | MN411497 |  |  |
| RMF115* | *Eremophila ovata* Chinnock | *Hygrophanae* | - | - | MELUD119125a | MN411510 |  |  |
| MJB2378* | *Eremophila fasciata* Chinnock | *Hygrophanae* | -26.88 | 118.66 | MELUD119038a | MN411451 |  |  |
| MJB2396* | *Eremophila warnesii* Chinnock | *Hygrophanae* | -25.58 | 117.98 | MELUD119052a | MN411464 |  |  |
| RMF183* | *Eremophila macgillivrayi* J. Black | *Contortae* | - | - | MELUD122771a | MN486175 |  |  |
| RMF104 | *Eremophila pterocarpa* subsp. *pterocarpa* W.V. Fitzg. | *Contortae* | -25.57502 | 114.2225 | MELUD119117a | MN411372 |  |  |
| NG6475 | *Eremophila pterocarpa* subsp. *acicularis* Chinnock | *Contortae* | -25.32736 | 120.86552 | MEL2390092 | MN411342 |  |  |
| RMF05* | *Eremophila bowmanii* subsp. *bowmanii* F. Muell. | *Eriocalyx* | -30.6075 | 145.7986 | CANB602733 | MN380713 |  |  |
| RMF152* | *Eremophila bowmanii* subsp. *nutans* Chinnock | *Eriocalyx* | - | - | MELUD119158a | MN411386 |  |  |
| MJB2405 | *Eremophila forrestii* subsp. *forrestii* F. Muell. | *Eriocalyx* | -28.94095 | 117.82689 | MELUD119059a | MN411352 |  |  |
| RMF142 | *Eremophila forrestii* subsp. *capensis* Chinnock | *Eriocalyx* | -22.11 | 114.04 | MELUD119148a | MN411381 |  |  |
| MJB2455 | *Eremophila glandulifera* Chinnock | *Eriocalyx* | -28.88918 | 121.37331 | MELUD119103a | MN411498 |  |  |
| MJB2407 | *Eremophila punicea* S. Moore | *Eriocalyx* | -28.26294 | 117.85965 | MELUD119061a | MN411471 |  |  |
| MJB2421 | *Eremophila latrobei* subsp. *latrobei* F. Muell. | *Eriocalyx* | -26.58322 | 118.46572 | MELUD119074a | MN411355 | [58, 86, 89, 94] | Semple *et al.* (unpub.) |
| NG6477 | *Eremophila latrobei* subsp. *latrobei* F. Muell. | *Eriocalyx* | -25.06827 | 120.6555 | MEL2390089 | MN411341 |  |  |
| RMF72 | *Eremophila latrobei* subsp. *filiformis* Chinnock | *Eriocalyx* | -22.78621 | 120.00291 | MELUD119261a | MN411412 |  |  |
| RMF268* | *Eremophila latrobei* subsp. *glabra* (L.S. Smith) Chinnock | *Eriocalyx* | - | - | MELUD119229a | MN411405 |  |  |
| RMF165* | *Eremophila compacta* subsp. *compacta* S. Moore | *Eriocalyx* | - | - | MELUD119166a | MN411392 |  |  |
| RMF135* | *Eremophila compacta* subsp. *fecunda* Chinnock | *Eriocalyx* | -26.37 | 117.34 | MELUD119143a | MN411380 |  |  |
| RMF101 | *Eremophila* aff. *compacta* 'Kennedy Ranges' S. Moore | *Eriocalyx* | -24.66147 | 115.17414 | MELUD119115a | MN486190 |  |  |
| MJB2370* | *Eremophila conferta* Chinnock | *Eriocalyx* | -24.2 | 116.44 | MELUD119033a | MN411447 |  |  |
| RMF93 | *Eremophila conferta* Chinnock | *Eriocalyx* | -24.32825 | 116.87273 | MELUD119275a | MN411599 |  |  |
| RMF97 | *Eremophila rhegos* Chinnock | *Eriocalyx* | -24.7399 | 116.06875 | MELUD119278a | MN411601 |  |  |
| RMF77 | *Eremophila* aff. *rhegos* Chinnock | *Eriocalyx* | -23.73387 | 119.72642 | MELUD119265a | MN486195 |  |  |
| RMF130 | *Eremophila congesta* Chinnock | *Eriocalyx* | -26.37 | 120.24 | MELUD119139a | MN411519 |  |  |
| MJB2436 | *Eremophila citrina* Chinnock | *Eriocalyx* | -26.20009 | 121.26826 | MELUD119086a | MN411489 |  |  |
| RMF59 | *Eremophila demissa* Chinnock | *Eriocalyx* | -25.48275 | 119.26962 | MELUD119252a | MN411591 |  |  |
| RMF82 | *Eremophila demissa* | *Eriocalyx* | -24.61331 | 118.60313 | MELUD119268a | MN486196 |  |  |
| MJB2416 | *Eremophila jucunda* subsp. *jucunda* Chinnock | *Eriocalyx* | -27.04921 | 118.17319 | MELUD119069a | MN411353 |  |  |
| RMF75 | *Eremophila jucunda* subsp. *pulcherrima* Chinnock | *Eriocalyx* | -23.04178 | 118.85008 | MELUD119264a | MN411414 |  |  |
| MJB2419 | *Eremophila retropila* Chinnock | *Eriocalyx* | -26.58322 | 118.46572 | MELUD119072a | MN411480 |  |  |
| MJB2425 | *Eremophila margarethae* S. Moore | *Eriocalyx* | -26.49111 | 119.78154 | MELUD119078a | MN411485 |  |  |
| RMF55 | *Eremophila* aff. *anomala* 'Meekatharra' Chinnock | *Eriocalyx* | -25.97079 | 118.90061 | MELUD119249a | MN486193 |  |  |
| MJB2372* | *Eremophila muelleriana* C. Gardner | *Eriocalyx* | -25.11 | 117.58 | MELUD119035a | MN411448 |  |  |
| MJB2371* | *Eremophila eriocalyx* F. Muell. | *Eriocalyx* | -27.74 | 115.81 | MELUD119034a | MN411326 |  |  |
| RMF68 | *Eremophila caespitosa* Chinnock | *Eriocalyx* | -24.22388 | 119.78333 | MELUD119258a | MN411328 |  |  |
| RMF128* | *Eremophila obovata* subsp. *obovata* L.S. Smith | *Eriocalyx* | - | - | MELUD119137a | MN411378 |  |  |
| RMF127* | *Eremophila obovata* subsp. *glabriuscula* (L.S. Smith) Chinnock | *Eriocalyx* | - | - | MELUD119136a | MN411377 |  |  |
| MJB2397* | *Eremophila maitlandii* F. Muell. ex Benth. | *Eriocalyx* | -24.9 | 113.67 | MELUD119053a | MN411465 |  |  |
| RMF99 | *Eremophila recurva* Chinnock | *Eriocalyx* | -25.26767 | 115.56365 | MELUD119280a | MN411602 |  |  |
| RMF141* | *Eremophila physocalyx* Chinnock | *Eriocalyx* | -32.2 | 121.76 | MELUD119147a | MN411525 |  |  |
| MJB2417 | *Eremophila lachnocalyx* C. Gardner | *Eriocalyx* | -26.58322 | 118.46572 | MELUD119070a | MN411479 |  |  |
| BB48a | *Eremophila ballythunnensis* Buirchell & A.P.Br. | *Eriocalyx* | -26.071 | 115.7495 | - | MN486160 |  |  |
| RMF96 | *Eremophila yinnetharrensis* Buirchell & A.P.Br. | *Eriocalyx* | -24.71546 | 116.1329 | MELUD119277a | MN411600 |  |  |
| RMF92 | *Eremophila buirchellii* A.P.Br. | *Eriocalyx* | -24.32963 | 116.86642 | MELUD119274a | MN411598 |  |  |
| MJB2448 | *Eremophila campanulata* Chinnock | *Campanulatae* | -26.58209 | 122.80328 | MELUD119096a | MN411495 |  |  |
| RMF81 | *Eremophila humilis* Chinnock | *Campanulatae* | -24.61329 | 118.60305 | MELUD119267a | MN411596 |  |  |
| RMF175* | *Eremophila tetraptera* C.T. White | *Subscissocarpae* | - | - | MELUD119173a | MN411537 |  | [18] |
| RMF109 | *Eremophila laanii* F. Muell. | *Amphichilus* | -26.83023 | 115.44857 | MELUD119121a | MN411507 |  |  |
| MJB2412 | *Eremophila longifolia* (R. Br.) F. Muell. | *Amphichilus* | -27.27829 | 117.97772 | MELUD119065a | MN411475 | [83, 89, 94, 95] | [109], Semple *et al.* (unpub.) |
| RMF225 | *Eremophila longifolia* (R. Br.) F. Muell. | *Amphichilus* | -34.28999 | 142.1859 | MELUD119204a | MN411557 |  |  |
| RMF241 | *Eremophila longifolia* (R. Br.) F. Muell. | *Amphichilus* | -31.3024 | 138.83626 | MELUD119212a | MN411563 |  |  |
| MJB2351* | *Eremophila rostrata* subsp. *trifida* Chinnock | *Amphichilus* | -29.46 | 116.48 | MELUD119015a | MN411346 |  |  |
| RMF233 | *Eremophila maculata* subsp. *maculata* (Ker Gawler) F. Muell. | *Stenochilus* | -34.13435 | 141.31055 | MELUD119209a | MN411401 | [96] |  |
| TMS14-08 | *Eremophila maculata* subsp. *maculata* (Ker Gawler) F. Muell. | *Stenochilus* | -34.3 | 142.23 | - | MN486176 |  |  |
| MJB2484 | *Eremophila maculata* subsp. *maculata* (Ker Gawler) F. Muell. | *Stenochilus* | -23.3011 | 133.7875 | MELUD119111a | MN411370 |  |  |
| MJB2462 | *Eremophila maculata* subsp. *brevifolia* (Benth) Chinnock | *Stenochilus* | -30.95319 | 121.1434 | MELUD119108a | MN411368 |  |  |
| RMF166* | *Eremophila maculata* subsp. *filifolia* Chinnock | *Stenochilus* | -21.09 | 119.83 | MELUD119167a | MN411393 |  |  |
| RMF189 | *Eremophila racemosa* (Endl.) F. Muell. | *Stenochilus* | -32.35143 | 119.74735 | MELUD119183a | MN411543 |  |  |
| RMF213 | *Eremophila calorhados* Diels | *Stenochilus* | -32.9413 | 120.34617 | MELUD119200a | MN411555 |  | [57] |
| MJB2350* | *Eremophila denticulata* subsp. *denticulata* F. Muell. | *Stenochilus* | -33.89 | 119.85 | MELUD119014a | MN411345 |  |  |
| MJB2365* | *Eremophila denticulata* subsp. *trisulcata* Chinnock | *Stenochilus* | -33.44 | 123.46 | MELUD119028a | MN411347 |  | [18] |
| RMF163* | *Eremophila splendens* Chinnock | *Stenochilus* | -26.27 | 113.89 | MELUD119164a | MN411531 |  |  |
| RMF188 | *Eremophila biserrata* Chinnock | *Stenochilus* | -32.4673 | 119.75875 | MELUD119182a | MN411542 |  | [18] |
| MJB2437 | *Eremophila glabra* subsp. *glabra* (R. Br.) Ostenf. | *Stenochilus* | -26.22714 | 121.445 | MELUD119087a | MN411359 |  | [18, 25] |
| RMF251 | *Eremophila glabra* subsp. *glabra* (R. Br.) Ostenf. | *Stenochilus* | -33.03468 | 136.79251 | MELUD119218a | MN411402 |  |  |
| RMF215 | *Eremophila glabra* subsp. *albicans* (Bartling) Chinnock | *Stenochilus* | -33.15895 | 120.09927 | MELUD119201a | MN411398 |  |  |
| RMF153* | *Eremophila glabra* subsp. *carnea* Chinnock | *Stenochilus* | -28.19 | 114.25 | MELUD119159a | MN411387 |  |  |
| RMF122* | *Eremophila glabra* subsp. *chlorella* (Gandoger) Chinnock | *Stenochilus* | -32.02 | 115.92 | MELUD119131a | MN411375 |  |  |
| RMF151* | *Eremophila glabra* subsp. *verrucosa* Chinnock | *Stenochilus* | -29.42 | 121.89 | MELUD119157a | MN411385 |  |  |
| RMF53 | *Eremophila glabra* 'Paynes Find' (R. Br.) Ostenf. | *Stenochilus* | -29.2907 | 117.59231 | MELUD119247a | MN411334 |  |  |
| RMF184 | *Eremophila glabra* 'York' (R. Br.) Ostenf. | *Stenochilus* | -31.94689 | 116.97558 | MELUD119178a | MN411332 |  |  |
| MJB2456 | *Eremophila decipiens* subsp. *decipiens* Ostenf. | *Stenochilus* | -29.727 | 120.82164 | MELUD119104a | MN411366 |  | [18, 57] |
| RMF190 | *Eremophila decipiens* subsp. *decipiens* Ostenf. | *Stenochilus* | -32.34968 | 119.74717 | MELUD119184a | MN411367 |  |  |
| MJB2356* | *Eremophila subteretifolia* Chinnock | *Stenochilus* | -32.82 | 119.52 | MELUD119017a | MN411433 |  | [18] |
| MJB2369* | *Eremophila hillii* E.A. Shaw | *Stenochilus* |  |  | MELUD119032a | MN411446 |  |  |
| RMF03* | *Eremophila subfloccosa* subsp. *subfloccosa* Benth. | *Stenochilus* | -31.7794 | 117.4839 | CBG9805407 | MN411340 |  | [18] |
| RMF205 | *Eremophila subfloccosa* subsp. *glandulosa* Chinnock | *Stenochilus* | -33.24094 | 123.22465 | MELUD119195a | MN411397 |  | [57] |
| RMF253 | *Eremophila subfloccosa* subsp. *lanata* Chinnock | *Stenochilus* | -33.2052 | 136.0592 | MELUD119220a | MN411403 |  |  |
| RMF140 | *Eremophila calcicola* R.W. Davis | *Stenochilus* | -33.699 | 122.6 | MELUD119146a | MN411524 |  |  |
| RMF196 | *Eremophila gibbosa* Chinnock | *Virides* | -32.02489 | 120.73406 | MELUD119188a | MN411547 |  | [18] |
| MJB2359* | *Eremophila virens* C. Gardner | *Virides* | -30.92 | 118.23 | MELUD119019a | MN411435 |  | [18, 57] |
| RMF146* | *Eremophila undulata* Chinnock | *Virides* | - | - | MELUD119152a | MN411527 | [18] |  |
| MJB2404 | *Eremophila serrulata* (Cunn. ex A. DC.) Druce | *Virides* | -28.94095 | 117.82689 | MELUD119058a | MN411468 |  | [18, 27] |
| RMF261* | *Eremophila serrulata* (Cunn. ex A. DC.) Druce | *Virides* | - | - | MELUD119225a | MN411574 |  |  |
| MJB2434 | *Eremophila alternifolia* R.Br. | *Scariosepalae* | -26.14361 | 121.2885 | MELUD119085a | MN411488 | [58, 89] | [9, 97], Semple *et al.* (unpub.) |
| RMF247 | *Eremophila alternifolia* R.Br. | *Scariosepalae* | -31.34098 | 138.56705 | MELUD119214a | MN411565 | [97] |  |
| RMF202 | *Eremophila alternifolia* 'Broadleaf' R. Br. | *Scariosepalae* | -32.02849 | 122.86559 | MELUD119192a | MN411333 | [98] |  |
| RMF168* | *Eremophila purpurascens* Chinnock | *Scariosepalae* | -32.2 | 121.68 | MELUD119168a | MN411533 |  |  |
| MJB2451 | *Eremophila abietina* subsp. *abietina* Kraenzlin | *Pulchrisepalae* | -28.48833 | 122.42218 | MELUD119099a | MN411364 |  |  |
| MJB2366* | *Eremophila abietina* subsp. *ciliata* Chinnock | *Pulchrisepalae* | -28.76 | 124.33 | MELUD119029a | MN411348 |  |  |
| RMF70 | *Eremophila fraserii* subsp. *fraserii* F. Muell. | *Pulchrisepalae* | -24.26112 | 119.71781 | MELUD119260a | MN411411 | [90] |  |
| RMF88 | *Eremophila fraserii* subsp. *fraserii* F. Muell. | *Pulchrisepalae* | -25.11716 | 118.28693 | MELUD119271a | MN411415 |  |  |
| RMF90 | *Eremophila fraserii* subsp. *parva* Chinnock | *Pulchrisepalae* | -25.17587 | 117.68446 | MELUD119273a | MN411416 |  |  |
| MJB2447 | *Eremophila galeata* Chinnock | *Pulchrisepalae* | -26.5822 | 123.05959 | MELUD119095a | MN411494 | [99] | [18], Semple *et al.* (unpub.) |
| RMF58 | *Eremophila galeata* Chinnock | *Pulchrisepalae* | -25.97263 | 118.89314 | MELUD119251a | MN411590 |  |  |
| MJB2445 | *Eremophila ramiflora* Dell | *Pulchrisepalae* | -25.85671 | 122.5485 | MELUD119093a | MN411492 |  |  |
| RMF67 | *Eremophila flaccida* subsp. *flaccida* Chinnock | *Pulchrisepalae* | -24.26702 | 119.71123 | MELUD119257a | MN411410 |  |  |
| RMF161 | *Eremophila flaccida* subsp. *attenuata* Chinnock | *Pulchrisepalae* | -25.18 | 115.49 | MELUD119162a | MN411390 |  |  |
| MJB2446 | *Eremophila miniata* Gardner | *Pulchrisepalae* | -26.53575 | 123.18797 | MELUD119094a | MN411493 |  |  |
| MJB2388* | *Eremophila miniata* Gardner | *Pulchrisepalae* | -30.46 | 117.5 | MELUD119045a | MN411458 |  |  |
| RMF129* | *Eremophila neglecta* J. Black | *Pulchrisepalae* | - | - | MELUD119138a | MN411518 | [84] | [7, 18] |
| MJB2360* | *Eremophila viscida* Endl. | *Pulchrisepalae* | -30.7 | 116.87 | MELUD119020a | MN411436 |  |  |
| RMF292 | *Eremophila lucida* Chinnock | *Pulchrisepalae* | -32.27932 | 120.26237 | MELUD119240a | MN411585 |  | [18, 26, 57] |
| MJB2395* | *Eremophila oldfieldii* subsp. *oldfieldii* F. Muell. | *Pulchrisepalae* | -30.29 | 116.68 | MELUD119051a | MN411349 |  |  |
| MJB2453 | *Eremophila oldfieldii* subsp. *angustifolia* (S. Moore) Chinnock | *Pulchrisepalae* | -28.7434 | 122.02129 | MELUD119101a | MN411365 |  |  |
| MJB2420 | *Eremophila linearis* Chinnock | *Pulchrisepalae* | -26.58322 | 118.46572 | MELUD119073a | MN411481 |  | [18], Semple *et al.* (unpub.) |
| MJB2483 | *Eremophila duttonii* F. Muell. | *Pulchrisepalae* | -23.30118 | 133.78698 | MELUD119110a | MN411502 | [58, 82] | [97, 110] |
| RMF136* | *Eremophila duttonii* F. Muell. | *Pulchrisepalae* | - | - | MELUD119144a | MN411522 |  |  |
| RMF250 | *Eremophila duttonii* F. Muell. | *Pulchrisepalae* | -31.3384 | 138.52936 | MELUD119217a | MN411568 |  |  |
| MJB2374* | *Eremophila grandiflora* A.P. Br & Buirchell | *Pulchrisepalae* | -29.09 | 117.19 | MELUD119036a | MN411449 |  |  |
| RMF49 | *Eremophila* sp. Wubin North (R. Wait 8305) | - | -30.09924 | 116.62247 | MELUD119245a | MN486192 |  |  |
| RMF89 | *Eremophila* sp. Yarlalweelor (M. Greeve 41) | - | -25.295 | 118.06142 | MELUD119272a | MN486197 |  |  |
| BB55 | *Eremophila* sp. Little Sandy Desert (A. A. Mitchell PRP 1263) | - | -25.49556 | 122.65833 | - | MN486177 |  |  |
| RMF31* | *Myoporum montanum* R. Br. | - | -27.615 | 152.9069 | CBG7810920 | MN411419 | [100, 101]<br>[102] | [102] |
| RMF46 | *Myoporum boninense* subsp. *australe* Chinnock | - | -35.70108 | 150.19981 | MELUD119244a | MN411589 |  |  |
| RMF281 | *Myoporum insulare* R. Br. | - | -33.15676 | 138.06808 | MELUD119235a | MN411580 | [103] | [111] |
| RMF45 | *Myoporum acuminatum* R. Br. | - | -35.70231 | 150.18958 | MELUD119243a | MN411588 | [104] | [112] |
| RMF29* | *Myoporum caprarioides* Benth. | - | -34.6833 | 117.95 | CBG7901554 | MN411420 |  |  |
| RMF32* | *Myoporum petiolatum* Chinnock | - | -38.4167 | 144.1333 | CBG18760 | MN380717 |  |  |
| RMF280 | *Myoporum petiolatum* Chinnock | - | -32.82301 | 138.18399 | MELUD119234a | MN411579 |  |  |
| RMF209 | *Myoporum tetrandrum* (Labill.) Domin. | - | -33.79922 | 121.96305 | MELUD119196a | MN411551 |  |  |
| RMF38* | *Myoporum betcheanum* subsp. *betcheanum* L. Smith | - | -28.05 | 152.3833 | CBG8604103 | MN411424 |  |  |
| RMF25* | *Myoporum parvifolium* R. Br. | - | -33.3667 | 136.6333 | CBG36473 | MN411417 |  |  |
| RMF260 | *Myoporum brevipes* Benth. | - | -33.65805 | 136.98543 | MELUD119224a | MN411573 |  |  |
| NTBG061301 | *Myoporum sandwicense* subsp. *sandwicense* A. Gray | - | 21.461 | -157.991 | PTBG075975 | MN411618 | [105] |  |
| RMF42 | *Myoporum bateae* F. Muell. | - | -35.77214 | 150.169 | MELUD119242a | MN411587 |  |  |
| RMF35* | *Myoporum platycarpum* subsp. *platycarpum* R. Br. | - | -33.1833 | 137.0333 | CBG8209994 | MN411338 | [103] |  |
| RMF214 | *Glycocystis beckeri* (F. Muell.) Chinnock | - | -32.95107 | 120.32957 | MELUD119023a | MN411439 |  |  |
| FTBG64271* | *Bontia daphnoides* L. | - | 25.67469 | -80.276301 | FTG4719 | MN411428 | [106-108] |  |
| RMF220* | *Calamphoreus inflatus* (C. Gardner) Chinnock | - | -33.1 | 119.66 | MELUD119024a | MN411440 |  |  |
| MJB2383* | *Diocirea microphylla* Chinnock | - | -31.01 | 120.9 | MELUD119042a | MN411455 |  |  |
| RMF131* | *Diocirea violacea* Chinnock | - | - | - | MELUD119022a | MN411438 |  |  |
| RMF210 | *Diocirea violacea* Chinnock | - | -32.93161 | 121.61952 | MELUD119197a | MN411552 |  |  |
| RMF275* | *Leucophyllum frutescens* I.M. Johnst | - | - | - | MELUD119231a | MN411577 |  |  |
| RMF288* | *Leucophyllum frutescens* I.M. Johnst | - | - | - | MELUD119025a | MN411441 |  |  |
| RMF286* | *Leucophyllum zygophyllum* I.M. Johnst | - | - | - | MELUD119237a | MN411582 |  |  |

**Table S2.** Statistical analysis on individual heatmap metabolic cluster using general linear models. Generalised linear models were used to test for significant differences of functional annotations among the investigated specimens within a heatmap metabolic cluster (HMC) background. Models were generated for each HMC-attribute combination using the clogloc model, while excluding specimens with missing functional annotation. Each test was conducted using the anova.manyglm() function from the mvabund package. A p-value threshold of 0.05 was considered significant.

| HMC | Attribute | Res.Df | Df.diff | Deviation | p |
| --- | --- | --- | --- | --- | --- |
| I | leaf resin | 288 | 1 | 45 | **0.001** |
|  | leaf hairiness | 289 | 1 | 4 | 0.299 |
|  | pollination | 289 | 1 | 18 | **0.001** |
|  | antibacterial activity | 124 | 1 | 17 | **0.002** |
|  | biogeography | 286 | 4 | 19 | 0.134 |
| II | leaf resin | 288 | 1 | 31 | **0.001** |
|  | leaf hairiness | 289 | 1 | 21 | **0.003** |
|  | pollination | 289 | 1 | 0 | 0.991 |
|  | antibacterial activity | 124 | 1 | 2 | 0.898 |
|  | biogeography | 286 | 4 | 44 | **0.004** |
| III | leaf resin | 288 | 1 | 29 | **0.001** |
|  | leaf hairiness | 289 | 1 | 7 | **0.044** |
|  | pollination | 289 | 1 | 9 | **0.025** |
|  | antibacterial activity | 124 | 1 | 12 | **0.015** |
|  | biogeography | 286 | 4 | 23 | 0.056 |
| IV | leaf resin | 288 | 1 | 44 | **0.001** |
|  | leaf hairiness | 289 | 1 | 7 | 0.362 |
|  | pollination | 289 | 1 | 20 | **0.004** |
|  | antibacterial activity | 124 | 1 | 5 | 0.547 |
|  | biogeography | 286 | 4 | 37 | 0.085 |
| V | leaf resin | 288 | 1 | 125 | **0.001** |
|  | leaf hairiness | 289 | 1 | 34 | **0.001** |
|  | pollination | 289 | 1 | 58 | **0.001** |
|  | antibacterial activity | 124 | 1 | 92 | **0.001** |
|  | biogeography | 286 | 4 | 27 | 0.405 |
| VI | leaf resin | 288 | 1 | 206 | **0.001** |
|  | leaf hairiness | 289 | 1 | 98 | **0.001** |
|  | pollination | 289 | 1 | 97 | **0.001** |
|  | antibacterial activity | 124 | 1 | 130 | **0.001** |
|  | biogeography | 286 | 4 | 173 | **0.001** |
| VII | leaf resin | 288 | 1 | 36 | **0.001** |
|  | leaf hairiness | 289 | 1 | 4 | 0.792 |
|  | pollination | 289 | 1 | 5 | 0.733 |
|  | antibacterial activity | 124 | 1 | 7 | 0.436 |
|  | biogeography | 286 | 4 | 6 | 1 |
| VIII | leaf resin | 288 | 1 | 95 | **0.001** |
|  | leaf hairiness | 289 | 1 | 27 | **0.001** |
|  | pollination | 289 | 1 | 2 | 0.753 |
|  | antibacterial activity | 124 | 1 | 6 | 0.174 |
|  | biogeography | 286 | 4 | 28 | **0.049** |
| IX | leaf resin | 288 | 1 | 112 | **0.001** |
|  | leaf hairiness | 289 | 1 | 30 | **0.008** |
|  | pollination | 289 | 1 | 18 | 0.163 |
|  | antibacterial activity | 124 | 1 | 23 | 0.065 |
|  | biogeography | 286 | 4 | 129 | **0.001** |
| X | leaf resin | 288 | 1 | 278 | **0.001** |
|  | leaf hairiness | 289 | 1 | 52 | **0.001** |
|  | pollination | 289 | 1 | 9 | 0.292 |
|  | antibacterial activity | 124 | 1 | 106 | **0.001** |
|  | biogeography | 286 | 4 | 104 | **0.001** |

**Table S3.**
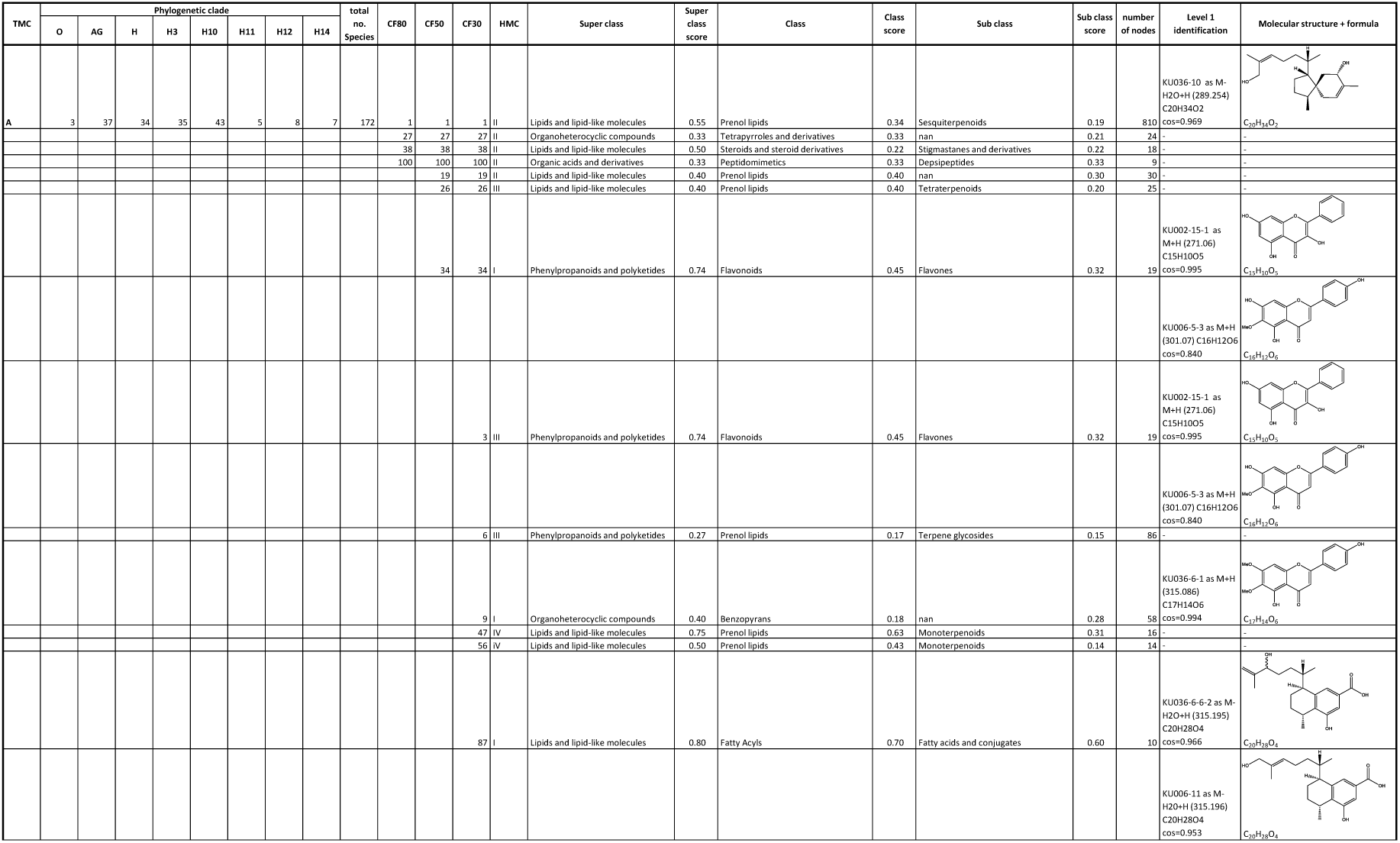
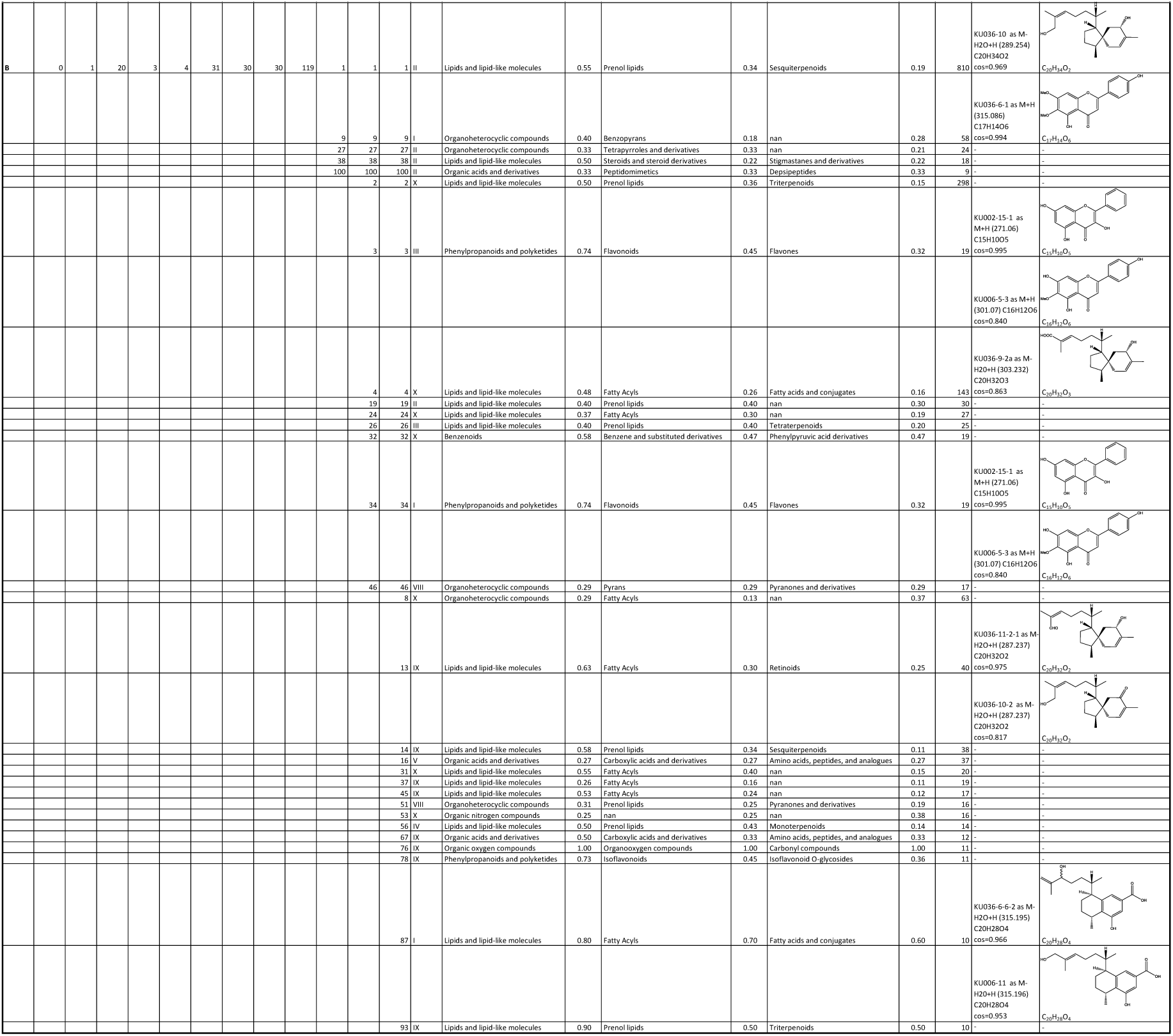
Chemical comparison of tanglegram metabolic cluster A and B. Comparison of the chemical content between tanglegram metabolic cluster (TMC) A and B. The underlying metabolic cluster analysis is based on the 100 largest chemical families of the generated molecular network of Myoporeae. Number of specimens from major phylogenetic clades are shown with “O”: outgroup, “AG”: clade A to G, “H3”: clade H3, “H10”: clade H10, “H11”: clade H11, “H12”: clade H12, “H14”: clade H14 and “H”: remaining clade H associated taxa. Dominant chemical families are depicted for each TMC, with “CF80”: chemical families present in 80% of the specimens, as well as “CF50” and “CF30”, depicting 50% and 30% abundancy of a given family among the given specimens, respectively. Different levels of predicted chemical classification and corresponding scores are shown for each chemical family. If present, level 1 identifications and their associated molecular structures are displayed.

**Table S4.**
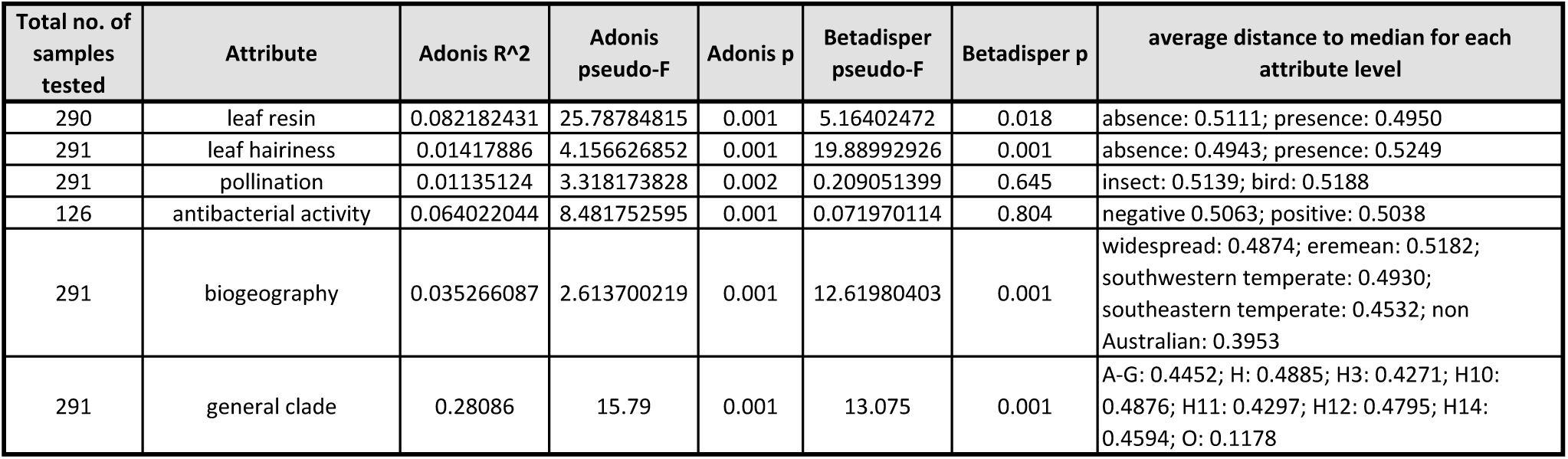
Multivariate statistical analysis involving the chemical space of Myoporeae. Permutational multivariate analysis of variance (PERMANOVA) analysis were conducted involving all samples associated with a given functional annotation. Results of the Adonis2() and betadisper() functions are shown, which give information about statistical significance and data dispersion of tested attribute levels, respectively.

**Table S5.** Univariate tests and indicator species analysis on individual chemical families. To determine individual chemical families as overall significantly associated to a functional annotation, an univariate test was conducted using generalised linear models including all specimens with corresponding metadata to receive adjusted p-values. An additional indicator species analysis using the indval() function was applied to indicate the direction of annotation (level) for each chemical family.

| Test | Attribute |  | 1 | 2 | 3 | 4 | 5 | 6 | 7 | 8 | 9 | 10 | 11 | 12 | 13 | 14 | 15 | 16 | 17 | 18 |
| --- | --- | --- | --- | --- | --- | --- | --- | --- | --- | --- | --- | --- | --- | --- | --- | --- | --- | --- | --- | --- |
| GLM univariate test | leaf resin | p | 0.9 | **0.001** | **0.004** | **0.001** | 1 | 0.101 | 0.34 | **0.001** | **0.001** | **0.003** | 0.917 | 0.8 | 0.109 | **0.014** | 0.87 | **0.001** | **0.001** | 0.059 |
| indicator species analysis |  | p | 0.076 | **0.001** | **0.001** | **0.001** | 0.402 | **0.004** | **0.004** | **0.001** | **0.001** | **0.001** | 0.119 | **0.042** | **0.003** | **0.001** | 0.050 | **0.001** | **0.001** | **0.001** |
|  |  | level | absence | presence | presence | presence | presence | absence | absence | presence | presence | presence | presence | presence | presence | presence | absence | presence | presence | absence |
| GLM univariate test | leaf hairiness | p | 1 | 0.67 | 0.468 | 0.541 | 0.398 | 1 | 1 | 0.425 | 1 | 0.786 | 0.066 | 0.786 | 0.964 | 1 | 1 | 0.058 | 0.061 | 1 |
| indicator species analysis |  | p | 0.528 | **0.023** | **0.005** | **0.038** | **0.007** | 1.000 | 0.768 | **0.032** | 0.913 | 0.062 | **0.001** | 0.050 | 0.101 | 0.773 | 0.690 | **0.007** | **0.003** | 1.000 |
|  |  | level | absence | presence | absence | presence | absence | presence | presence | presence | absence | presence | absence | presence | presence | presence | absence | presence | absence | presence |
| GLM univariate test | pollination | p | 1 | 1 | 1 | 1 | 1 | 0.273 | 0.97 | 1 | **0.006** | 1 | 1 | 1 | 1 | 0.888 | 1 | 0.349 | 1 | 1 |
| indicator species analysis |  | p | 1.000 | 0.780 | 0.901 | 0.592 | 0.128 | **0.020** | **0.029** | 0.195 | **0.001** | 0.076 | 1.000 | 1.000 | 0.233 | 0.093 | 0.419 | **0.003** | 0.177 | 0.140 |
|  |  | level | insect | bird | insect | bird | bird | insect | bird | bird | bird | bird | bird | insect | bird | insect | bird | bird | bird | bird |
| GLM univariate test | antibacterial activity | p | 1 | **0.001** | 1 | 0.956 | 1 | 0.099 | 1 | **0.001** | 0.558 | **0.001** | 1 | 1 | 1 | 1 | 1 | **0.034** | 0.967 | 1 |
| indicator species analysis |  | p | 0.575 | **0.001** | 0.600 | **0.033** | 0.216 | **0.002** | 0.418 | **0.001** | **0.010** | **0.001** | 0.604 | 1.000 | 0.117 | 0.646 | 0.376 | **0.002** | **0.045** | 0.164 |
|  |  | level | negative | positive | positive | positive | positive | negative | negative | positive | positive | positive | negative | negative | positive | positive | positive | positive | positive | negative |
| GLM univariate test | biogeography | p | 1 | 0.129 | 1 | **0.001** | 0.902 | 0.392 | 1 | 0.513 | 1 | 0.999 | 0.729 | 0.143 | 0.925 | 0.138 | 1 | 1 | 0.925 | 1 |
| indicator species analysis |  | p | 0.127 | 0.260 | 0.541 | **0.041** | 0.167 | 0.220 | 0.782 | 0.308 | 0.713 | 0.089 | **0.044** | 0.153 | 0.370 | 0.193 | 0.136 | 0.538 | **0.049** | **0.010** |
|  |  | level | widespread | south-western temperate | non Australian | eremean | south-eastern temperate | widespread | widespread | eremean | south-western temperate | south-western temperate | south-eastern temperate | eremean | eremean | eremean | non Australian | south-western temperate | south-eastern temperate | non Australian |

| Test | Attribute |  | 19 | 20 | 21 | 22 | 23 | 24 | 25 | 26 | 27 | 28 | 29 | 30 | 31 | 32 | 33 | 34 | 35 | 36 |
| --- | --- | --- | --- | --- | --- | --- | --- | --- | --- | --- | --- | --- | --- | --- | --- | --- | --- | --- | --- | --- |
| GLM univariate test | leaf resin | p | 1 | 1 | 1 | **0.009** | 0.13 | **0.001** | 0.444 | 1 | 0.9 | 1 | 0.754 | 0.789 | **0.001** | **0.001** | **0.009** | 0.141 | 1 | 0.428 |
| indicator species analysis |  | p | 0.795 | 0.538 | 0.475 | **0.002** | **0.004** | **0.001** | **0.020** | 0.712 | 0.092 | 0.386 | 0.050 | **0.037** | **0.001** | **0.001** | **0.001** | **0.005** | 0.407 | 0.075 |
|  |  | level | presence | presence | absence | presence | presence | presence | presence | presence | presence | absence | absence | presence | presence | presence | absence | presence | presence | presence |
| GLM univariate test | leaf hairiness | p | 1 | 1 | 1 | 0.379 | 1 | **0.01** | 1 | 1 | 1 | 1 | 1 | 1 | 0.978 | 0.776 | 1 | 1 | 1 | 1 |
| indicator species analysis |  | p | 0.178 | 0.305 | 0.795 | **0.026** | 1.000 | **0.003** | 0.243 | 1.000 | 0.461 | 0.227 | 0.721 | 0.232 | 0.139 | 0.050 | 0.397 | 1.000 | 0.144 | 0.638 |
|  |  | level | absence | presence | presence | presence | presence | presence | presence | absence | absence | presence | presence | presence | presence | presence | presence | presence | presence | presence |
| GLM univariate test | pollination | p | 1 | 1 | 1 | 0.12 | 1 | 1 | 1 | 1 | 1 | 1 | 1 | 1 | 1 | 1 | 1 | 1 | 1 | 0.886 |
| indicator species analysis |  | p | 0.897 | 0.385 | 1.000 | **0.002** | 0.329 | 0.251 | 0.079 | 0.776 | 0.696 | 0.796 | 0.249 | 0.531 | 0.435 | 0.064 | 1.000 | 0.122 | 0.782 | **0.031** |
|  |  | level | insect | bird | insect | bird | insect | bird | bird | insect | bird | bird | insect | insect | bird | bird | bird | bird | bird | bird |
| GLM univariate test | antibacterial activity | p | 1 | 1 | 1 | 0.95 | 0.562 | **0.002** | **0.027** | 1 | 1 | 1 | 0.103 | 1 | 0.089 | **0.001** | 0.999 | 0.192 | 1 | 1 |
| indicator species analysis |  | p | 0.709 | 0.406 | 0.669 | **0.027** | **0.020** | **0.001** | **0.002** | 0.285 | 0.785 | 1.000 | **0.001** | 0.373 | **0.002** | **0.001** | 0.123 | **0.004** | 0.234 | 0.590 |
|  |  | level | negative | positive | negative | positive | positive | positive | positive | positive | positive | positive | negative | positive | positive | positive | negative | positive | positive | positive |
| GLM univariate test | biogeography | p | 0.951 | 0.999 | 1 | 1 | 1 | 0.443 | 1 | 1 | 0.902 | 1 | 1 | 0.951 | 0.951 | 0.94 | 1 | 0.951 | 1 | 1 |
| indicator species analysis |  | p | 0.231 | 0.698 | **0.008** | 0.909 | 0.666 | 0.385 | 0.631 | 0.107 | 0.710 | **0.003** | 0.637 | 0.430 | 0.586 | 0.657 | 0.347 | 0.063 | 0.837 | 0.562 |
|  |  | level | south-western temperate | eremean | non Australian | south-western temperate | eremean | eremean | eremean | non Australian | widespread | non Australian | eremean | eremean | eremean | eremean | south-eastern temperate | non Australian | south-western temperate | widespread |

| Test | Attribute |  | 37 | 38 | 39 | 40 | 41 | 42 | 43 | 44 | 45 | 46 | 47 | 48 | 49 | 50 | 51 | 52 | 53 | 54 |
| --- | --- | --- | --- | --- | --- | --- | --- | --- | --- | --- | --- | --- | --- | --- | --- | --- | --- | --- | --- | --- |
| GLM univariate test | leaf resin | p | 0.535 | **0.021** | 0.721 | 1 | 0.997 | 0.208 | 0.96 | 0.13 | 0.109 | **0.001** | 0.722 | 0.725 | **0.002** | 0.8 | **0.001** | 1 | **0.001** | 1 |
| indicator species analysis |  | p | **0.032** | **0.002** | **0.035** | 0.418 | 0.251 | **0.004** | 0.207 | **0.003** | **0.002** | **0.001** | **0.032** | **0.043** | **0.001** | **0.044** | **0.001** | 0.412 | **0.001** | 0.850 |
|  |  | level | presence | presence | absence | presence | absence | absence | presence | presence | presence | presence | absence | presence | presence | absence | presence | presence | presence | presence |
| GLM univariate test | leaf hairiness | p | 1 | 1 | 1 | 1 | 1 | 1 | 1 | 0.122 | 0.786 | 1 | 1 | 1 | 1 | 1 | 1 | 0.974 | 0.786 | 0.81 |
| indicator species analysis |  | p | 1.000 | 0.101 | 0.324 | 0.776 | 1.000 | 0.696 | 1.000 | **0.019** | **0.046** | 0.337 | 0.907 | 0.359 | 0.782 | 0.745 | 0.091 | 0.051 | **0.040** | **0.012** |
|  |  | level | presence | absence | presence | presence | presence | presence | presence | presence | presence | absence | absence | presence | presence | absence | absence | absence | presence | absence |
| GLM univariate test | pollination | p | 1 | 1 | 1 | 1 | 1 | 1 | 1 | 0.651 | 1 | 1 | 1 | 0.997 | 1 | 0.97 | 1 | 1 | 1 | 1 |
| indicator species analysis |  | p | 0.068 | 1.000 | 0.456 | 0.236 | 0.131 | 0.546 | 0.391 | **0.014** | 1.000 | 1.000 | 0.125 | 0.181 | 0.781 | **0.021** | 0.556 | 0.696 | 0.195 | 0.289 |
|  |  | level | bird | insect | insect | insect | bird | bird | bird | bird | insect | insect | bird | insect | bird | bird | insect | bird | insect | insect |
| GLM univariate test | antibacterial activity | p | 1 | 1 | 1 | 1 | 1 | 1 | 1 | 0.14 | 1 | 1 | 1 | 1 | 1 | 1 | 1 | 1 | 0.558 | 1 |
| indicator species analysis |  | p | 0.320 | 0.514 | 0.343 | 1.000 | 0.542 | 1.000 | 0.311 | **0.012** | 0.081 | 0.370 | 0.550 | 1.000 | 1.000 | 0.845 | 0.529 | 1.000 | **0.011** | 1.000 |
|  |  | level | positive | positive | negative | negative | positive | negative | positive | positive | positive | positive | negative | positive | negative | negative | positive | negative | positive | negative |
| GLM univariate test | biogeography | p | 1 | 0.999 | 1 | 1 | 1 | 1 | 1 | 1 | 0.925 | 1 | 1 | 0.916 | 1 | 1 | 1 | 1 | 0.248 | 1 |
| indicator species analysis |  | p | 0.756 | 0.987 | **0.001** | 0.610 | 0.343 | 0.688 | 1.000 | 0.814 | 0.414 | 0.277 | 0.444 | 0.255 | 0.602 | 0.935 | 0.924 | **0.009** | 0.362 | **0.003** |
|  |  | level | widespread | widespread | non Australian | eremean | non Australian | non Australian | eremean | south-western temperate | eremean | south-eastern temperate | south-eastern temperate | eremean | eremean | south-eastern temperate | south-eastern temperate | south-eastern temperate | eremean | non Australian |

| Test | Attribute |  | 55 | 56 | 57 | 58 | 59 | 60 | 61 | 62 | 63 | 64 | 65 | 66 | 67 | 68 | 69 | 70 | 71 | 72 |
| --- | --- | --- | --- | --- | --- | --- | --- | --- | --- | --- | --- | --- | --- | --- | --- | --- | --- | --- | --- | --- |
| GLM univariate test | leaf resin | p | 0.931 | 0.967 | 0.138 | 1 | 1 | 1 | 0.191 | **0.006** | **0.003** | 1 | 0.994 | 0.967 | 0.994 | 0.523 | 0.451 | 0.105 | 0.229 | 0.917 |
| indicator species analysis |  | p | 0.117 | 0.105 | **0.012** | 0.575 | 0.395 | 0.540 | **0.007** | **0.001** | **0.001** | 0.443 | 0.385 | 0.136 | 0.227 | **0.022** | **0.012** | **0.003** | **0.031** | 0.072 |
|  |  | level | absence | absence | presence | presence | presence | absence | absence | presence | presence | presence | presence | absence | presence | presence | absence | absence | presence | absence |
| GLM univariate test | leaf hairiness | p | 1 | 0.906 | 1 | 0.838 | 1 | 1 | 1 | 1 | 1 | 0.337 | 1 | 0.232 | 1 | 0.997 | 1 | 1 | 1 | 1 |
| indicator species analysis |  | p | 1.000 | 0.076 | 0.452 | 0.062 | 0.755 | 1.000 | 1.000 | 0.204 | 0.250 | **0.009** | 0.688 | **0.003** | 0.324 | 0.123 | 0.423 | 1.000 | 0.688 | 0.539 |
|  |  | level | presence | presence | presence | presence | presence | presence | presence | presence | presence | absence | presence | absence | presence | presence | presence | presence | presence | presence |
| GLM univariate test | pollination | p | 1 | 0.273 | 1 | 1 | 0.651 | 1 | 1 | 0.144 | 0.067 | 1 | 1 | **0.015** | 1 | 1 | 1 | 1 | 1 | 1 |
| indicator species analysis |  | p | 0.627 | **0.002** | 0.220 | 0.239 | **0.009** | 0.346 | 1.000 | **0.002** | **0.002** | 0.225 | 0.652 | **0.001** | 0.667 | 0.340 | 0.613 | 0.534 | 1.000 | 0.825 |
|  |  | level | bird | bird | bird | bird | bird | insect | bird | bird | bird | bird | bird | bird | bird | bird | insect | bird | bird | bird |
| GLM univariate test | antibacterial activity | p | 1 | 1 | 0.32 | 1 | 1 | 0.217 | 0.676 | 0.079 | **0.003** | 1 | 0.629 | 1 | 1 | 0.979 | 1 | 1 | 1 | 1 |
| indicator species analysis |  | p | 0.490 | 0.472 | **0.023** | 0.243 | 0.498 | **0.013** | 0.072 | **0.003** | **0.001** | 1.000 | **0.041** | 0.623 | 0.434 | 0.083 | 0.244 | 0.388 | NA | 0.673 |
|  |  | level | negative | positive | positive | positive | positive | negative | negative | positive | positive | positive | positive | negative | positive | positive | negative | negative | NA | negative |
| GLM univariate test | biogeography | p | 1 | **0.008** | 1 | 1 | 1 | 1 | 0.971 | 1 | 1 | 1 | 1 | 0.991 | 1 | 0.925 | 1 | 1 | 1 | 1 |
| indicator species analysis |  | p | **0.003** | 0.281 | 0.138 | 0.632 | 1.000 | 0.642 | 0.263 | 0.626 | 0.299 | **0.008** | 0.107 | 0.254 | 0.933 | 0.338 | 0.766 | **0.001** | 1.000 | 0.972 |
|  |  | level | non Australian | eremean | south-western temperate | eremean | south-western temperate | eremean | south-eastern temperate | south-western temperate | south-western temperate | south-eastern temperate | south-western temperate | non Australian | south-western temperate | eremean | south-eastern temperate | non Australian | eremean | south-western temperate |

| Test | Attribute |  | 73 | 74 | 75 | 76 | 77 | 78 | 79 | 80 | 81 | 82 | 83 | 84 | 85 | 86 | 87 | 88 | 89 | 90 |
| --- | --- | --- | --- | --- | --- | --- | --- | --- | --- | --- | --- | --- | --- | --- | --- | --- | --- | --- | --- | --- |
| GLM univariate test | leaf resin | p | 0.87 | 1 | 0.96 | **0.001** | 0.212 | **0.024** | **0.006** | 0.869 | 0.869 | 0.905 | 0.931 | 0.823 | 0.542 | 0.215 | 1 | 0.145 | 0.721 | 1 |
| indicator species analysis |  | p | 0.252 | 1.000 | 0.124 | **0.001** | **0.007** | **0.002** | **0.001** | 0.251 | 0.240 | 0.062 | 0.102 | 0.115 | **0.031** | **0.012** | 0.292 | **0.003** | **0.032** | 0.598 |
|  |  | level | presence | presence | absence | presence | presence | presence | absence | presence | presence | absence | presence | absence | presence | presence | absence | absence | absence | presence |
| GLM univariate test | leaf hairiness | p | 1 | 1 | 1 | 1 | 1 | 1 | 1 | 1 | 1 | 1 | 0.998 | 1 | 1 | 1 | 0.986 | 1 | 1 | 1 |
| indicator species analysis |  | p | 0.548 | 0.391 | 1.000 | 1.000 | 0.323 | 0.633 | 0.263 | 1.000 | 0.550 | 0.741 | 0.060 | 0.535 | 0.209 | 0.348 | 0.105 | 0.741 | 0.406 | 0.297 |
|  |  | level | presence | absence | presence | presence | presence | presence | presence | presence | presence | presence | absence | presence | presence | presence | presence | presence | absence | absence |
| GLM univariate test | pollination | p | 1 | 1 | 1 | 1 | 1 | 1 | 1 | 1 | 1 | 1 | 1 | 0.551 | 1 | 1 | 1 | 1 | **0.021** | 1 |
| indicator species analysis |  | p | 0.232 | 0.361 | 0.319 | 0.208 | 1.000 | 0.081 | 0.642 | 0.563 | 1.000 | 1.000 | 1.000 | **0.027** | 0.202 | 0.352 | 0.414 | 0.739 | **0.001** | 1.000 |
|  |  | level | bird | bird | bird | bird | insect | bird | insect | insect | bird | bird | insect | bird | bird | bird | bird | insect | bird | bird |
| GLM univariate test | antibacterial activity | p | 0.997 | 1 | 1 | 0.987 | 1 | 1 | 1 | 1 | 1 | 1 | 1 | 1 | **0.045** | 0.999 | 1 | 1 | 0.987 | 1 |
| indicator species analysis |  | p | 0.215 | 0.504 | 0.384 | 0.057 | 0.161 | 0.102 | 1.000 | 0.461 | 1.000 | 0.610 | 1.000 | NA | **0.002** | 0.204 | 0.307 | 0.526 | 0.094 | 0.679 |
|  |  | level | positive | negative | negative | positive | positive | positive | positive | positive | negative | negative | negative | NA | positive | positive | positive | negative | negative | negative |
| GLM univariate test | biogeography | p | 1 | 1 | 0.998 | **0.015** | 0.951 | 0.997 | 0.934 | 1 | 1 | 1 | 0.998 | 1 | 0.98 | 1 | 1 | 1 | 0.923 | 1 |
| indicator species analysis |  | p | 0.266 | 0.056 | 0.260 | 0.068 | 0.440 | 0.704 | 0.267 | 1.000 | 1.000 | 0.358 | 0.526 | 1.000 | **0.044** | 0.693 | 0.909 | 0.077 | 0.102 | 0.849 |
|  |  | level | south-western temperate | non Australian | non Australian | eremean | eremean | widespread | eremean | eremean | eremean | south-eastern temperate | widespread | eremean | south-western temperate | eremean | eremean | non Australian | south-eastern temperate | south-western temperate |

| Test | Attribute |  | 91 | 92 | 93 | 94 | 95 | 96 | 97 | 98 | 99 | 100 |
| --- | --- | --- | --- | --- | --- | --- | --- | --- | --- | --- | --- | --- |
| GLM univariate test | leaf resin | p | 0.949 | 1 | 0.827 | 1 | **0.038** | 1 | 1 | 0.87 | **0.001** | 0.115 |
| indicator species analysis |  | p | 0.266 | 1.000 | 0.060 | 0.452 | **0.003** | 1.000 | 0.326 | 0.262 | **0.001** | **0.005** |
|  |  | level | absence | presence | presence | absence | presence | presence | absence | presence | presence | presence |
| GLM univariate test | leaf hairiness | p | 0.96 | 1 | 0.997 | 1 | 1 | 0.969 | 1 | 1 | 0.988 | **0.01** |
| indicator species analysis |  | p | 0.118 | 1.000 | 0.117 | 0.102 | 1.000 | 0.169 | 0.423 | 1.000 | 0.144 | **0.001** |
|  |  | level | absence | presence | presence | absence | presence | presence | presence | presence | presence | absence |
| GLM univariate test | pollination | p | 1 | 1 | 1 | 1 | 1 | 0.886 | 1 | 1 | **0.031** | 1 |
| indicator species analysis |  | p | 1.000 | 1.000 | 1.000 | 0.659 | 0.737 | **0.034** | 0.802 | 0.585 | **0.001** | 0.718 |
|  |  | level | bird | insect | insect | insect | insect | bird | insect | insect | bird | bird |
| GLM univariate test | antibacterial activity | p | 1 | 1 | 1 | 0.989 | 0.98 | 1 | 1 | 0.998 | **0.001** | 1 |
| indicator species analysis |  | p | NA | NA | 0.811 | 0.256 | 0.096 | 0.488 | 1.000 | 0.201 | **0.001** | 1.000 |
|  |  | level | NA | NA | positive | negative | positive | negative | negative | positive | positive | negative |
| GLM univariate test | biogeography | p | 1 | 1 | 0.966 | 1 | 1 | 1 | 1 | 1 | 1 | 0.486 |
| indicator species analysis |  | p | 1.000 | 1.000 | 0.488 | 1.000 | 0.799 | 1.000 | 0.603 | 0.062 | 0.578 | 0.854 |
|  |  | level | eremean | eremean | eremean | eremean | eremean | eremean | widespread | south-western temperate | south-western temperate | widespread |

**Table S6.**
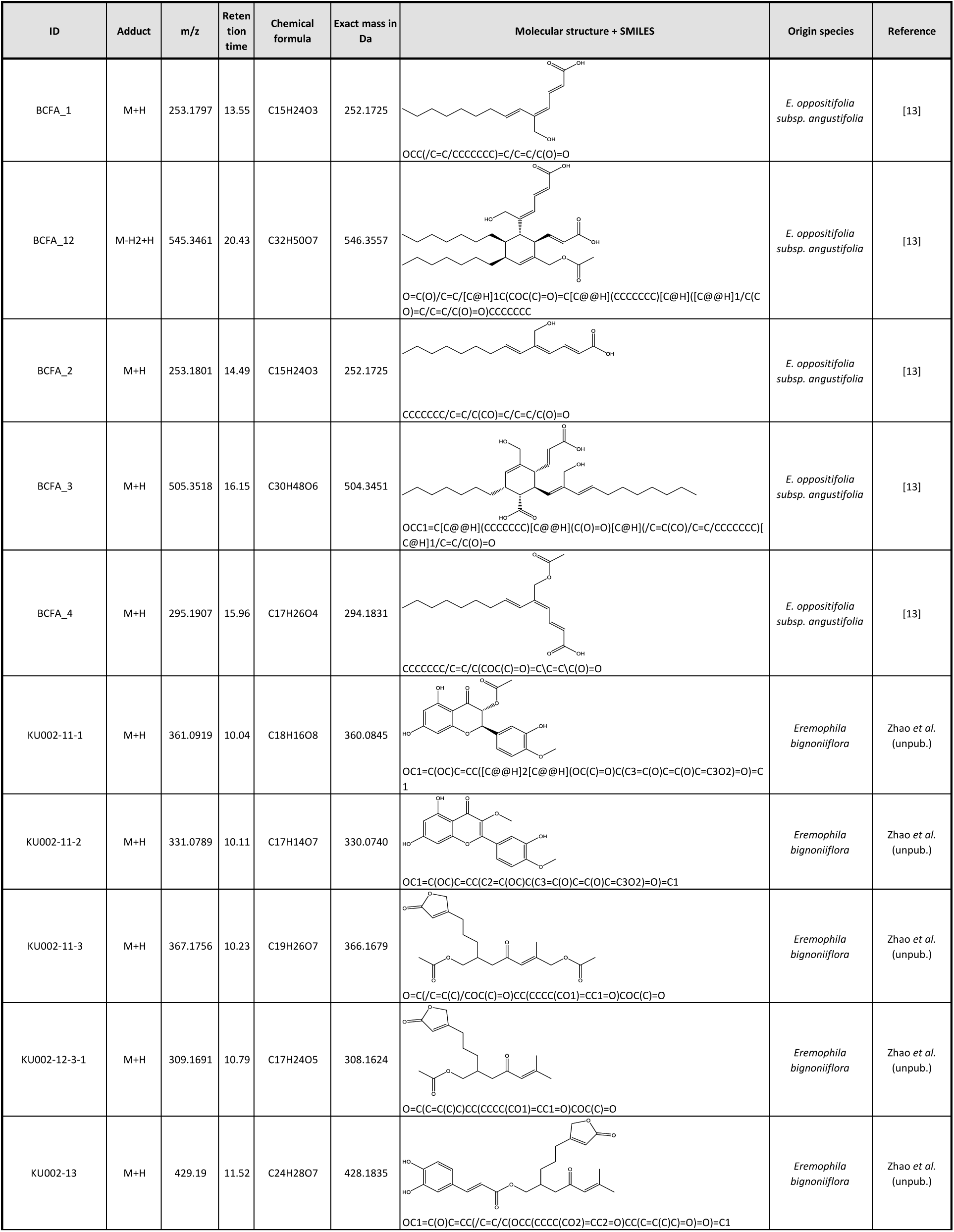
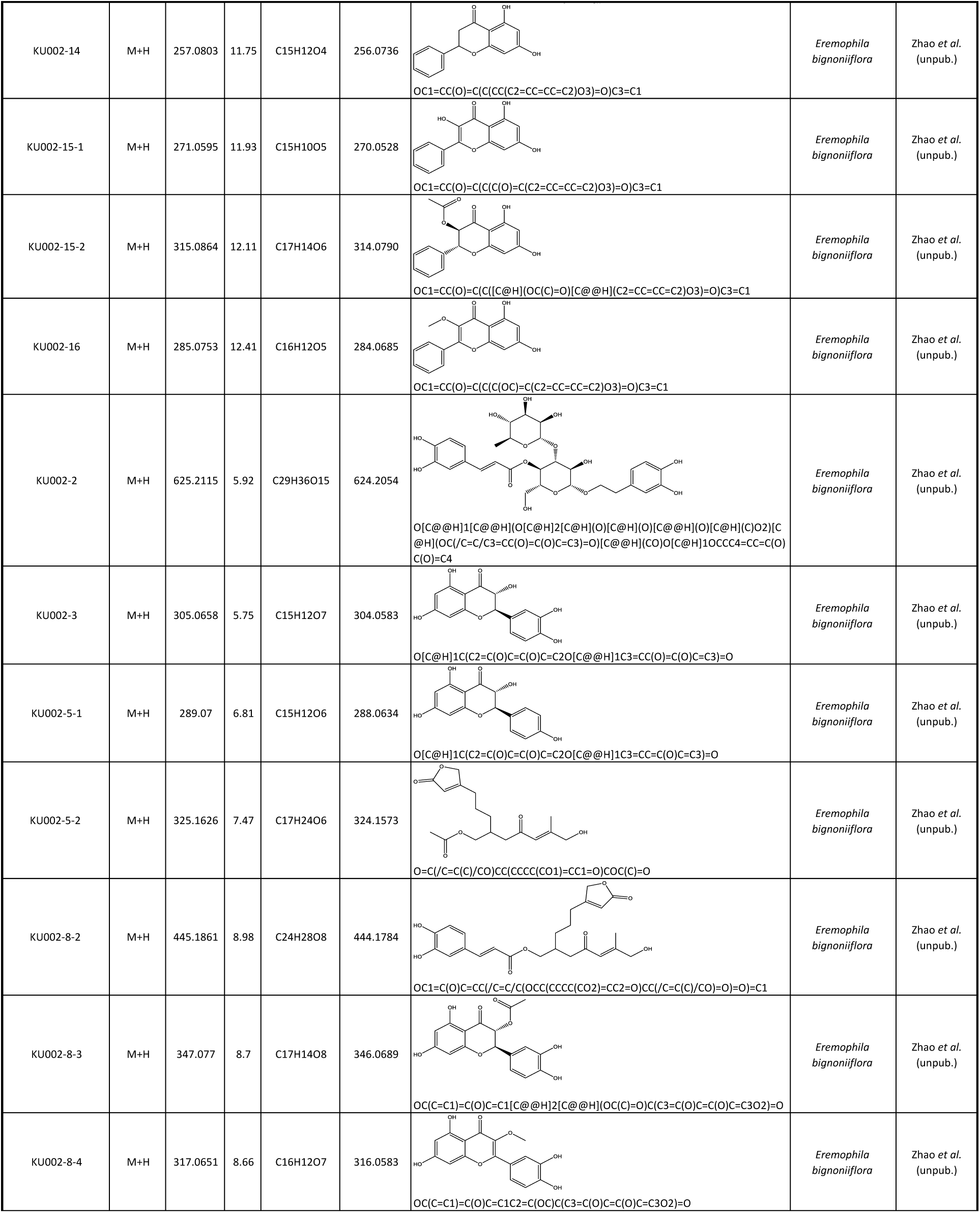
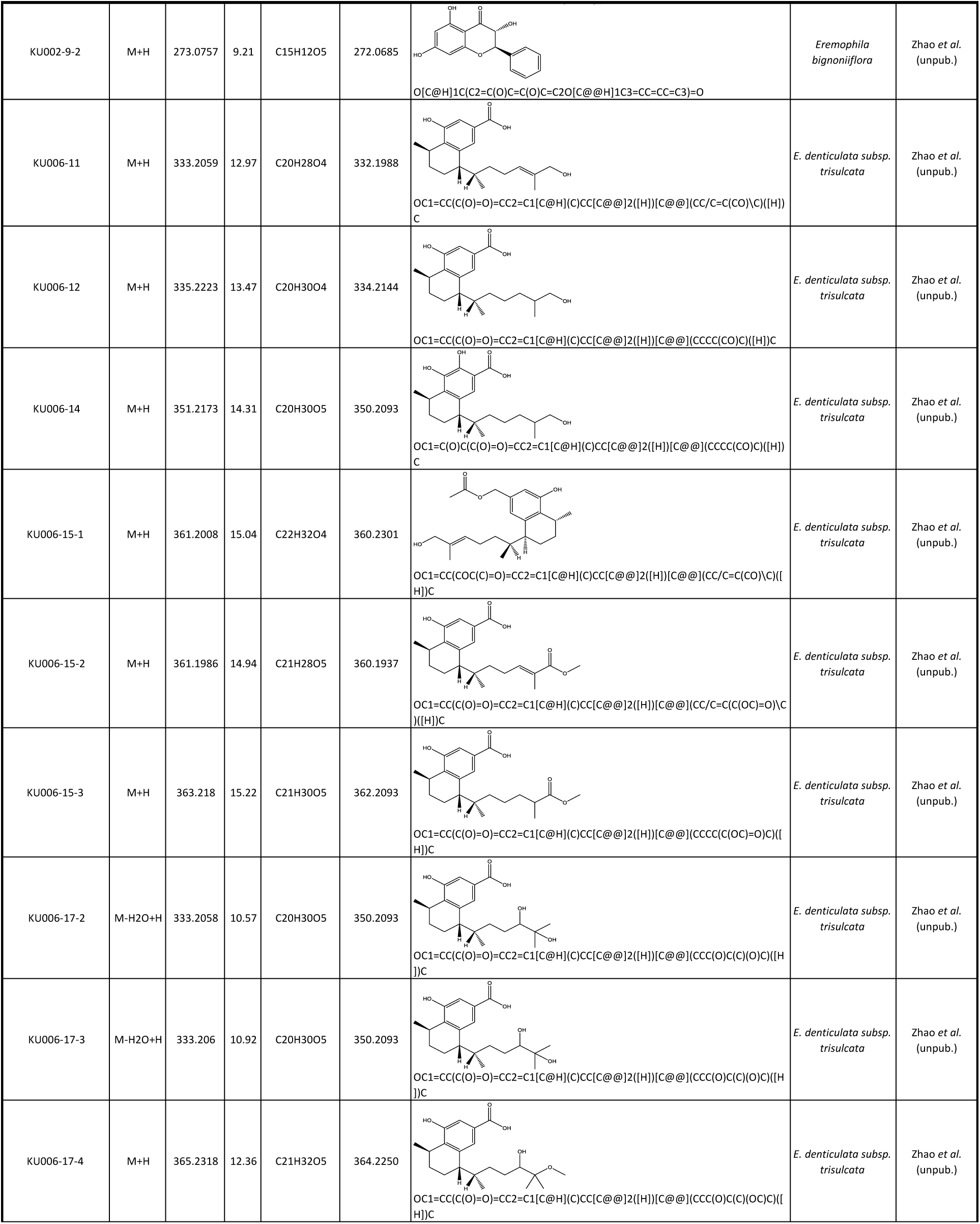
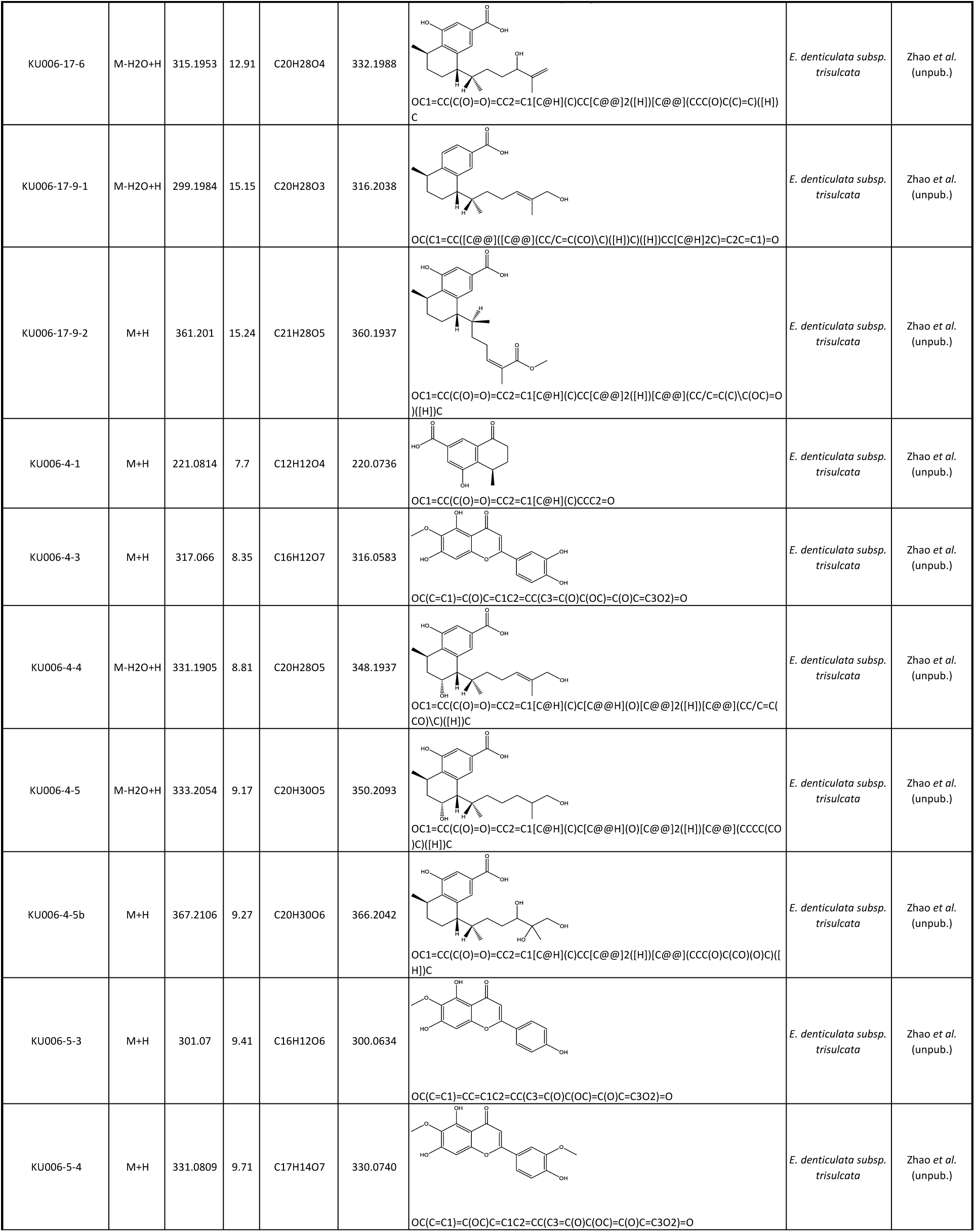
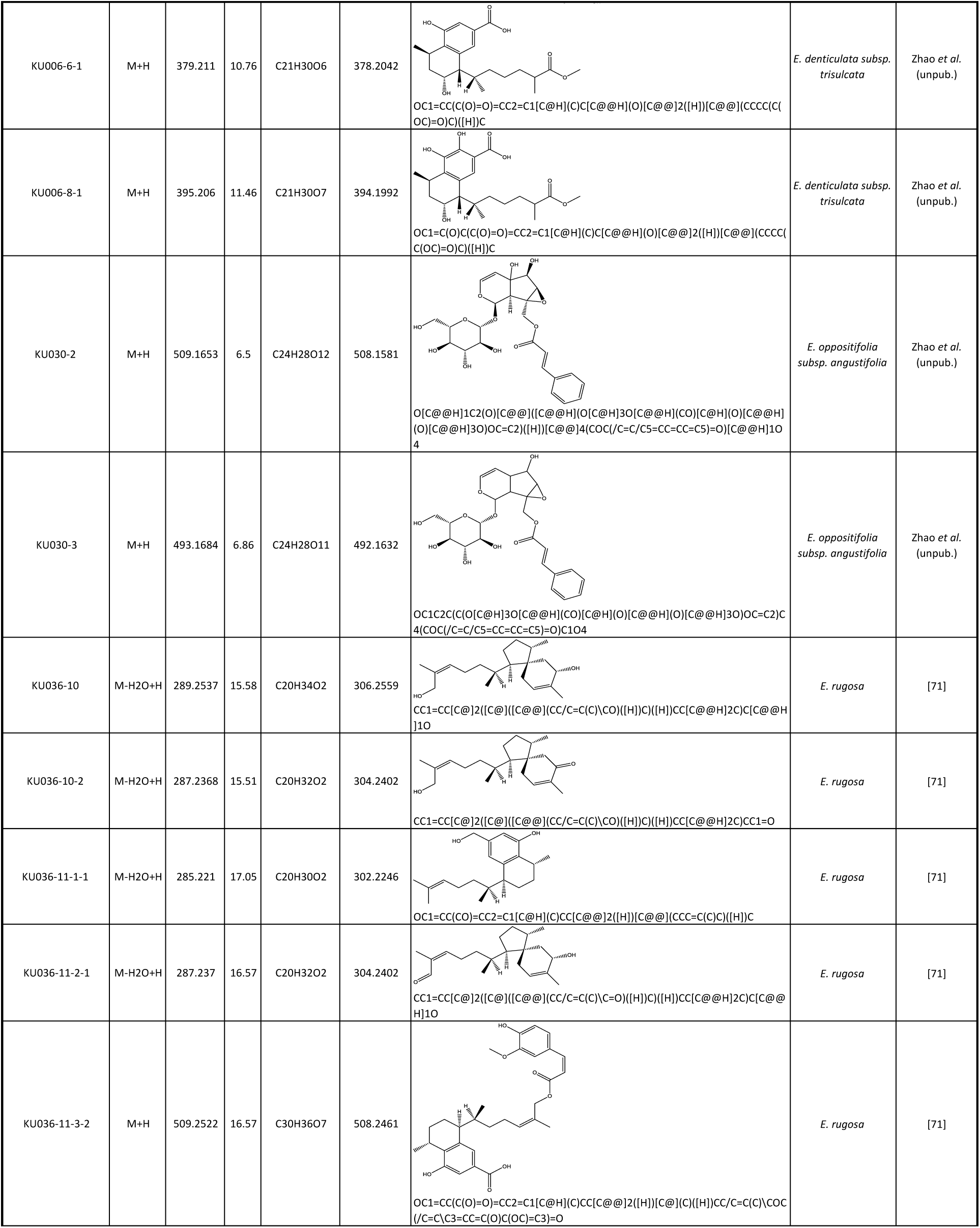
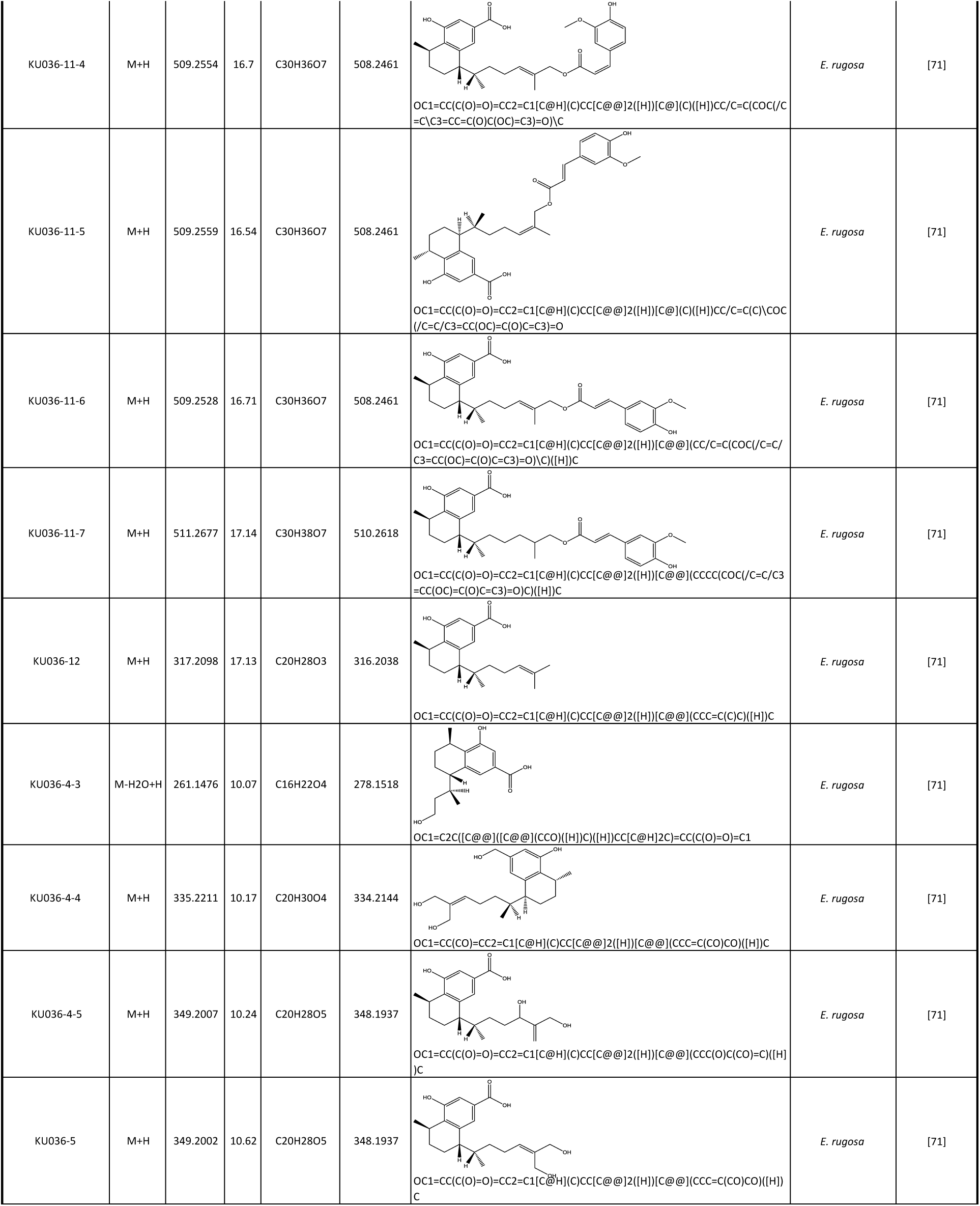
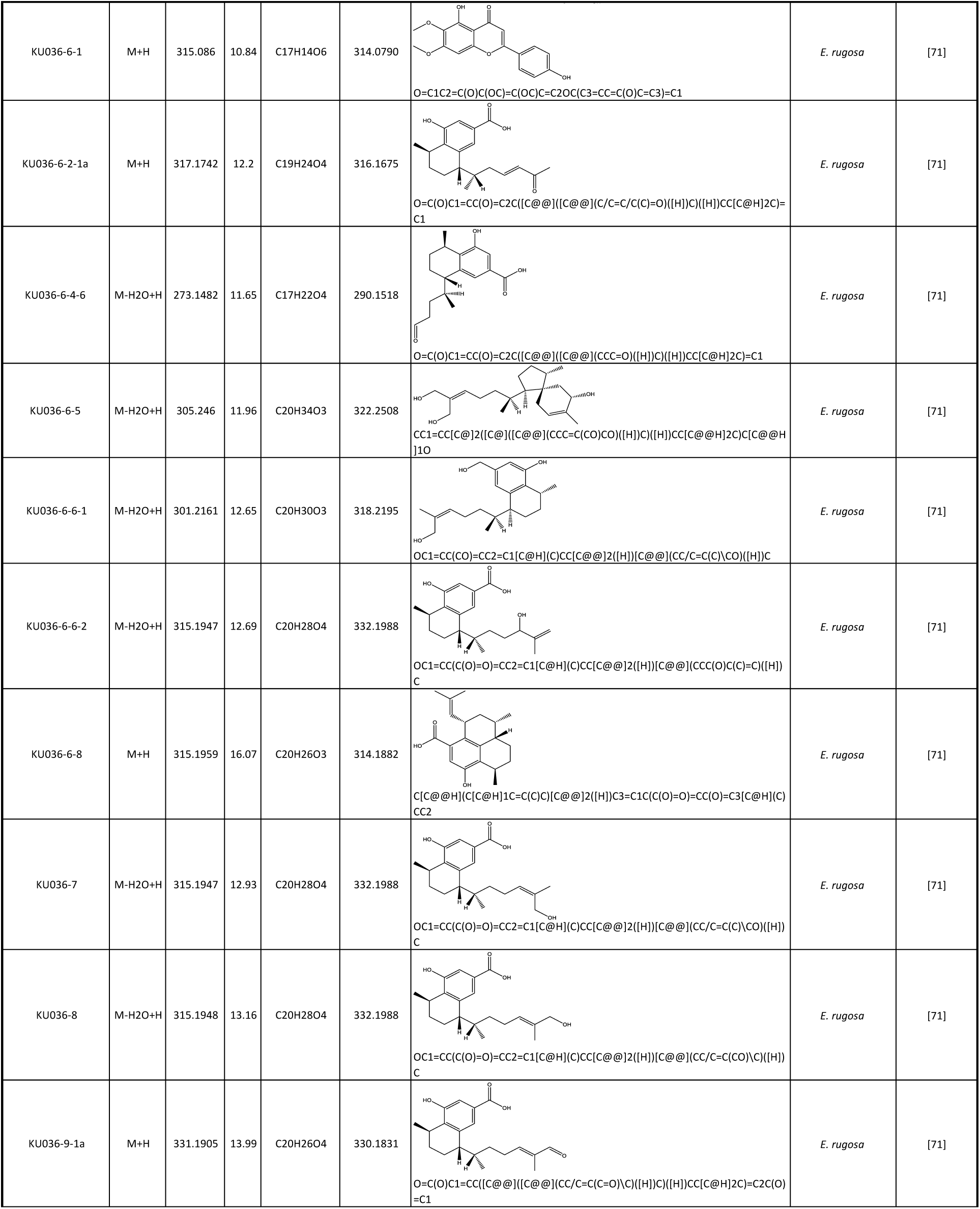
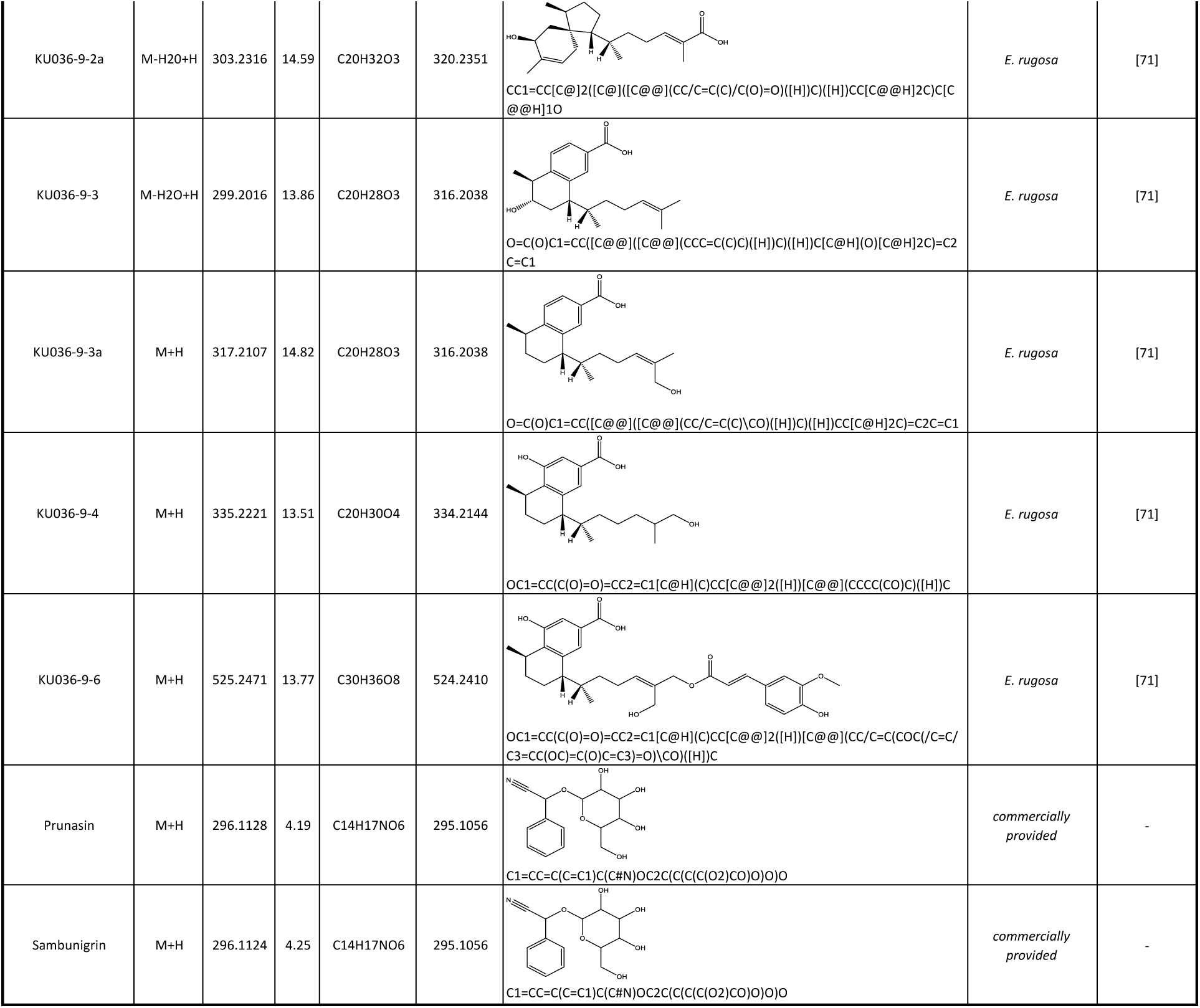
List of reference compounds used as in house spectral database. Detailed spectrometric and structural information about all reference compounds used in this study.

**Table S7.**
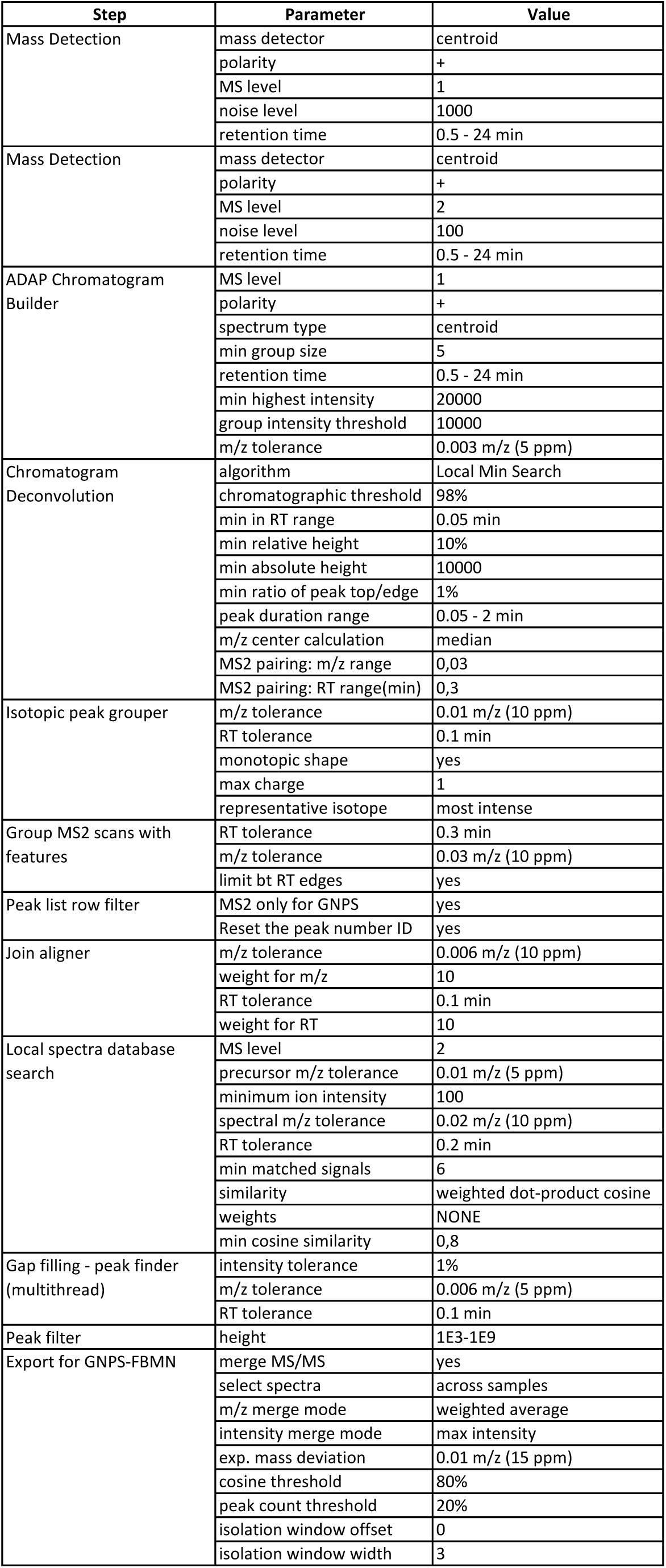
MZmine2 parameters used for raw chromatogram data processing. Detailed overview about parameters used for raw chromatogram data processing done in this study. This was carried out using the software MzMine2 [70].

**Table S8.** Molecular networking parameters. Detailed overview about parameters used for the feature-based molecular networking approach presented in this study. The analysis was carried out using the Global Natural Products Social Molecular Networking (GNPS) toolbox [38]

| Step | Parameter | Value |
| --- | --- | --- |
| Feature-based-molecular-networking (v18) | quantification table source | Mzmine |
|  | precursor ion mass tolerance | 0.02 Da |
|  | fragment ion mass tolerance | 0.02 Da |
|  | min pairs cos | 0.8 |
|  | network topk | 5 |
|  | min matched fragment ions | 8 |
|  | max connected component size | 0 |
|  | library search min matched peaks | 6 |
|  | search analogs | no |
|  | top results to report per query | 1 |
|  | score threshold | 0.7 |
|  | filter precursor window | yes |
|  | filter peaks in 50 Da window | yes |
|  | filter library | yes |
|  | normalization | no |
| NAP CCMS2 (v1.2.5) | number of cluster index | 0 |
|  | n first candidates for consensus score | 10 |
|  | accuracy for candidate search | 15 ppm |
|  | acquisition mode | positive |
|  | adduct types | [M+H], [M+Na], [M+ACN+H] |
|  | structure databases | GNPS,SUPNAT,NPATlas,CHEBI,DRUGBANK |
|  | user provided databases | Eremophila ISDB |
|  | max candidate structures in graph | 1 |
|  | cosine value to subselect | 0.5 |
|  | fusion result for consensus | yes |
|  | workflow type | MZmine |
| MS2LDA (v14) | bin width | 0.005 |
|  | LDA iterations | 1000 |
|  | min MS2 intensity | 100 |
|  | LDA free motifs | 300 |
|  | motifDB selection | all included |
|  | overlap score threshold | 0.3 |
|  | probability value threshold | 0.1 |
|  | topX node | 5 |

**Table S9.**
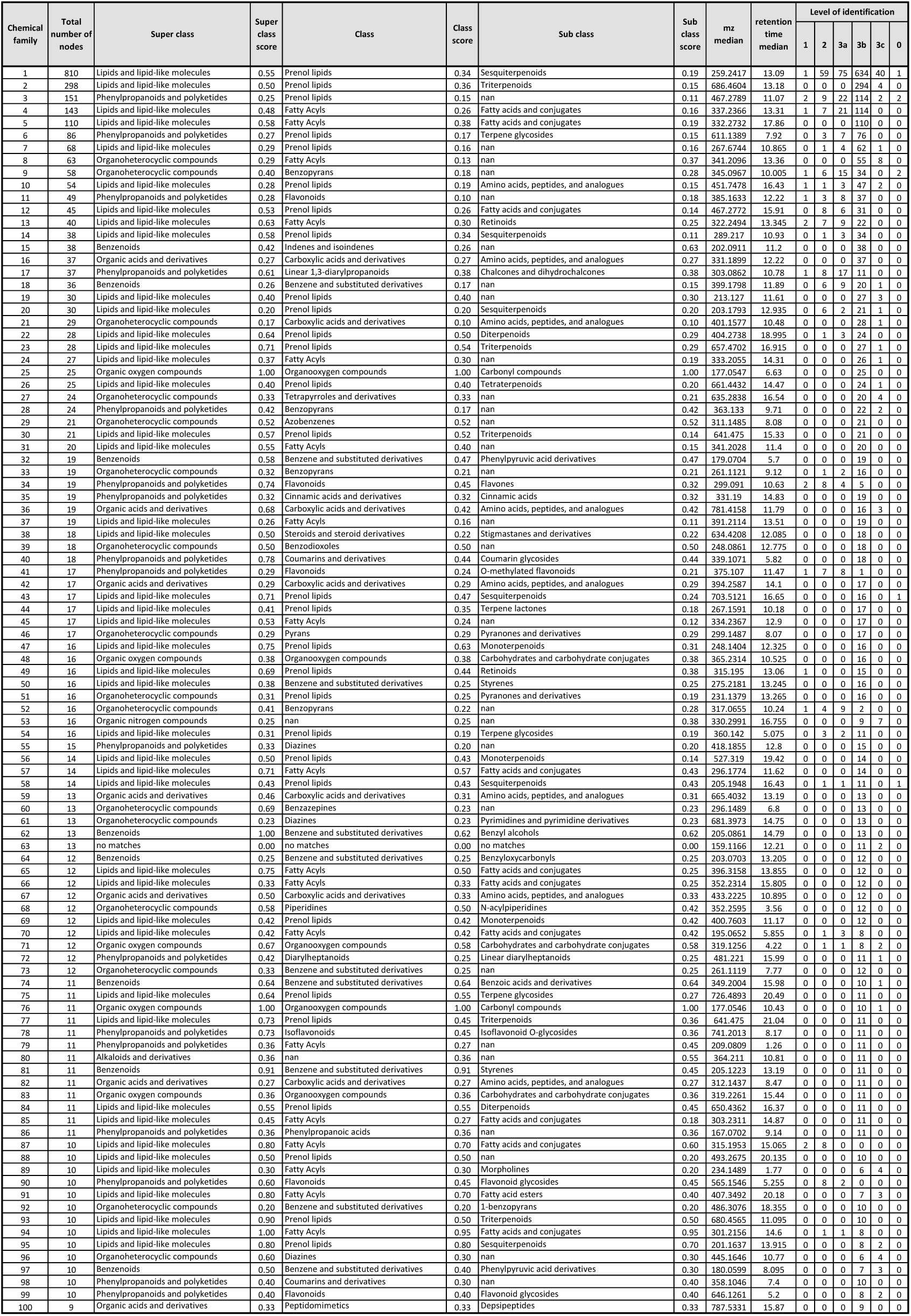
List of 100 largest chemical families derived from the molecular network of Myoporeae. Detailed information about the largest 100 chemical families generated by the molecular networking approach conducted in this study. Chemical family classification is predicted based on the applied dereplication pipeline. Further indicated are total node count and total numbers of different levels of chemical identification determined in this analysis.

**Table S10.**
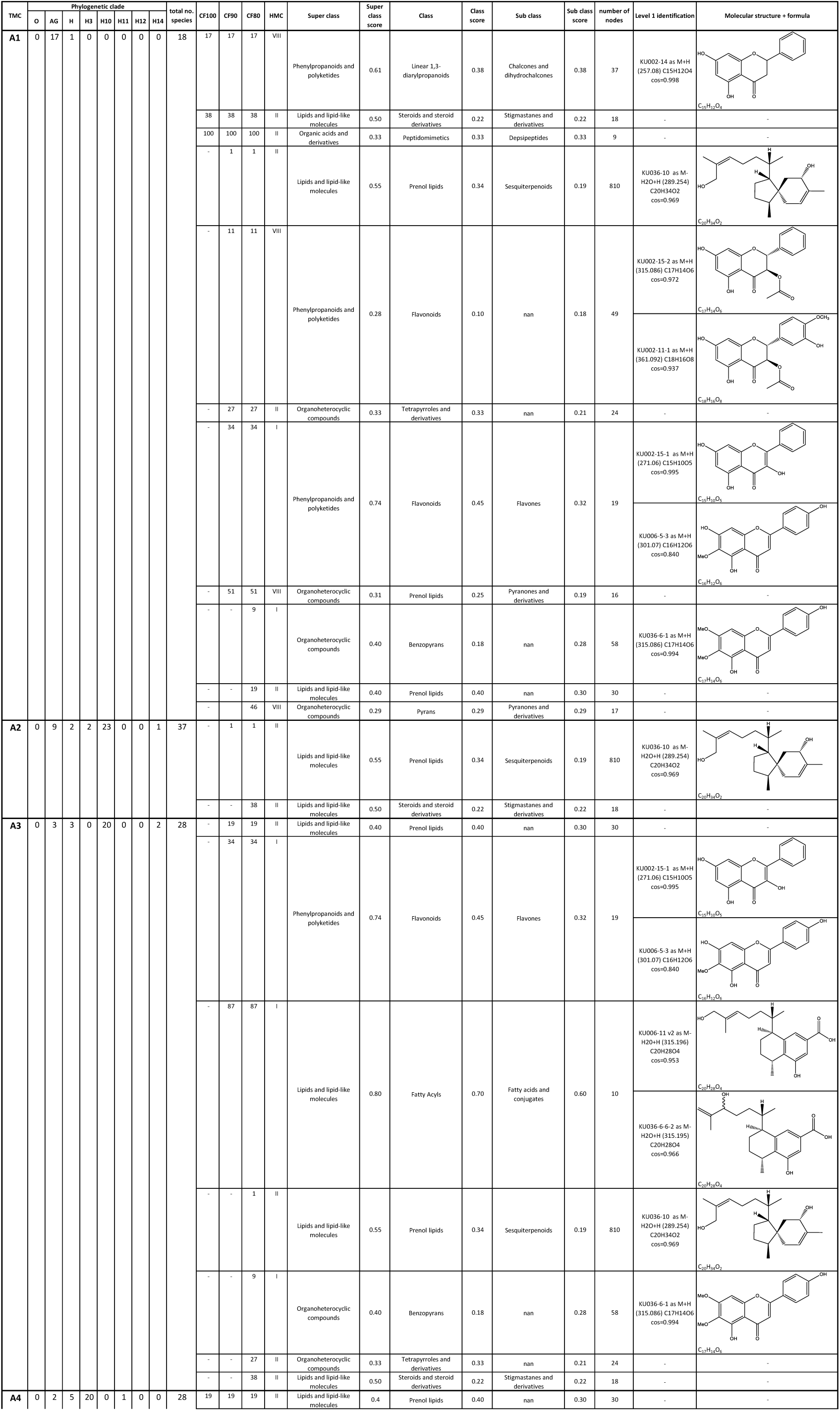
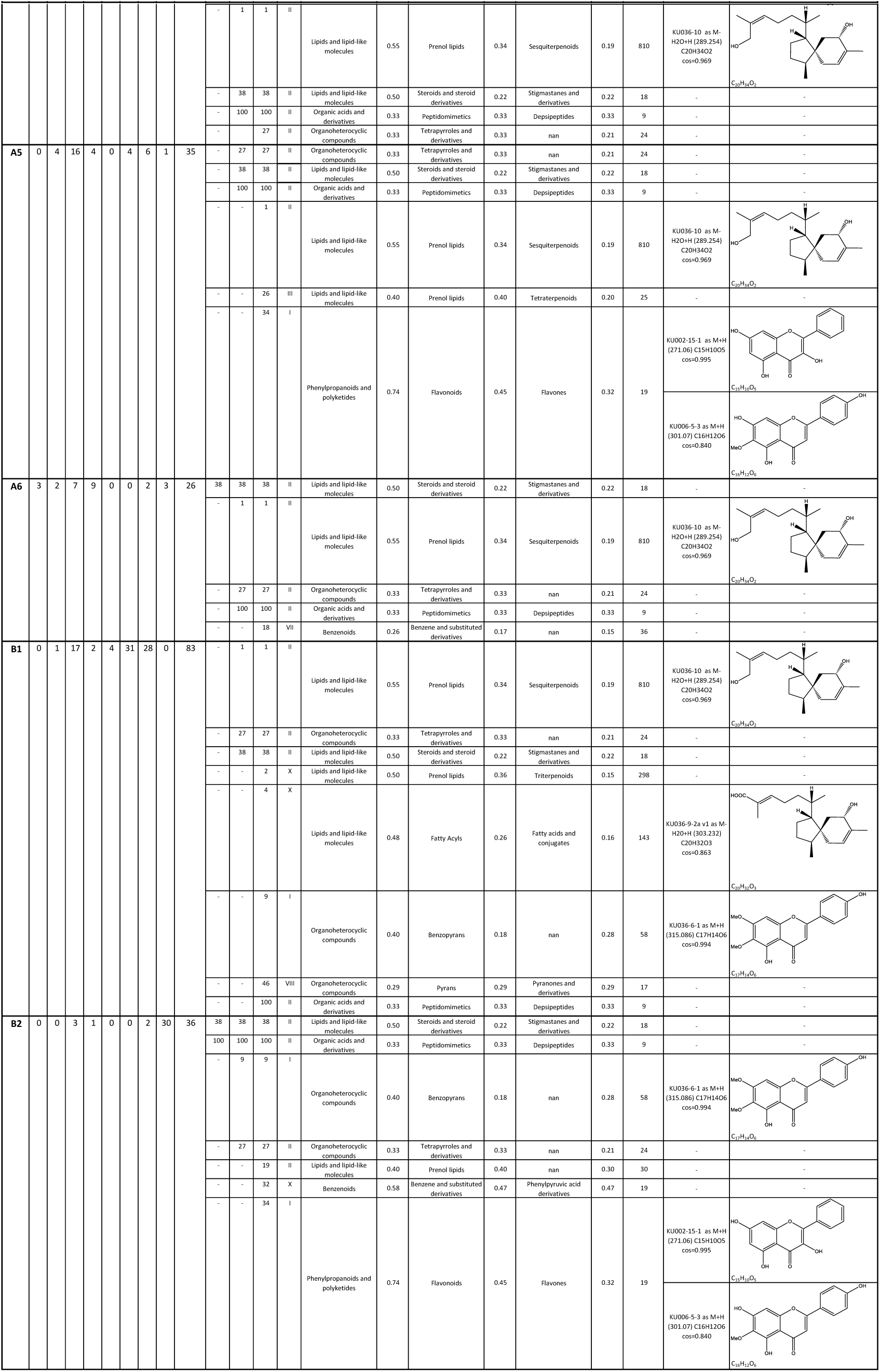
Phylogenetic and chemical information about the performed metabolic cluster analysis in Myoporaea. Comparison of the chemical content information extracted from determined tanglegram metabolic clusters (TMC). The underlying metabolic cluster analysis is based on the absence/presence of the 100 largest chemical families of the generated molecular network of Myoporeae. Number of specimens from major phylogenetic clades are shown with “O”: outgroup, “AG”: clade A to G, “H3”: clade H3, “H10”: clade H10, “H11”: clade H11, “H12”: clade H12, “H14”: clade H14 and “H”: remaining clade H associated taxa. Dominant chemical families are depicted for each TMC, with “CF100” indicating chemical families present in all species (100%) of the corresponding TMC, as well as “CF90” and “CF80”, depicting 90% and 80% abundancy of chemical families among the associated specimens, respectively. Different levels of predicted chemical classification and associated scores are shown. If present, level 1 identifications and their associated molecular structures are displayed.

**Table S11.**
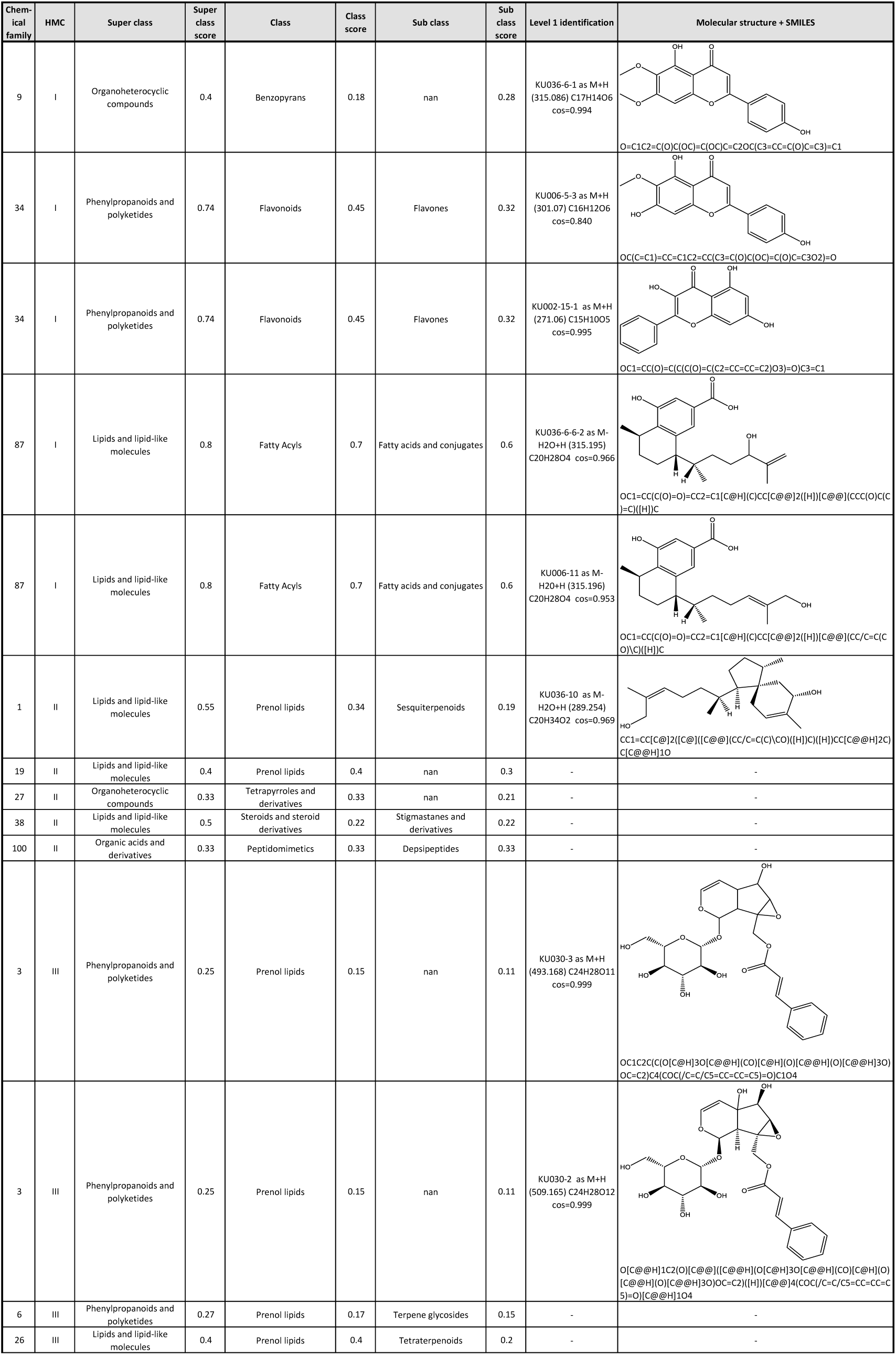
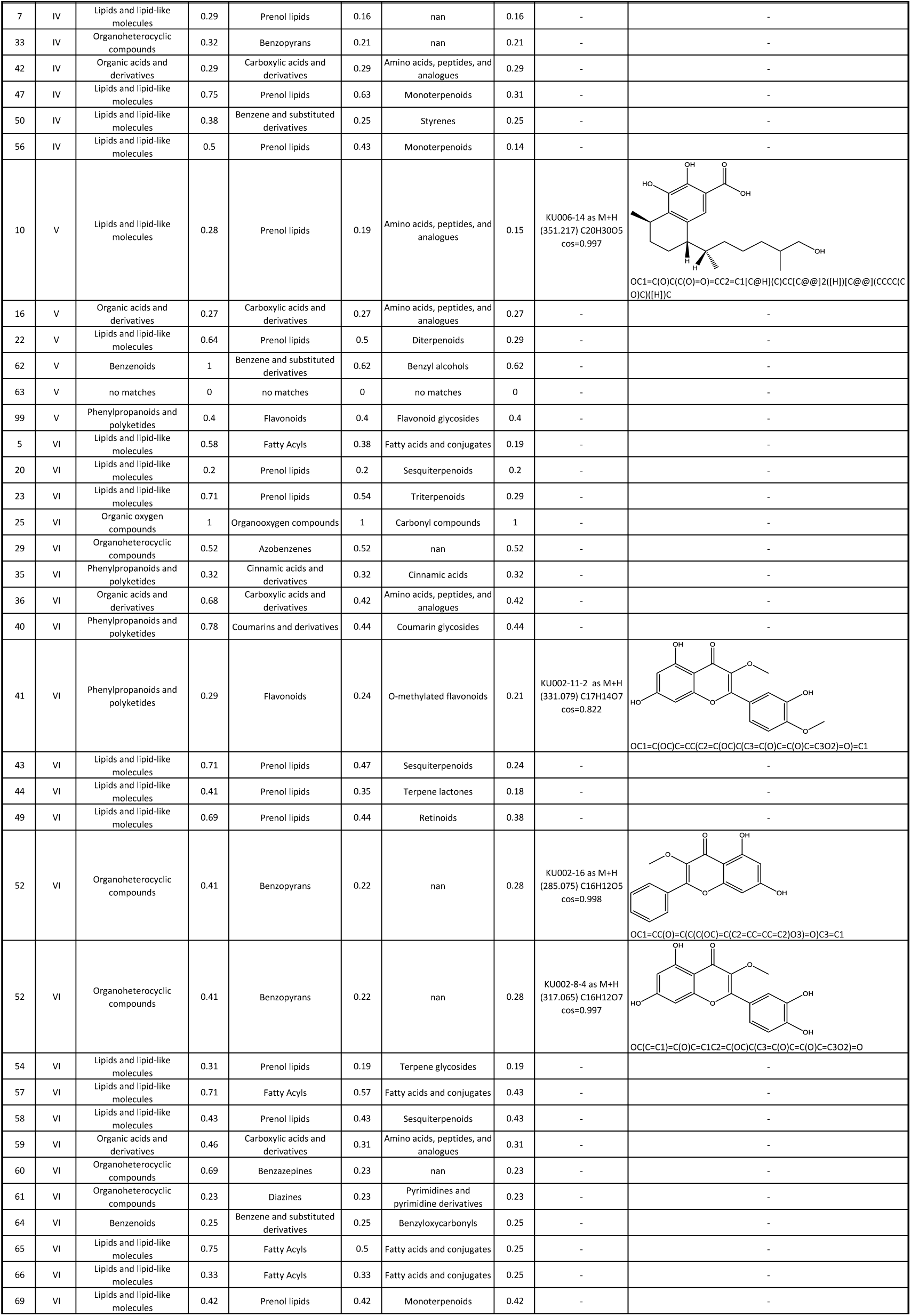
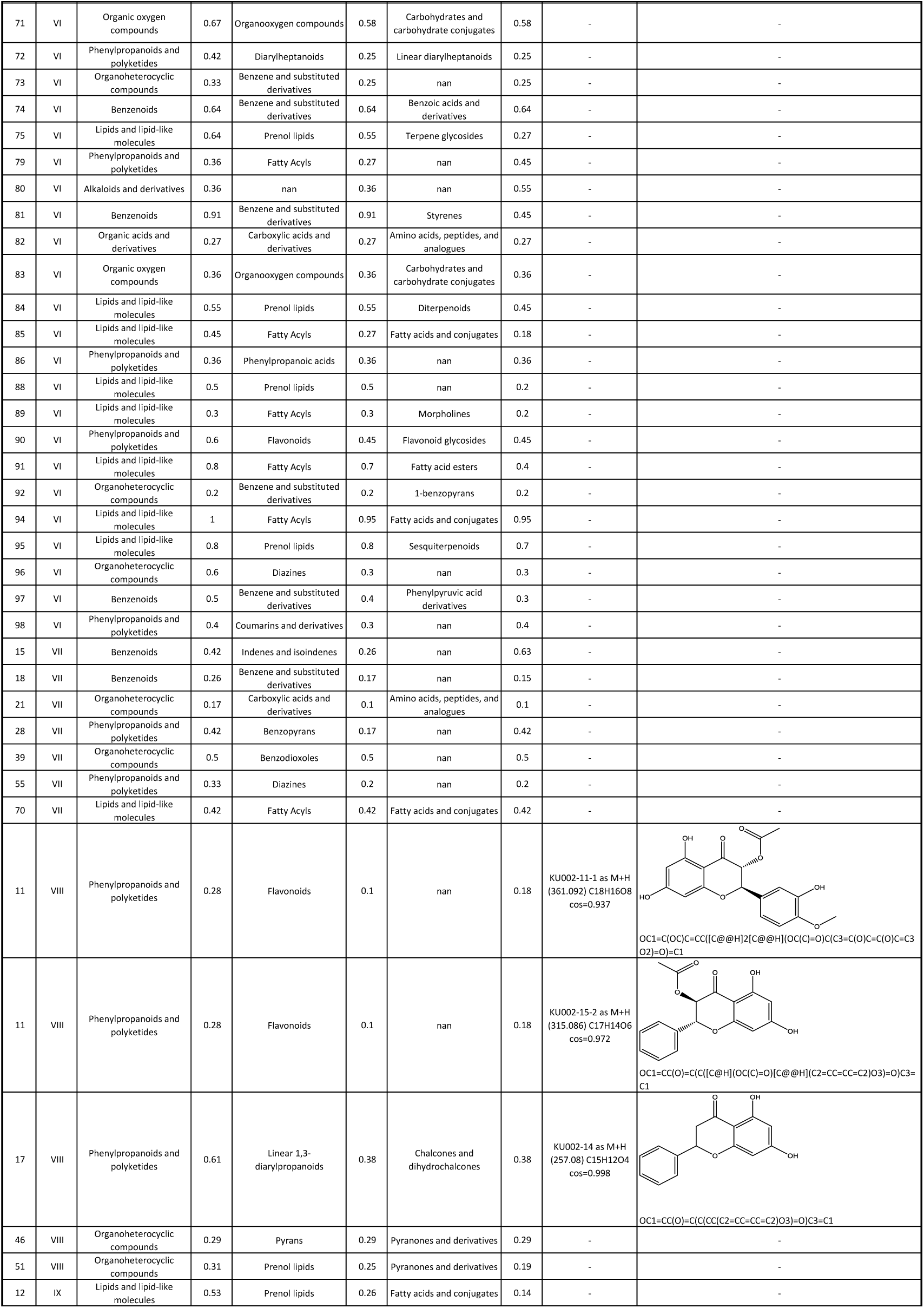
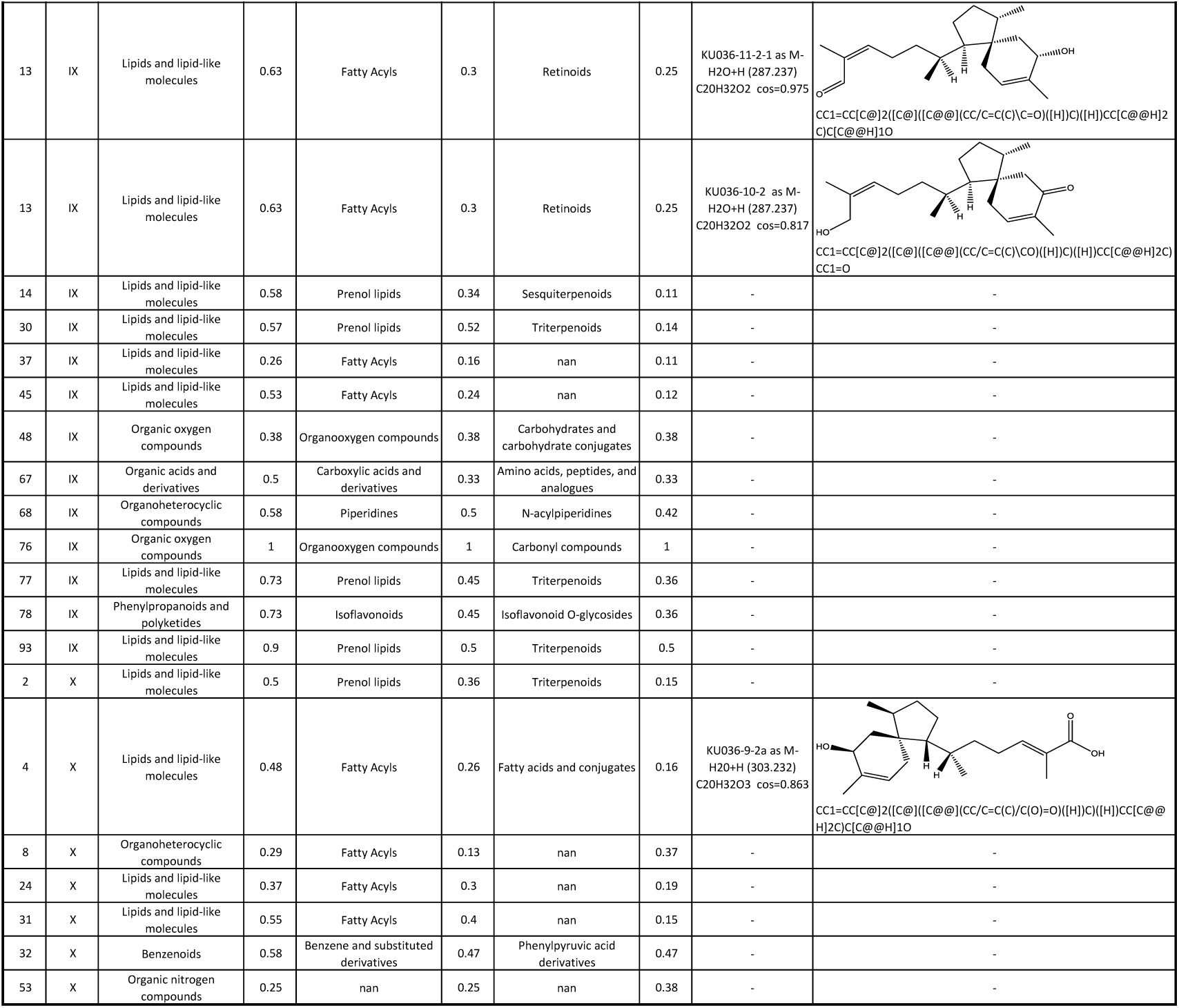
Detailed chemical information about heatmap metabolic clusters. Heatmap metabolic cluster (HMC) were generated using a categorical heatmap analysis based on the presence/absence of the 100 largest chemical families found in the molecular network of Myoporeae. Information about chemical identity is given based on the conducted dereplication pipeline. Further indicated are spectrometric and structural information about corresponding level 1 identification events.

**Table S12.**
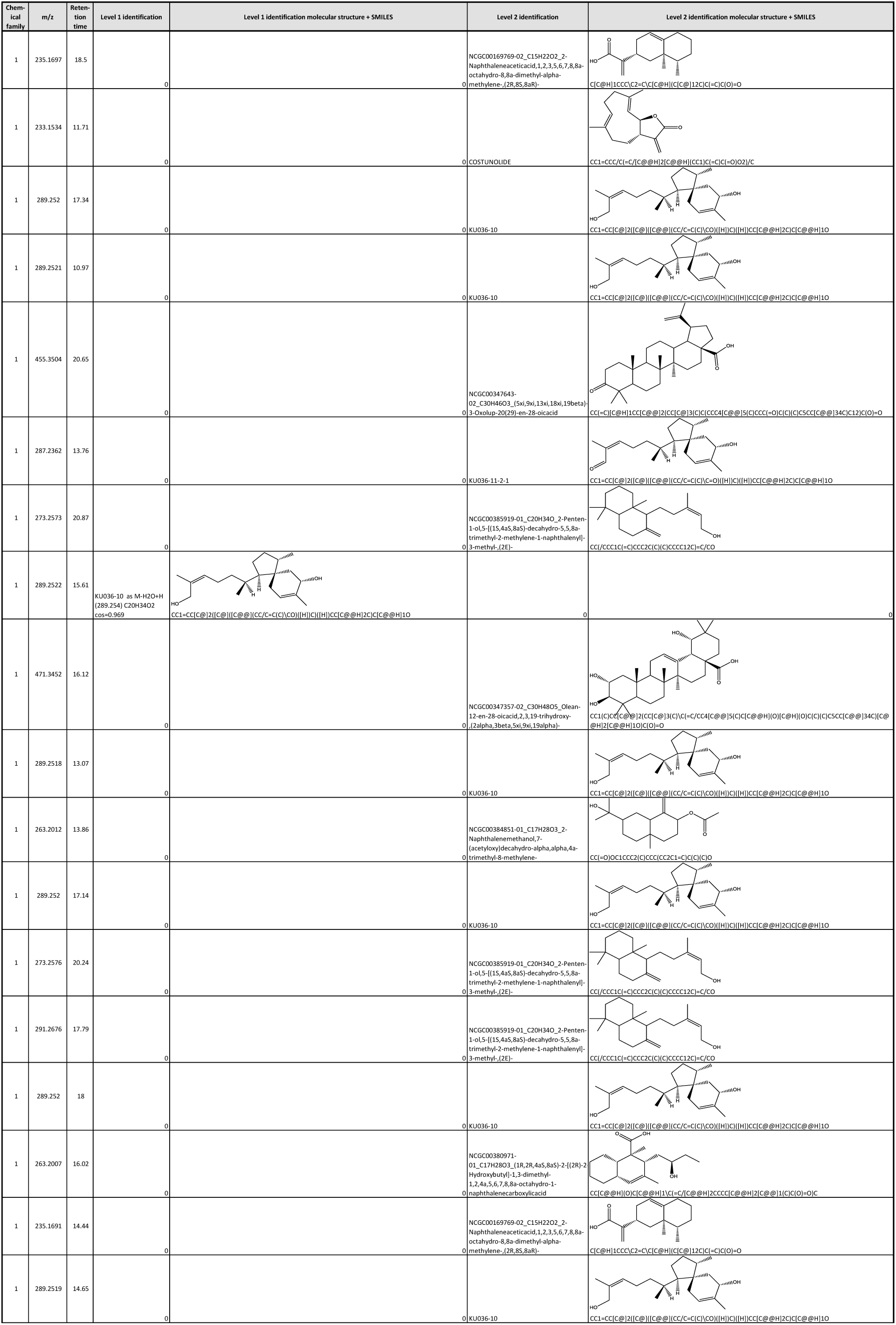
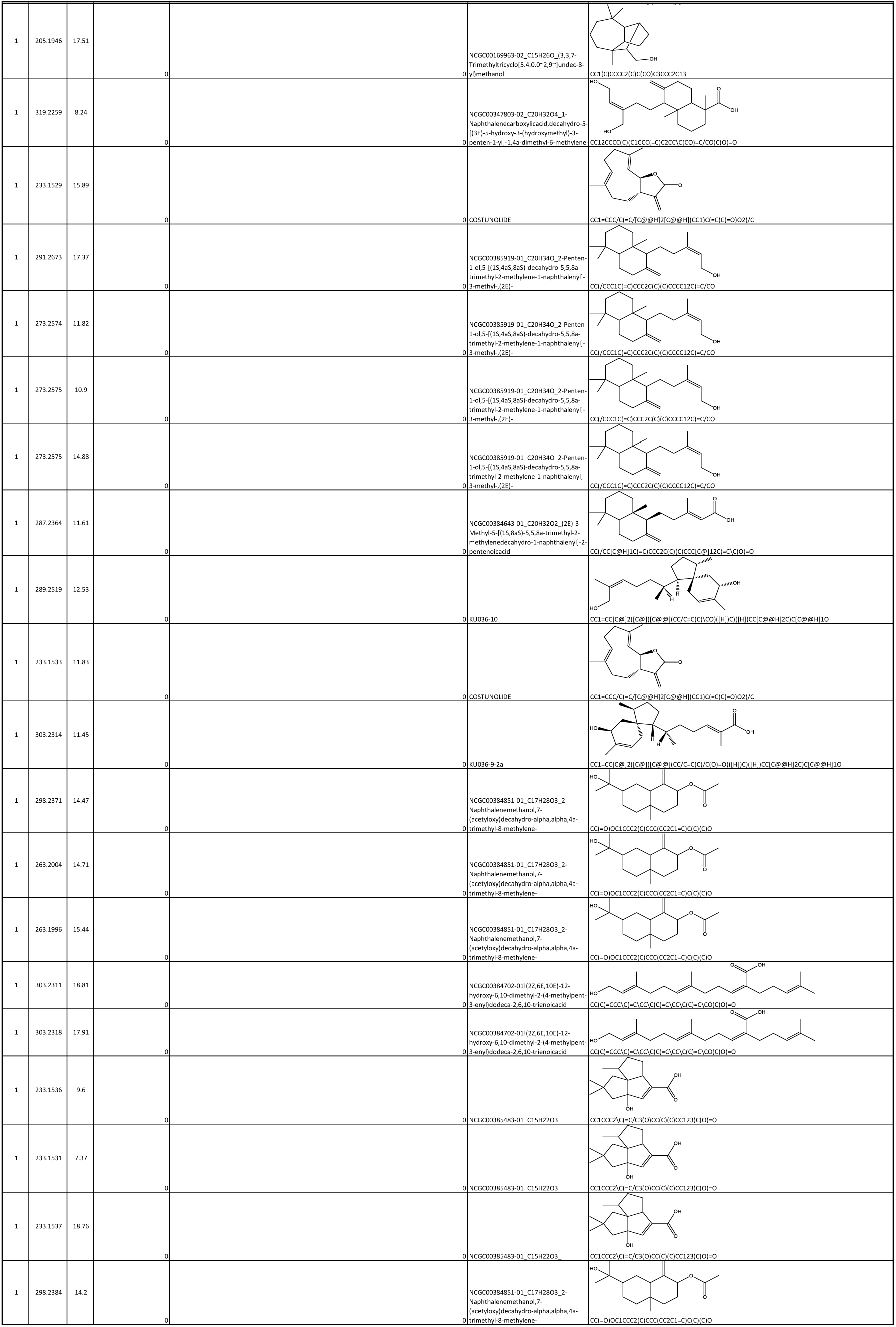
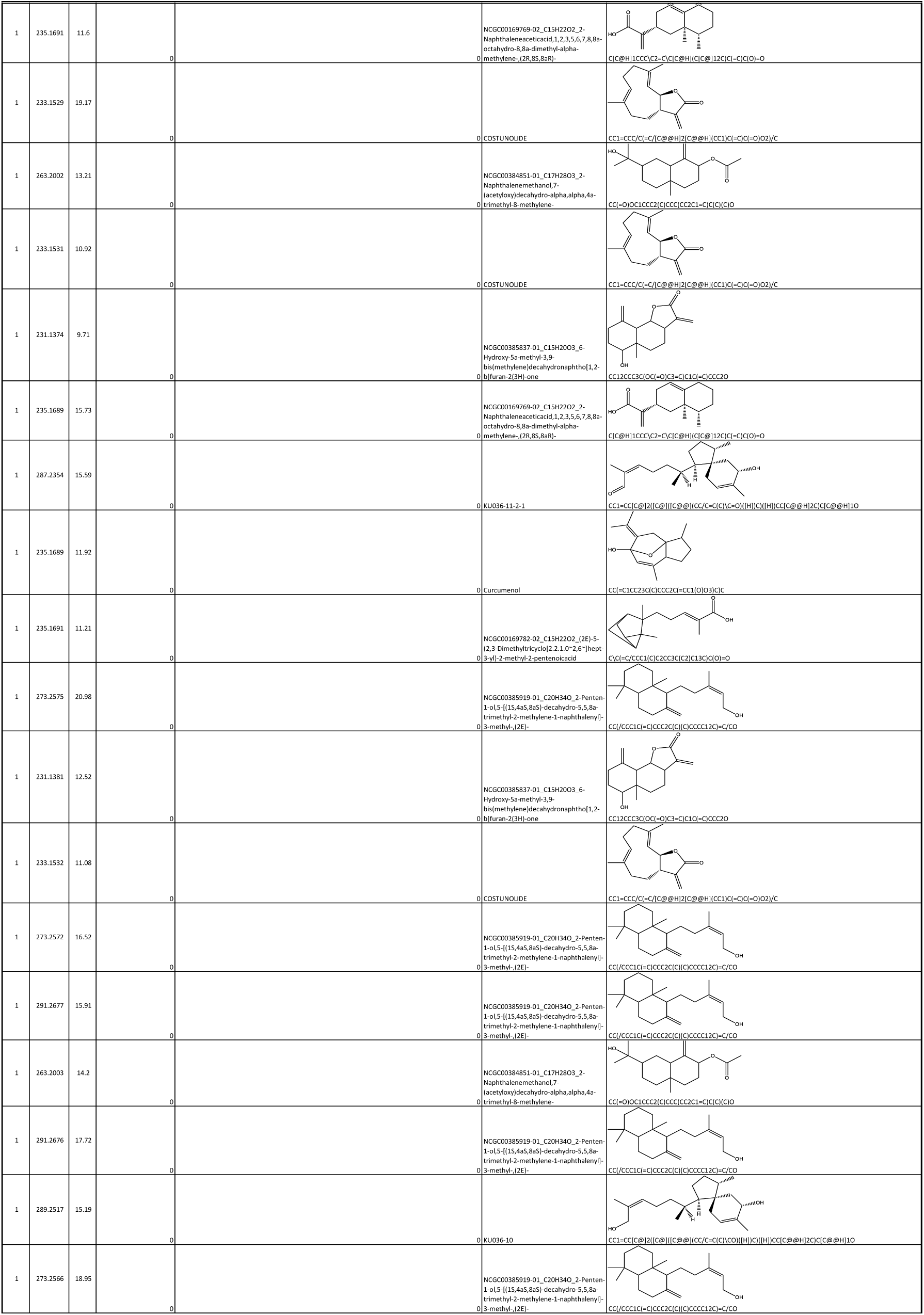
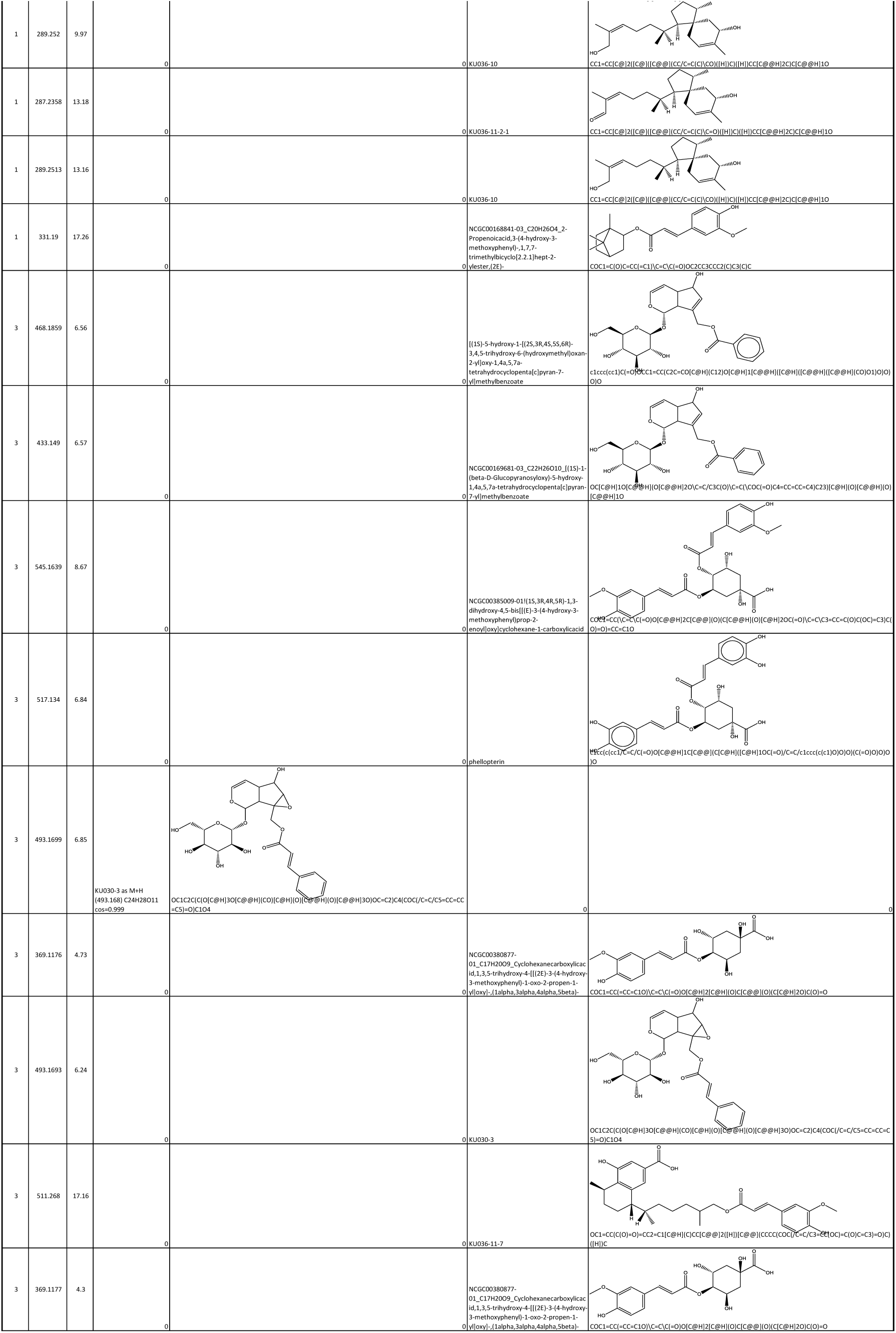
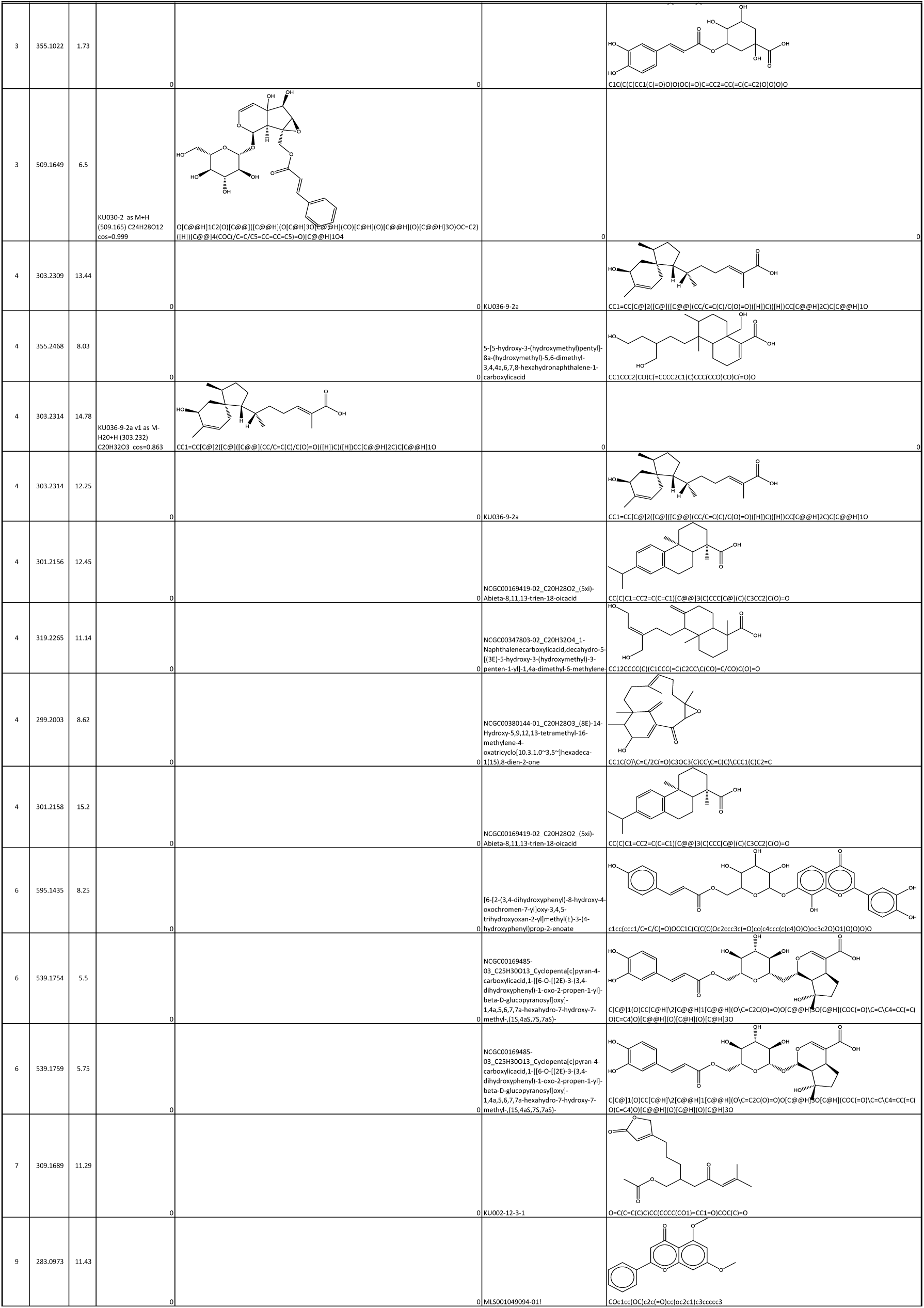
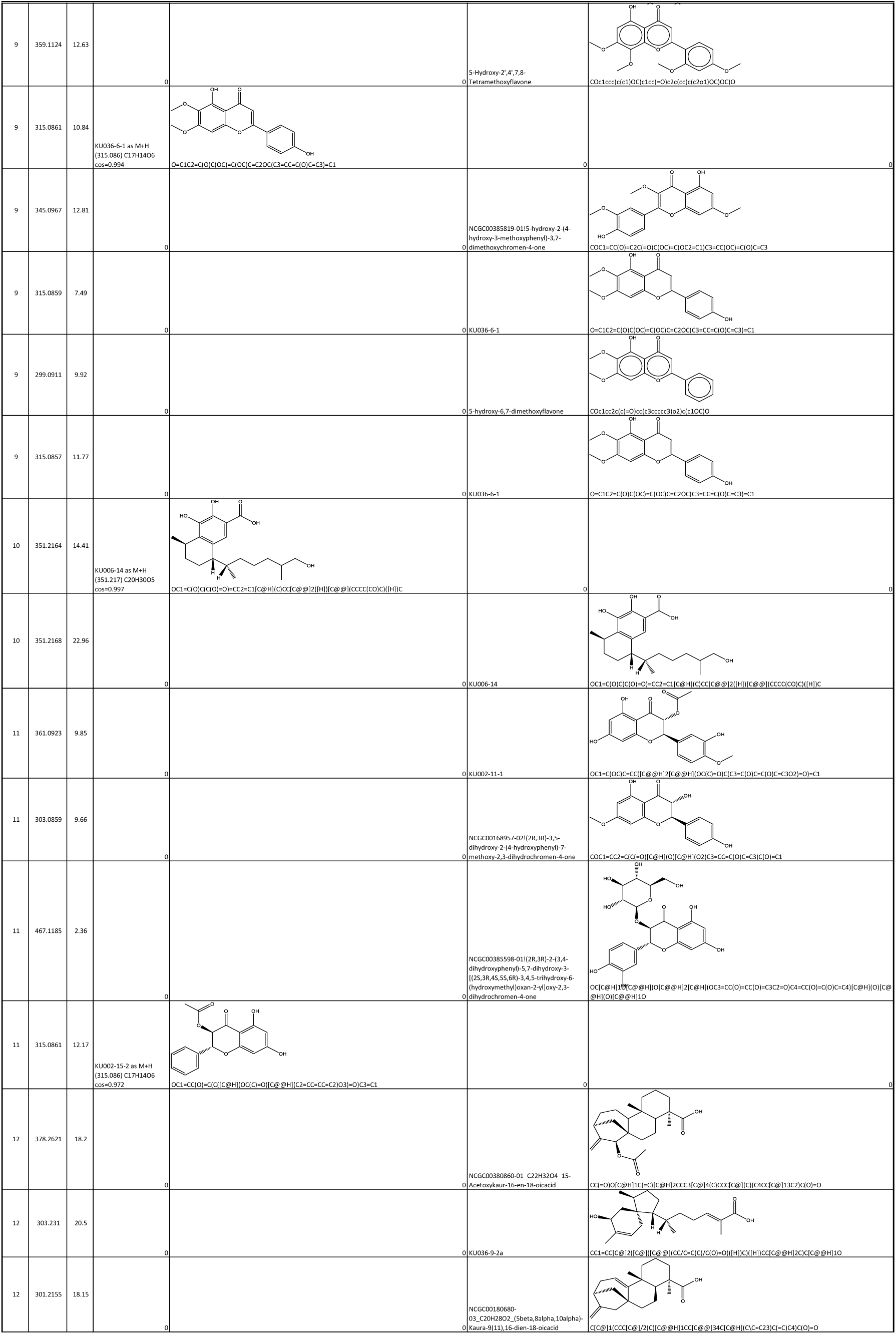
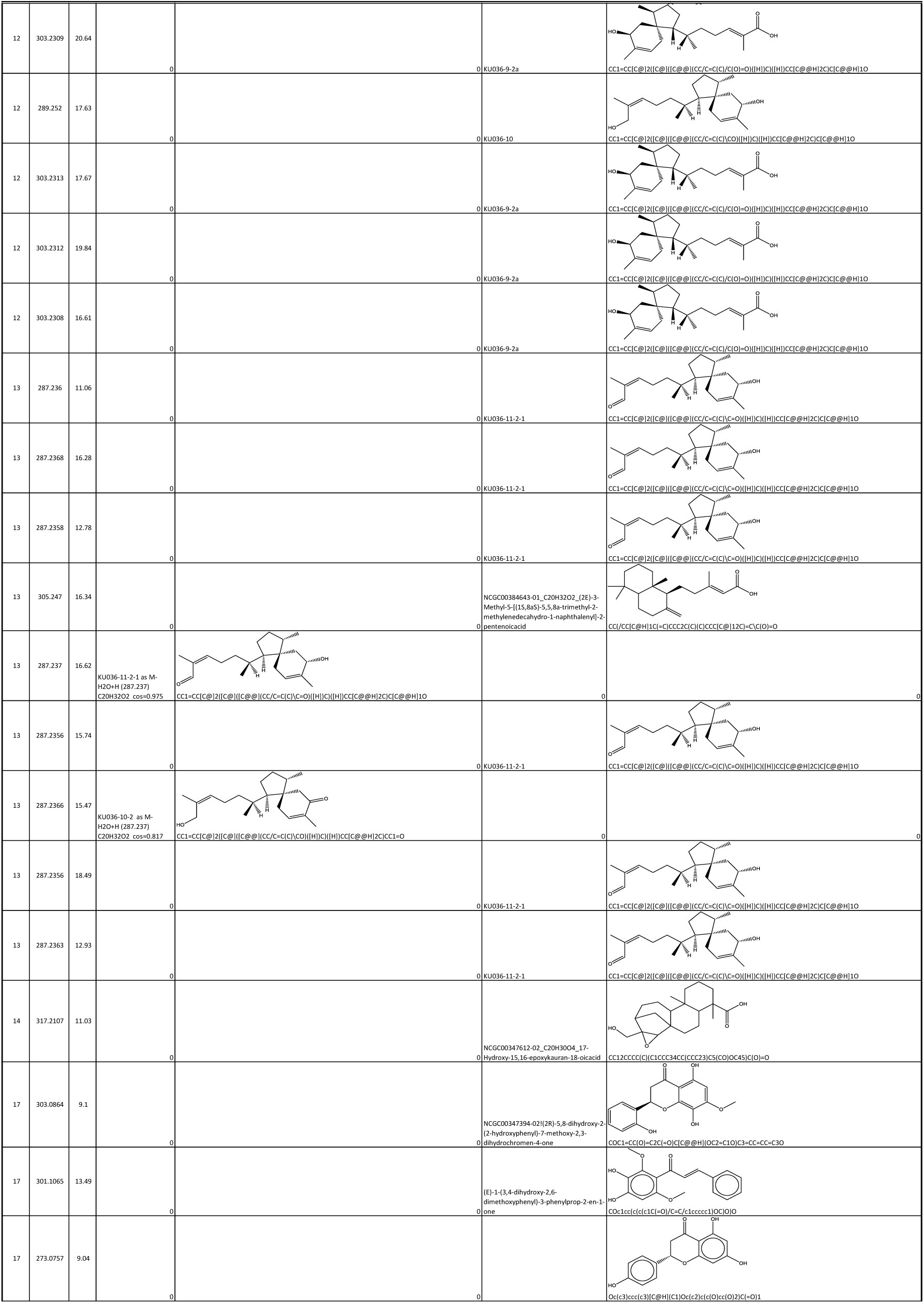
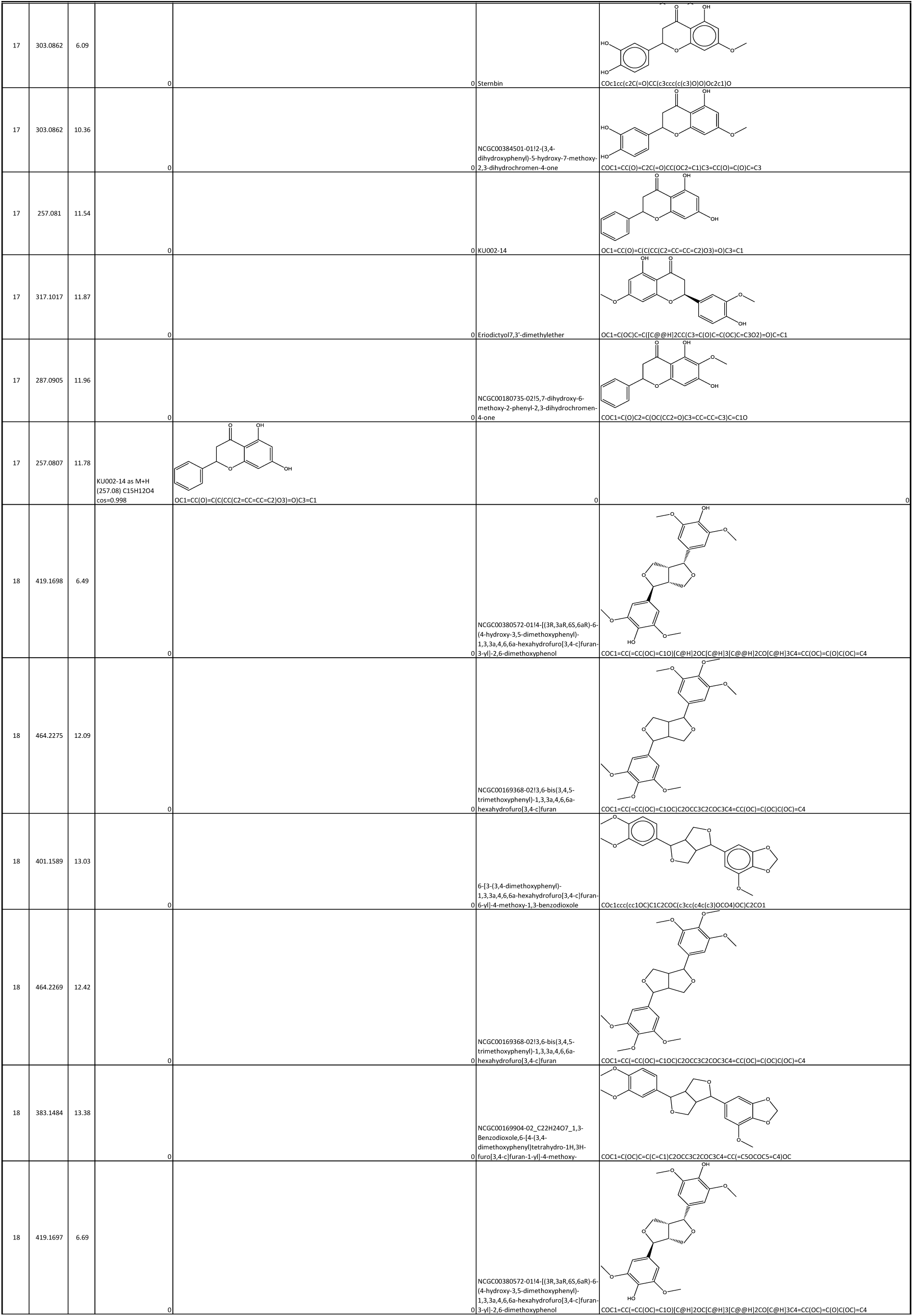
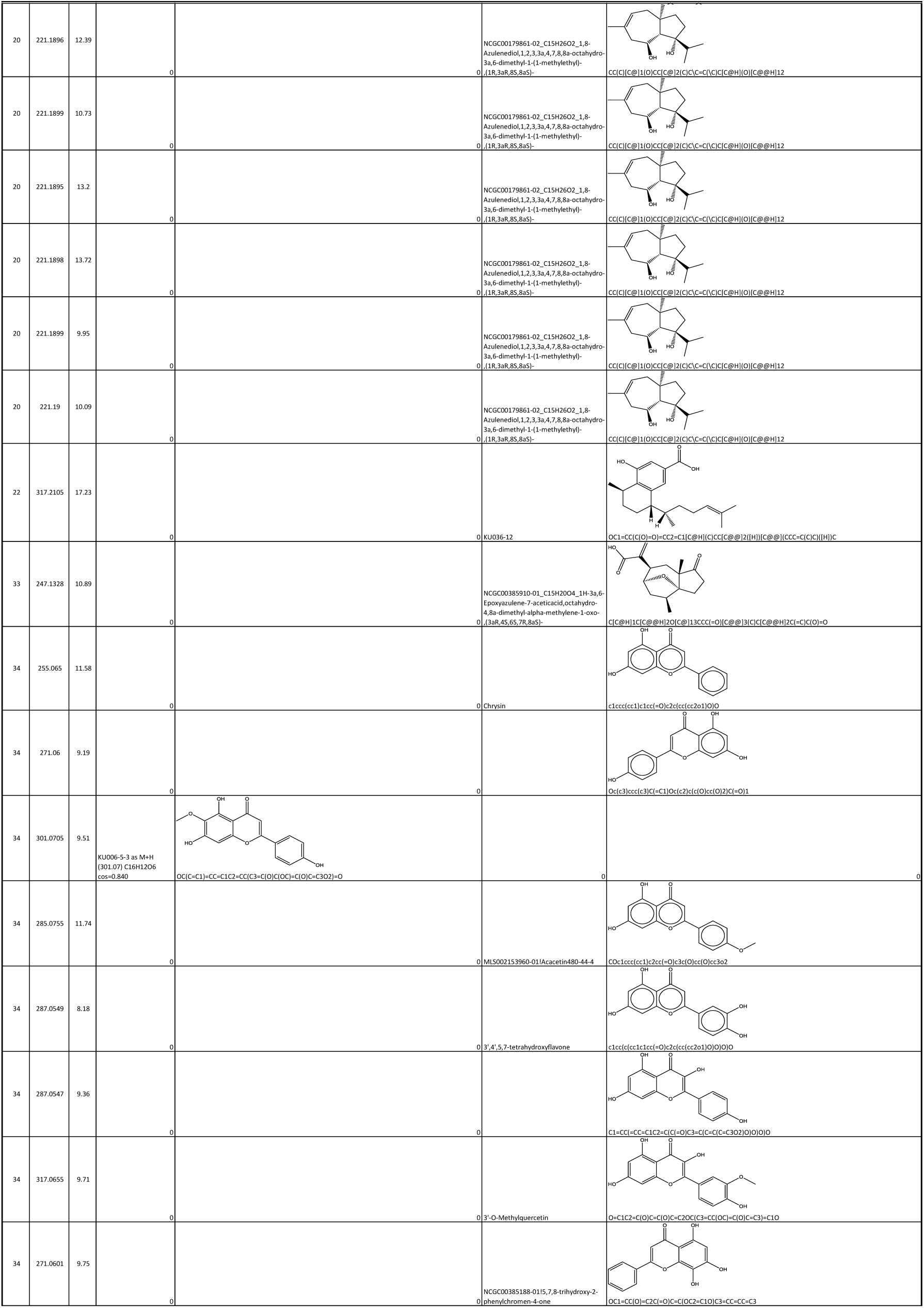
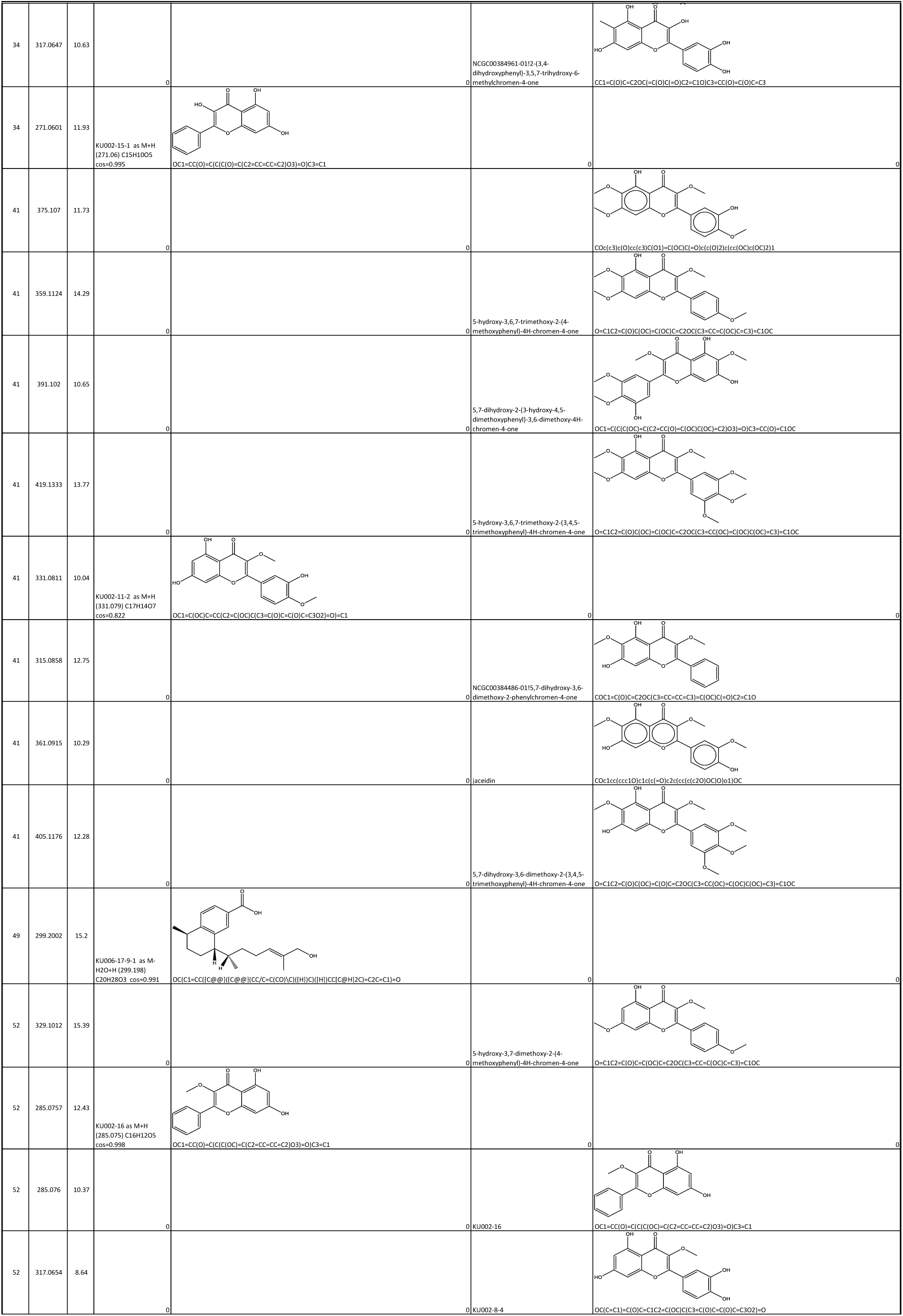
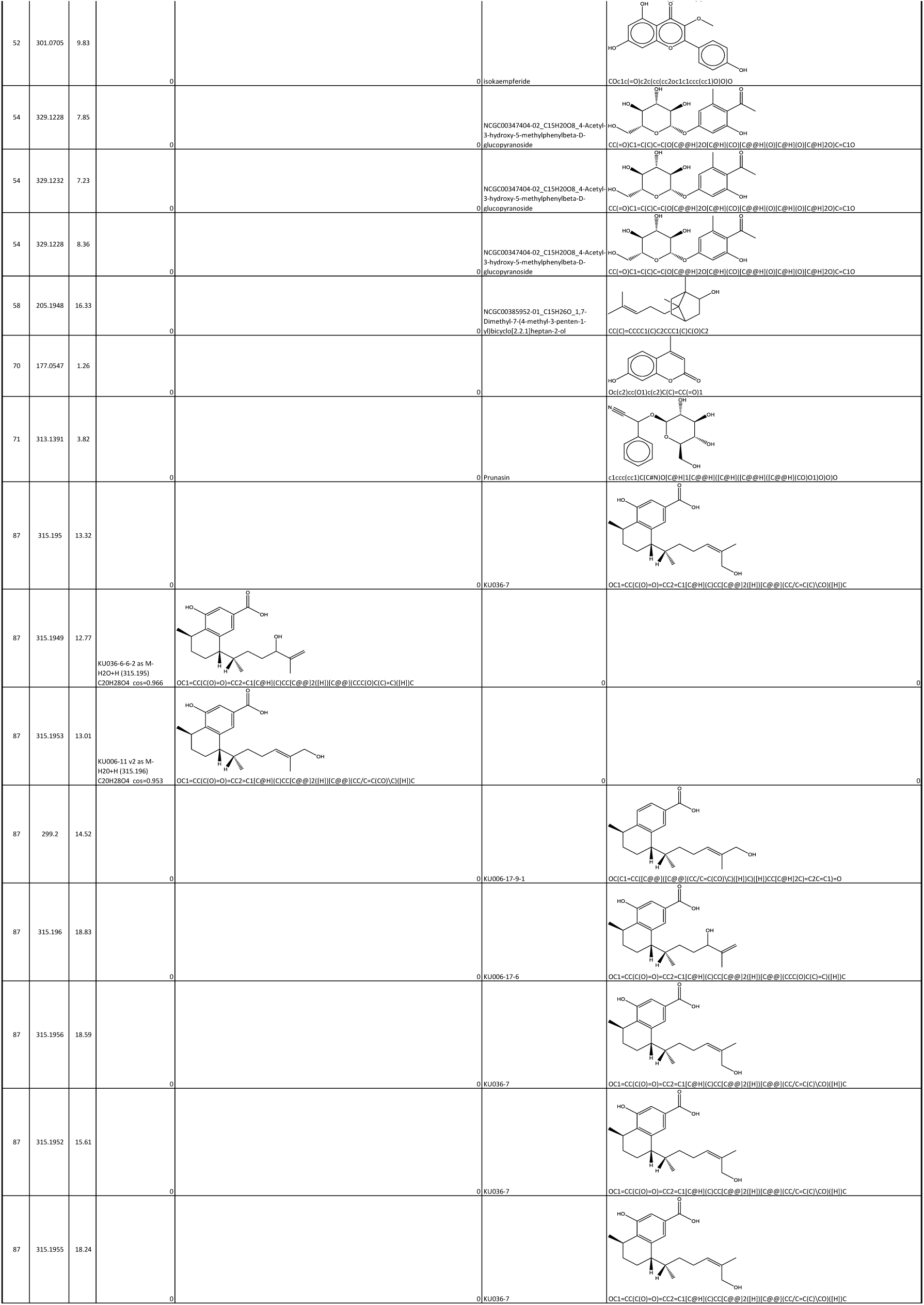
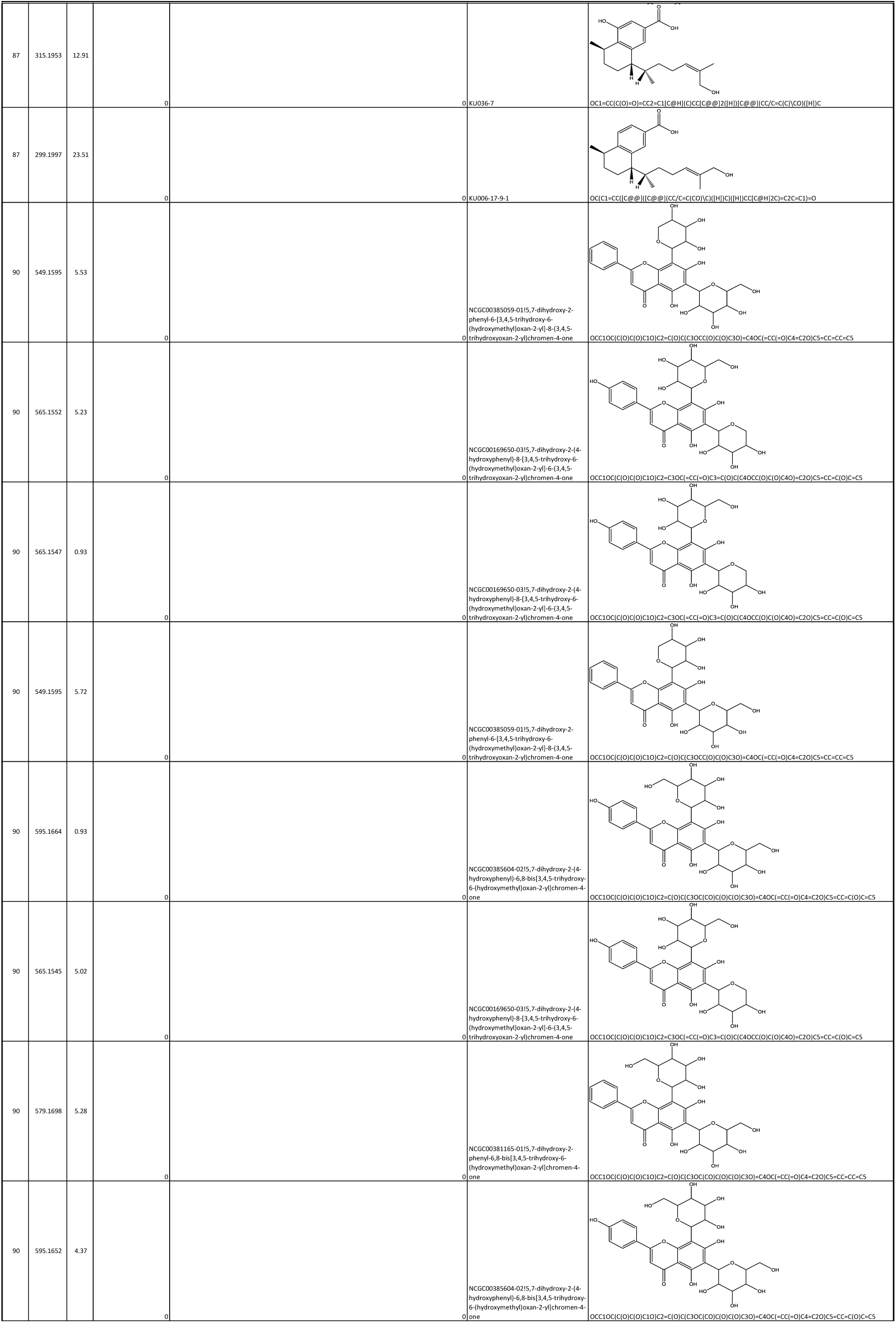
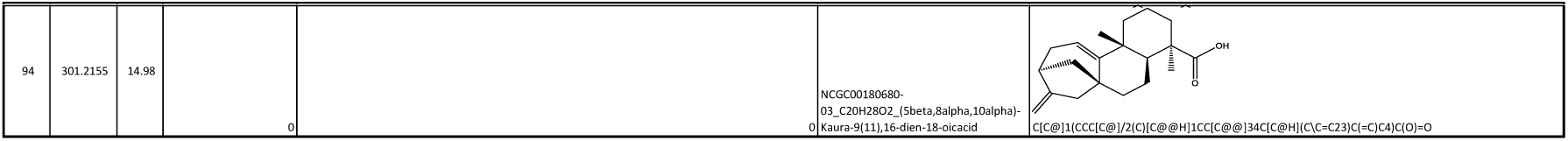
Identified metabolites within the 100 largest chemical families in Myoporeae. Detailed spectrometric and structural information about level 1 and 2 identification events found within the largest 100 chemical families from the molecular network of Myoporeae.

**Table S13.** Dereplication of serrulatane and viscidane-related chemical families. List of 30 chemical families from Myoporeae that have been dereplicated as serrulatane or viscidane related. Chemical family classification is predicted based on the applied dereplication pipeline. Further indicated are total node count and total numbers of different levels of chemical identification determined in this analysis as well as mass and retention time median of each chemical family.

| Chemical family | Total number of nodes | Super class | Super class score | Class | Class score | Sub class | Sub class score | Diterpenoid class | m/z median | Retention time median | Level of identification |  |  |  |  |  |
| --- | --- | --- | --- | --- | --- | --- | --- | --- | --- | --- | --- | --- | --- | --- | --- | --- |
|  |  |  |  |  |  |  |  |  |  |  | 1 | 2 | 3a | 3b | 3c | 0 |
| 1 | 810 | Lipids and lipid-like molecules | 0.55 | Prenol lipids | 0.34 | Sesquiterpenoids | 0.19 | Viscidane | 259.2417 | 13.09 | 1 | 59 | 75 | 634 | 40 | 1 |
| 3 | 151 | Phenylpropanoids and polyketides | 0.25 | Prenol lipids | 0.15 | nan | 0.11 | Serrulatane | 467.2789 | 11.07 | 2 | 9 | 22 | 114 | 2 | 2 |
| 4 | 143 | Lipids and lipid-like molecules | 0.48 | Fatty Acyls | 0.26 | Fatty acids and conjugates | 0.16 | Viscidane | 337.2366 | 13.31 | 1 | 7 | 21 | 114 | 0 | 0 |
| 8 | 63 | Organoheterocyclic compounds | 0.29 | Fatty Acyls | 0.13 | nan | 0.37 | Serrulatane | 341.2096 | 13.36 | 0 | 0 | 0 | 55 | 8 | 0 |
| 10 | 54 | Lipids and lipid-like molecules | 0.28 | Prenol lipids | 0.19 | Amino acids, peptides, and analogues | 0.15 | Serrulatane | 451.74775 | 16.43 | 1 | 1 | 3 | 47 | 2 | 0 |
| 12 | 45 | Lipids and lipid-like molecules | 0.53 | Prenol lipids | 0.26 | Fatty acids and conjugates | 0.14 | Viscidane | 467.2772 | 15.91 | 0 | 8 | 6 | 31 | 0 | 0 |
| 13 | 40 | Lipids and lipid-like molecules | 0.63 | Fatty Acyls | 0.30 | Retinoids | 0.25 | Viscidane | 322.24935 | 13.345 | 2 | 7 | 9 | 22 | 0 | 0 |
| 22 | 28 | Lipids and lipid-like molecules | 0.64 | Prenol lipids | 0.50 | Diterpenoids | 0.29 | Serrulatane | 404.2738 | 18.995 | 0 | 1 | 3 | 24 | 0 | 0 |
| 24 | 27 | Lipids and lipid-like molecules | 0.37 | Fatty Acyls | 0.30 | nan | 0.19 | Serrulatane | 333.2055 | 14.31 | 0 | 0 | 0 | 26 | 1 | 0 |
| 42 | 17 | Organic acids and derivatives | 0.29 | Carboxylic acids and derivatives | 0.29 | Amino acids, peptides, and analogues | 0.29 | Serrulatane | 394.2587 | 14.1 | 0 | 0 | 0 | 17 | 0 | 0 |
| 49 | 16 | Lipids and lipid-like molecules | 0.69 | Prenol lipids | 0.44 | Retinoids | 0.38 | Serrulatane | 315.19495 | 13.06 | 1 | 0 | 0 | 15 | 0 | 0 |
| 74 | 11 | Benzenoids | 0.64 | Benzene and substituted derivatives | 0.64 | Benzoic acids and derivatives | 0.64 | Serrulatane | 349.2004 | 15.98 | 0 | 0 | 0 | 10 | 1 | 0 |
| 87 | 10 | Lipids and lipid-like molecules | 0.80 | Fatty Acyls | 0.70 | Fatty acids and conjugates | 0.60 | Serrulatane | 315.19525 | 15.065 | 2 | 8 | 0 | 0 | 0 | 0 |
| 101 | 8 | Lipids and lipid-like molecules | 0.38 | Prenol lipids | 0.25 | Retinoids | 0.13 | Serrulatane | 355.20085 | 12.555 | 0 | 0 | 0 | 8 | 0 | 0 |
| 102 | 8 | Lipids and lipid-like molecules | 0.88 | Prenol lipids | 0.63 | Diterpenoids | 0.38 | Serrulatane | 294.2263 | 15.86 | 0 | 1 | 2 | 5 | 0 | 0 |
| 103 | 7 | Lipids and lipid-like molecules | 0.86 | Prenol lipids | 0.43 | Fatty acids and conjugates | 0.43 | Viscidane | 305.2463 | 12.63 | 0 | 1 | 3 | 3 | 0 | 0 |
| 104 | 6 | Lipids and lipid-like molecules | 0.83 | Fatty Acyls | 0.50 | Retinoids | 0.33 | Viscidane | 312.2358 | 10.96 | 0 | 2 | 1 | 3 | 0 | 0 |
| 105 | 5 | Lipids and lipid-like molecules | 0.60 | Fatty Acyls | 0.40 | Lineolic acids and derivatives | 0.40 | Viscidane | 291.2314 | 16.26 | 0 | 1 | 1 | 3 | 0 | 0 |
| 106 | 5 | Phenylpropanoids and polyketides | 0.20 | Coumarins and derivatives | 0.20 | nan | 0.20 | Serrulatane | 333.2056 | 11.02 | 0 | 0 | 0 | 5 | 0 | 0 |
| 107 | 4 | Phenylpropanoids and polyketides | 0.50 | Stilbenes | 0.25 | nan | 0.50 | Serrulatane | 322.18485 | 14.155 | 1 | 0 | 2 | 1 | 0 | 0 |
| 108 | 4 | Lipids and lipid-like molecules | 1.00 | Prenol lipids | 1.00 | Diterpenoids | 0.50 | Viscidane | 289.2523 | 14.595 | 1 | 1 | 2 | 0 | 0 | 0 |
| 109 | 4 | Phenylpropanoids and polyketides | 0.75 | Coumarins and derivatives | 0.50 | nan | 0.50 | Serrulatane | 273.1484 | 13.32 | 0 | 2 | 2 | 0 | 0 | 0 |
| 110 | 3 | Benzenoids | 0.33 | Benzene and substituted derivatives | 0.33 | Monoterpenoids | 0.33 | Serrulatane | 392.2422 | 15.62 | 0 | 0 | 0 | 3 | 0 | 0 |
| 111 | 2 | Lipids and lipid-like molecules | 1.00 | Fatty Acyls | 1.00 | Fatty acids and conjugates | 1.00 | Serrulatane | 310.2213 | 11.02 | 0 | 0 | 0 | 2 | 0 | 0 |
| 112 | 2 | Lipids and lipid-like molecules | 1.00 | Fatty Acyls | 1.00 | Fatty acids and conjugates | 1.00 | Viscidane | 312.23675 | 11.45 | 0 | 1 | 1 | 0 | 0 | 0 |
| 113 | 2 | no matches | 0.00 | no matches | 0.00 | no matches | 0.00 | Serrulatane | 333.20545 | 15.59 | 0 | 0 | 0 | 2 | 0 | 0 |
| 114 | 2 | Lipids and lipid-like molecules | 0.50 | Prenol lipids | 0.50 | Diterpenoids | 0.50 | Serrulatane | 342.211 | 11.64 | 0 | 0 | 0 | 2 | 0 | 0 |
| 115 | 2 | no matches | 0.00 | no matches | 0.00 | no matches | 0.00 | Serrulatane | 369.71415 | 15.04 | 1 | 0 | 1 | 0 | 0 | 0 |
| 116 | 2 | Organic acids and derivatives | 0.50 | Furofurans | 0.50 | nan | 0.50 | Serrulatane | 343.219 | 7.7 | 0 | 0 | 0 | 2 | 0 | 0 |
| 117 | 2 | no matches | 0.00 | no matches | 0.00 | no matches | 0.00 | Serrulatane | 365.1956 | 14.735 | 0 | 0 | 0 | 0 | 2 | 0 |

**Table S14.**
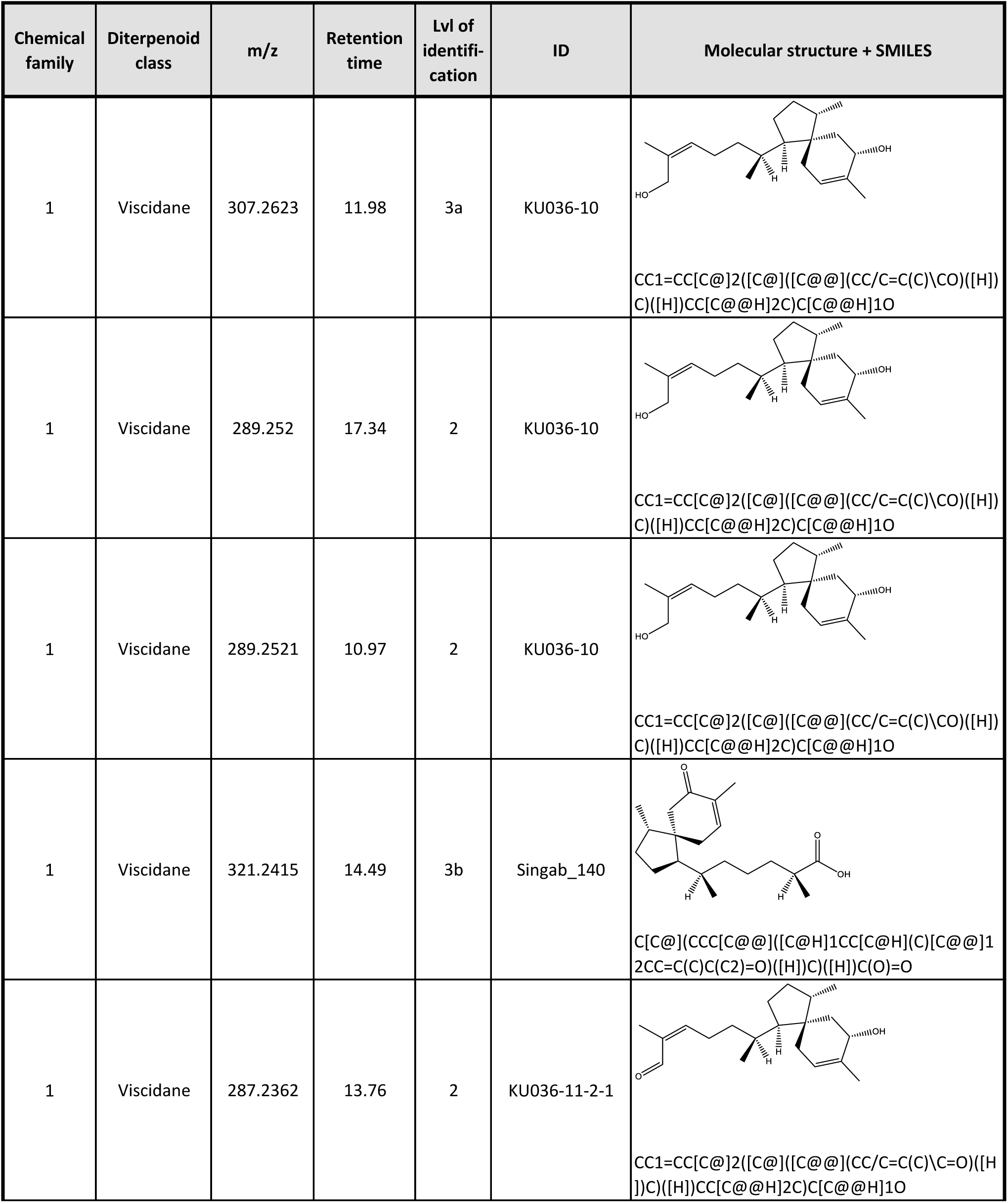
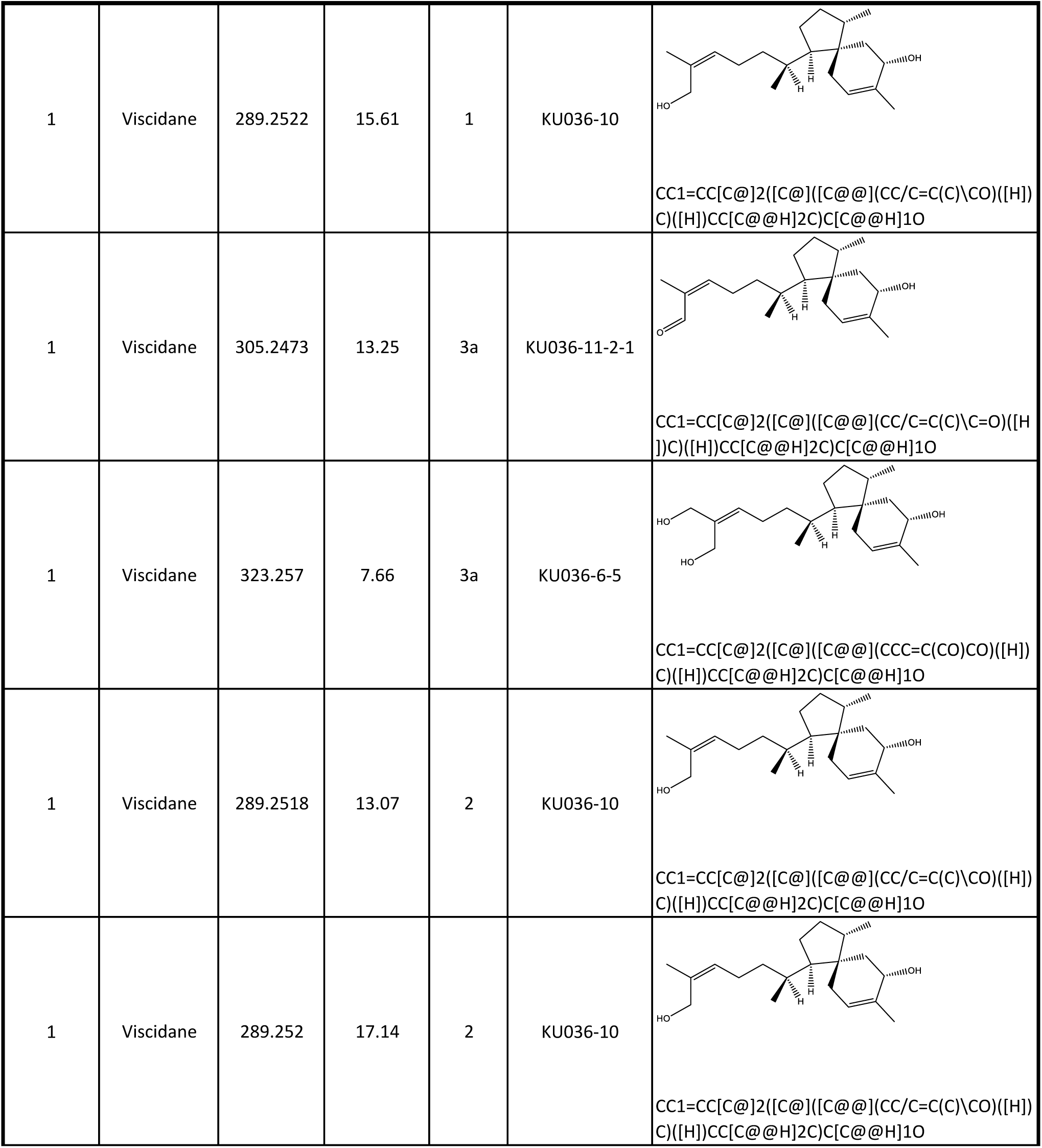
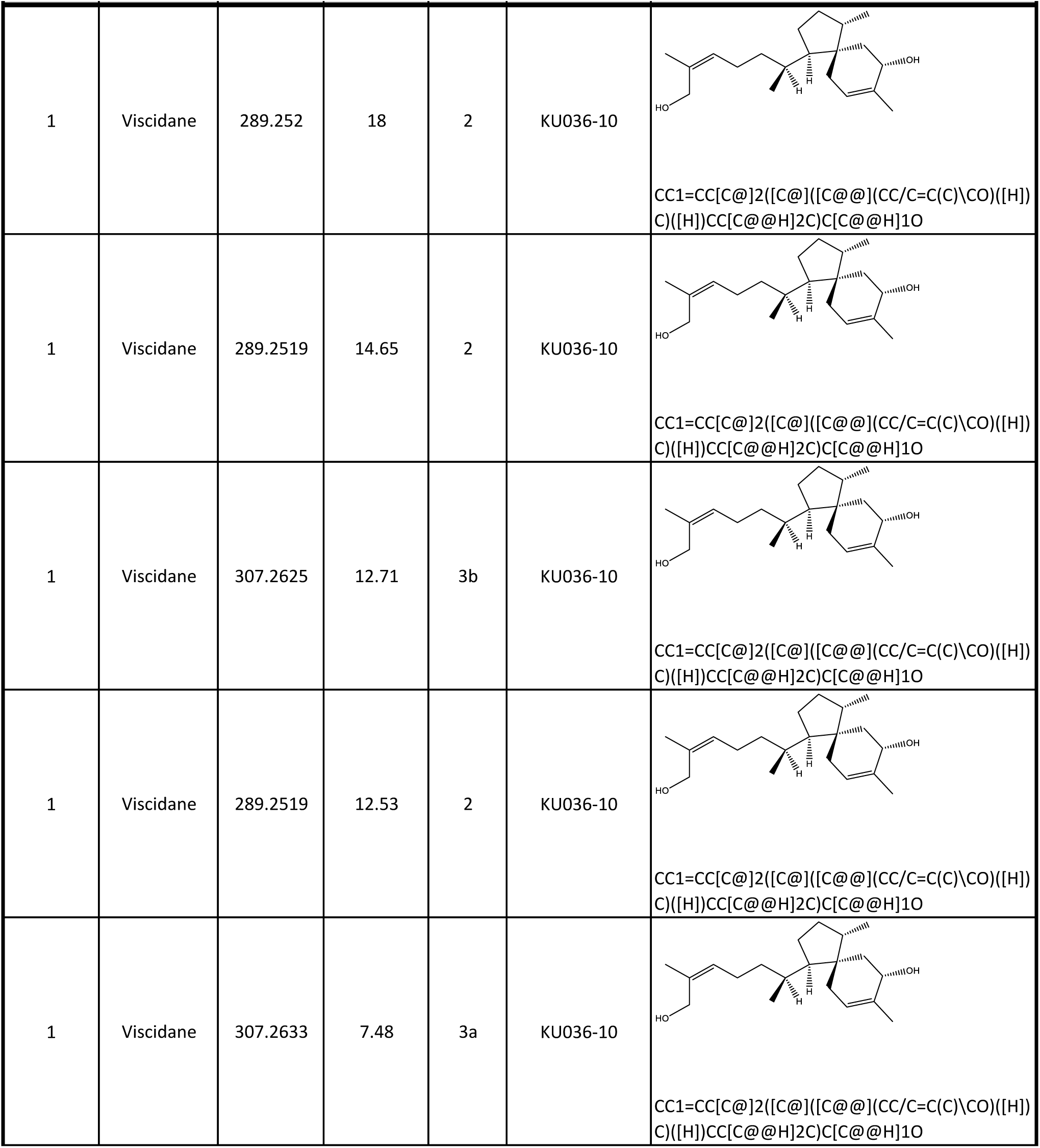
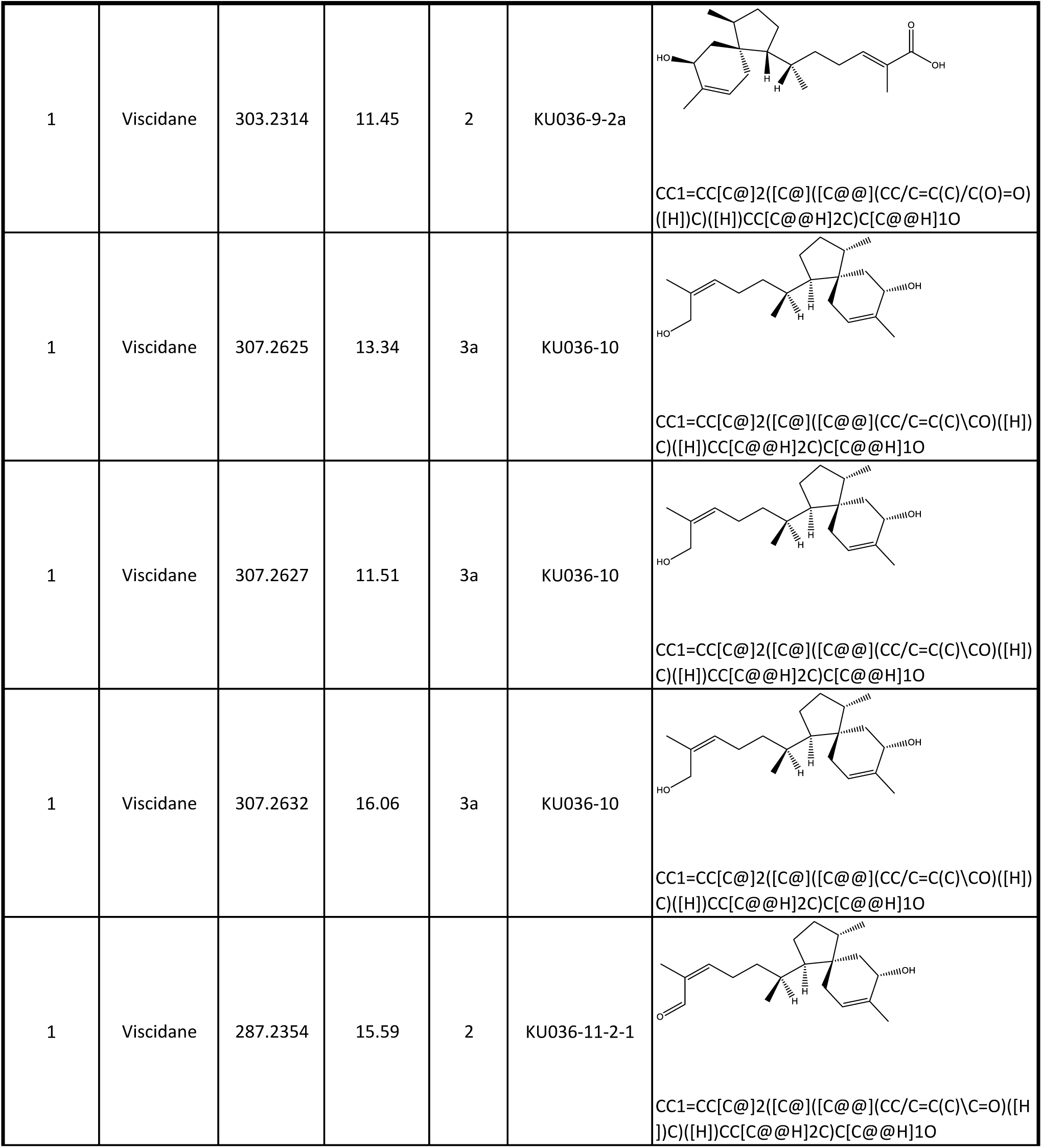
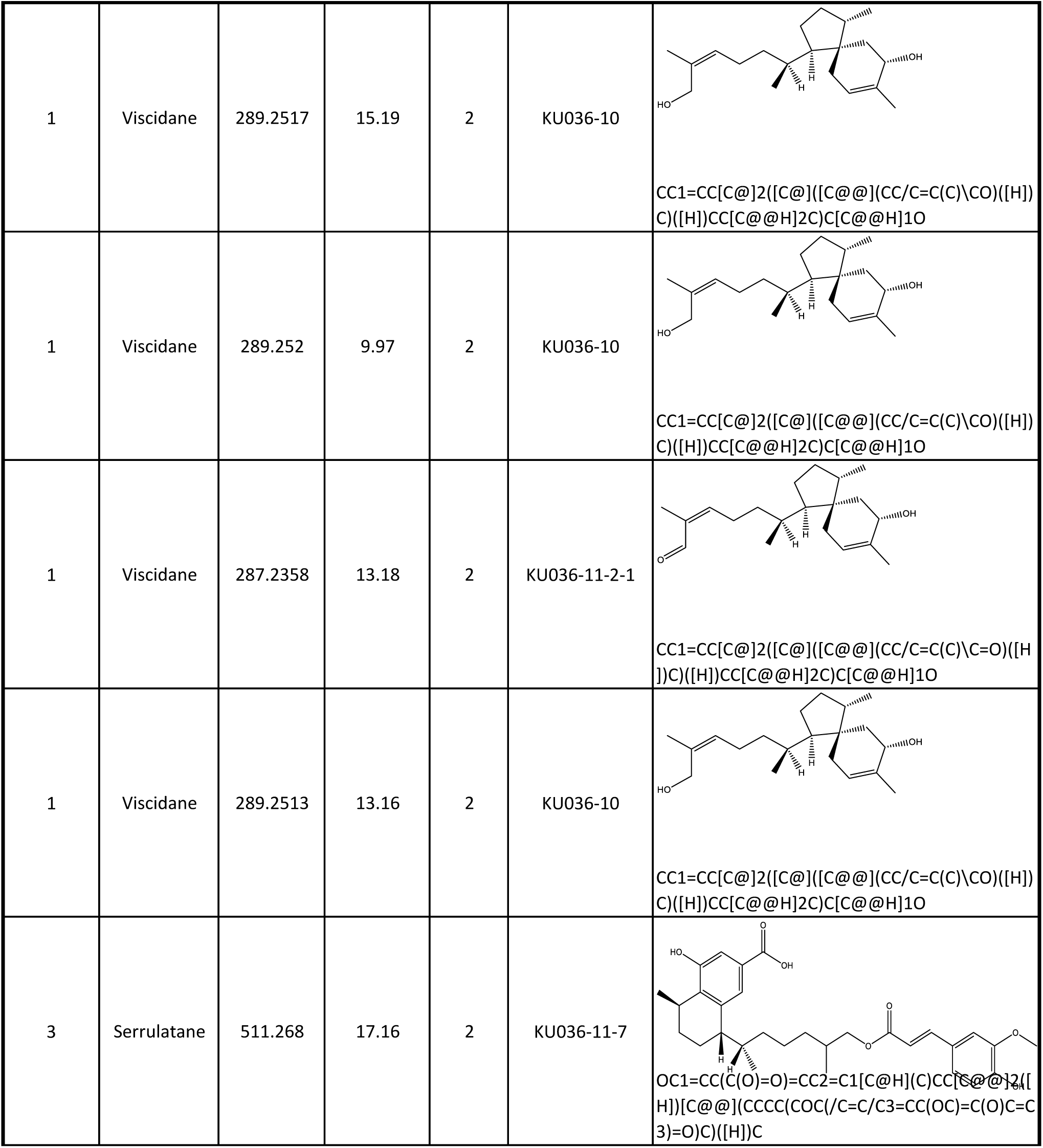
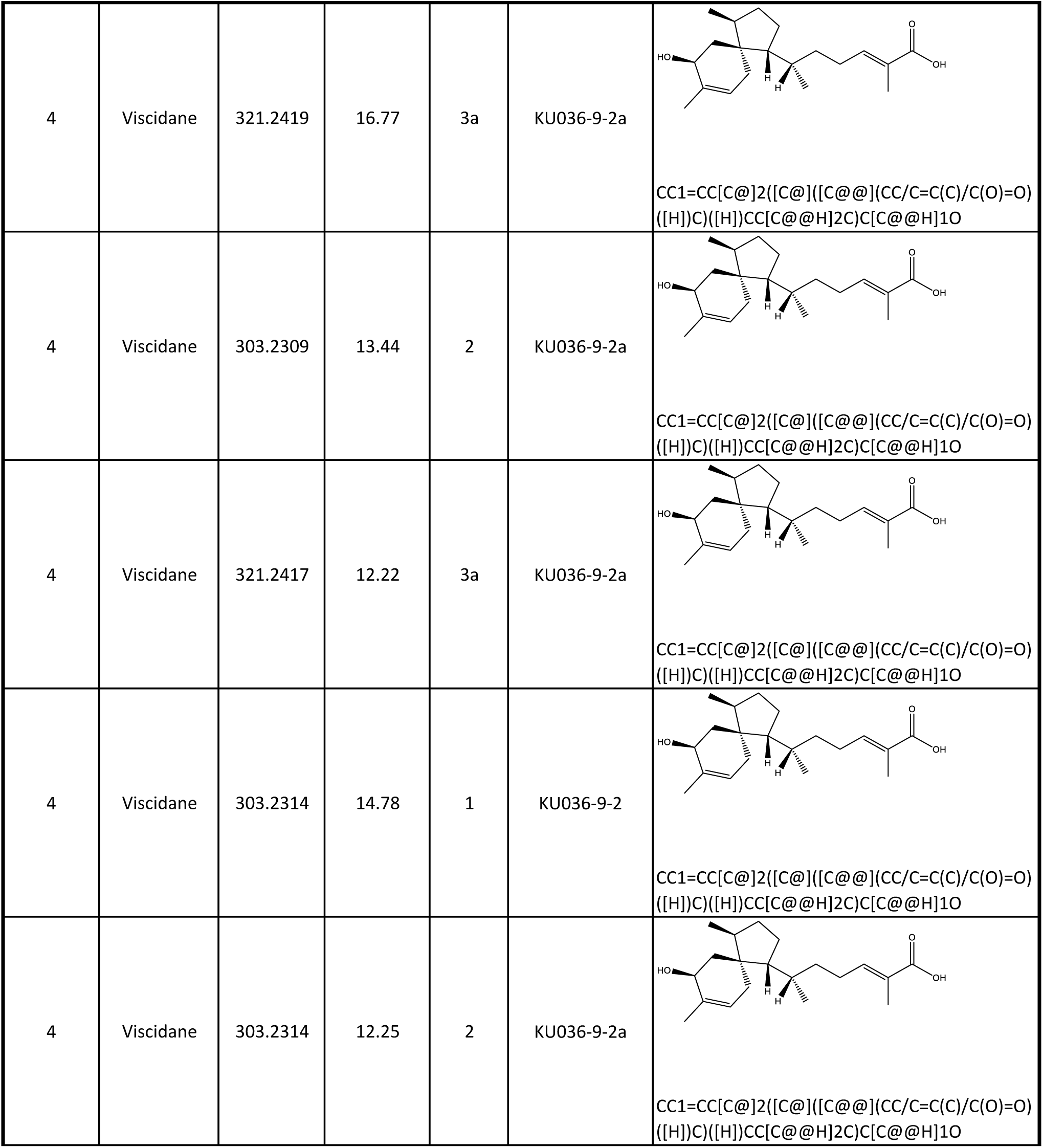
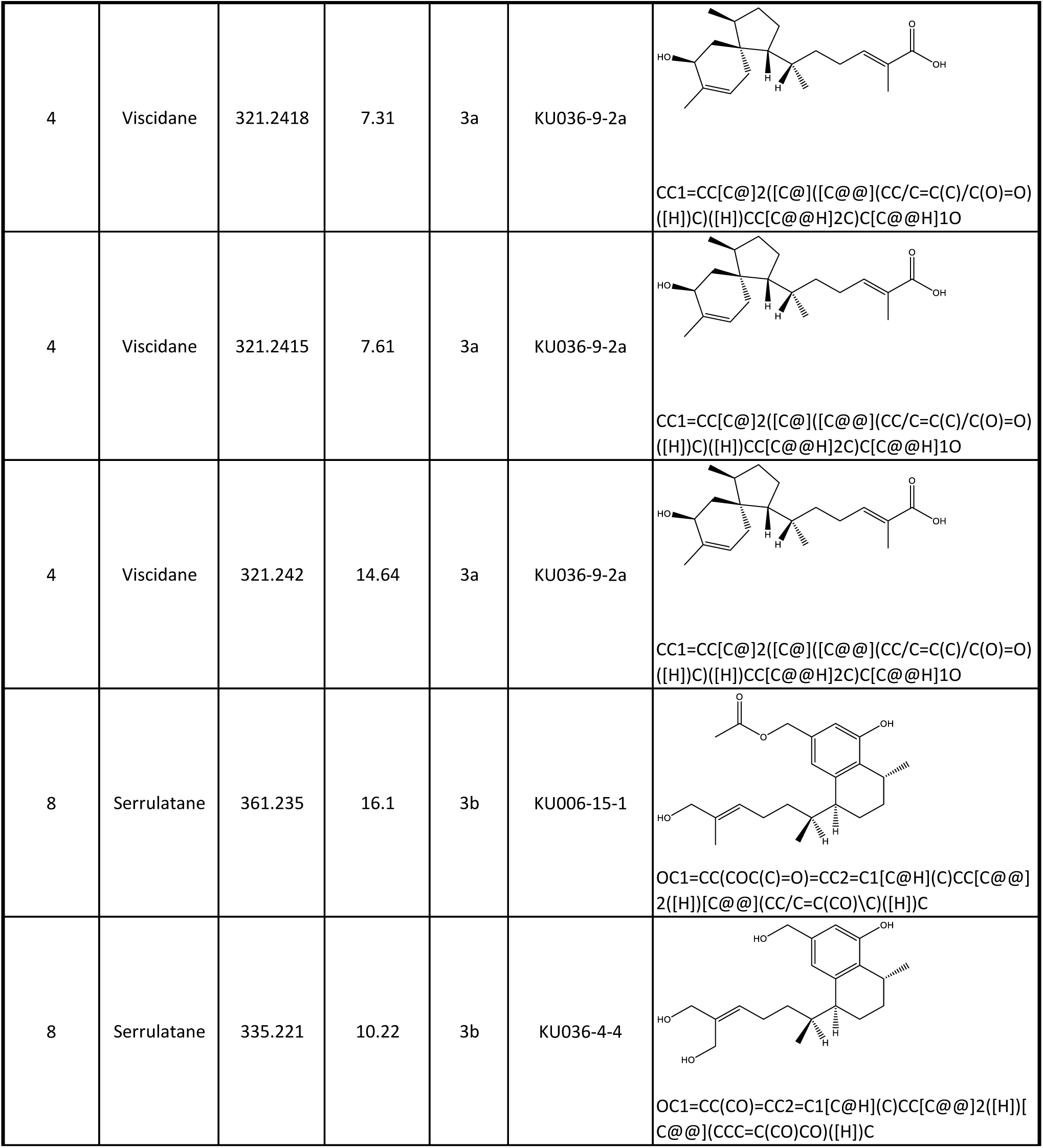
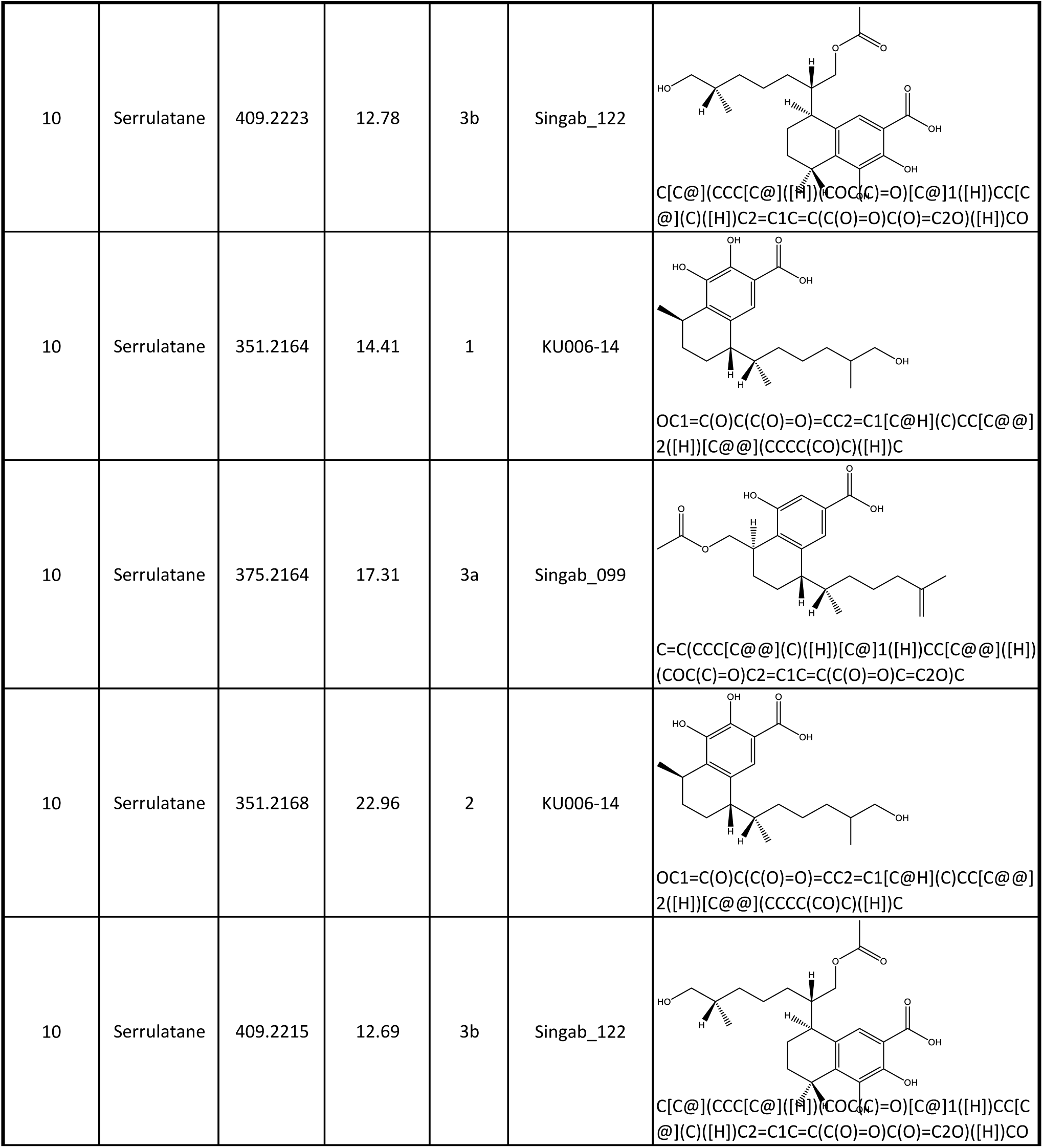
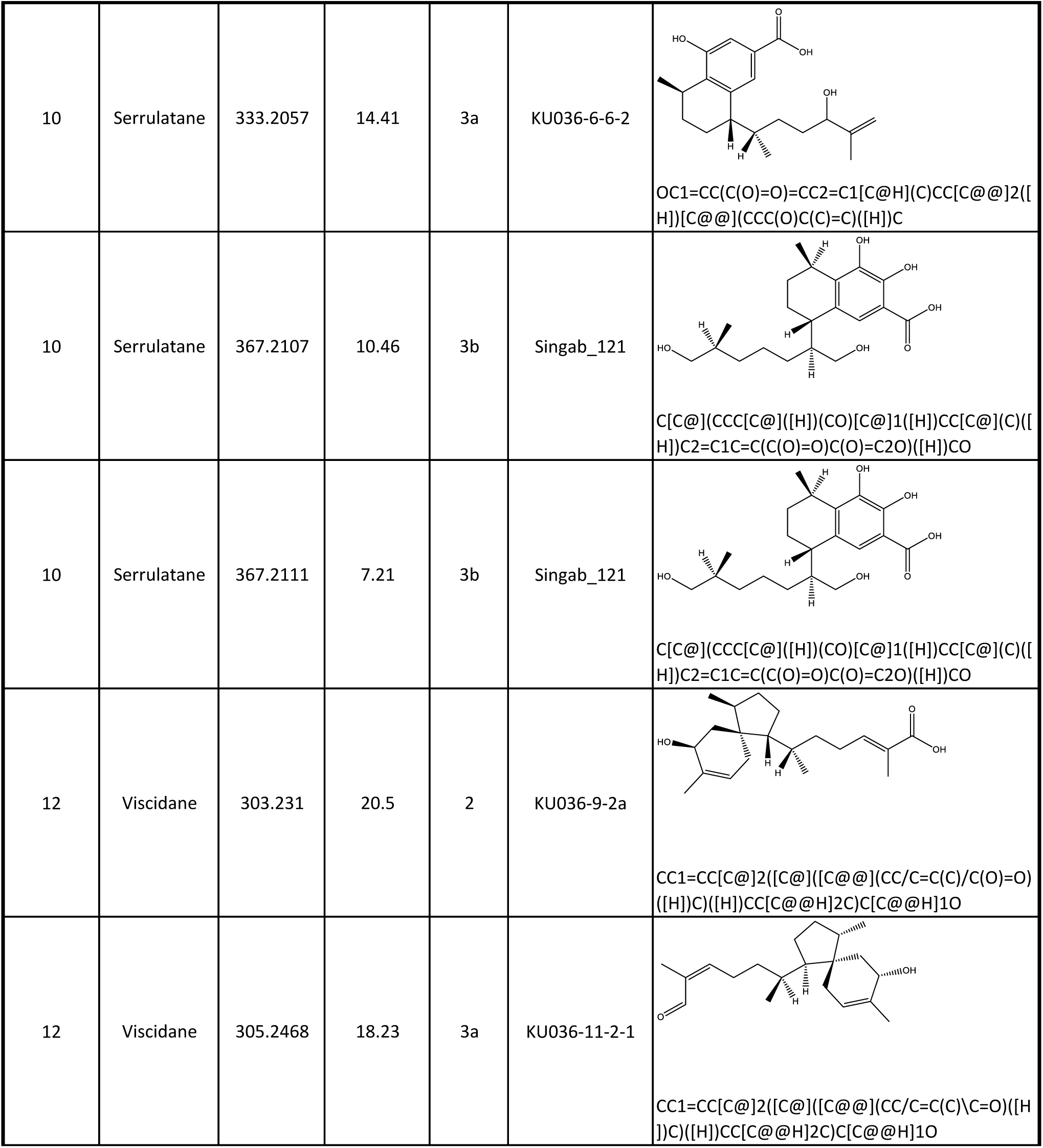
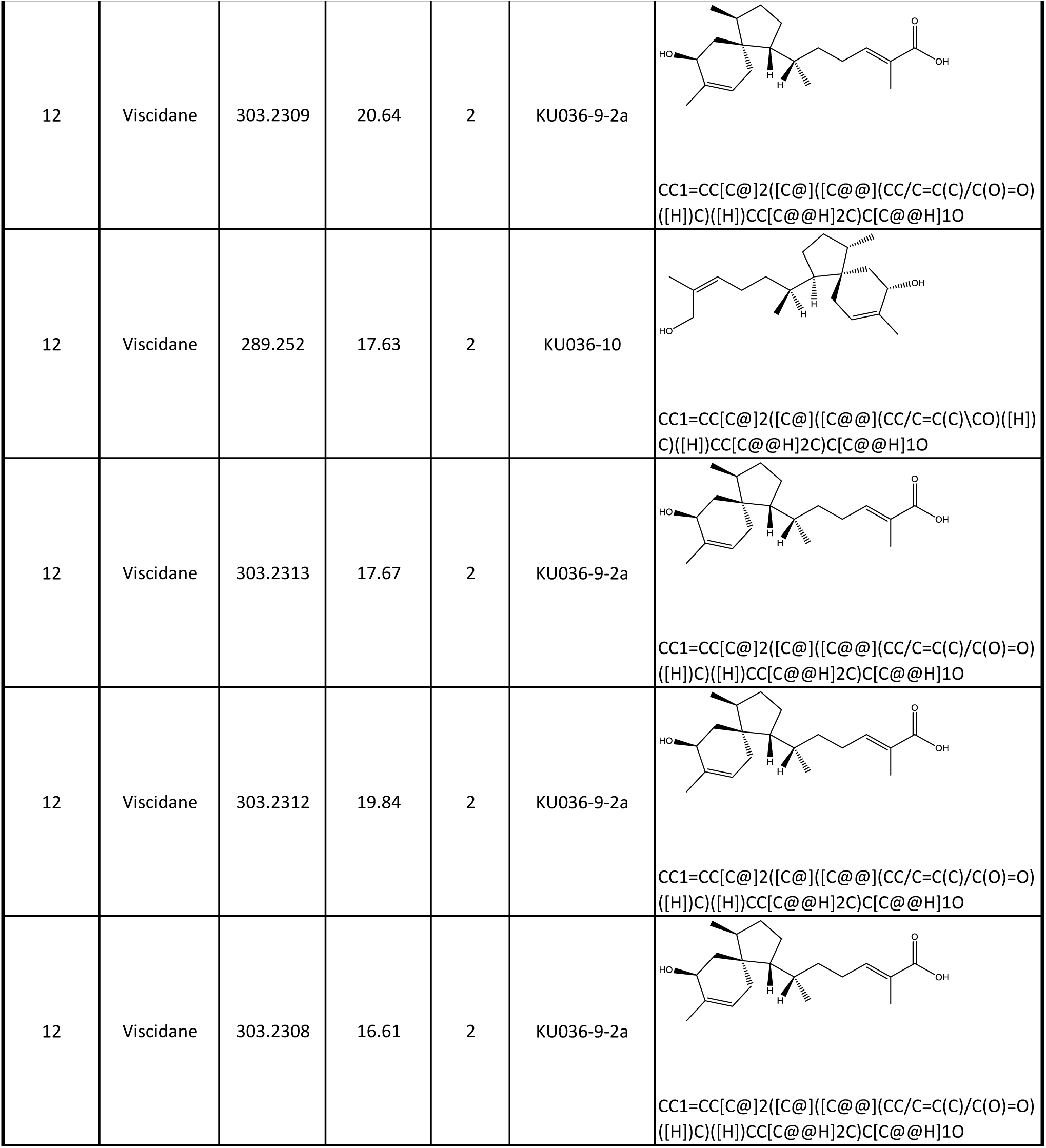
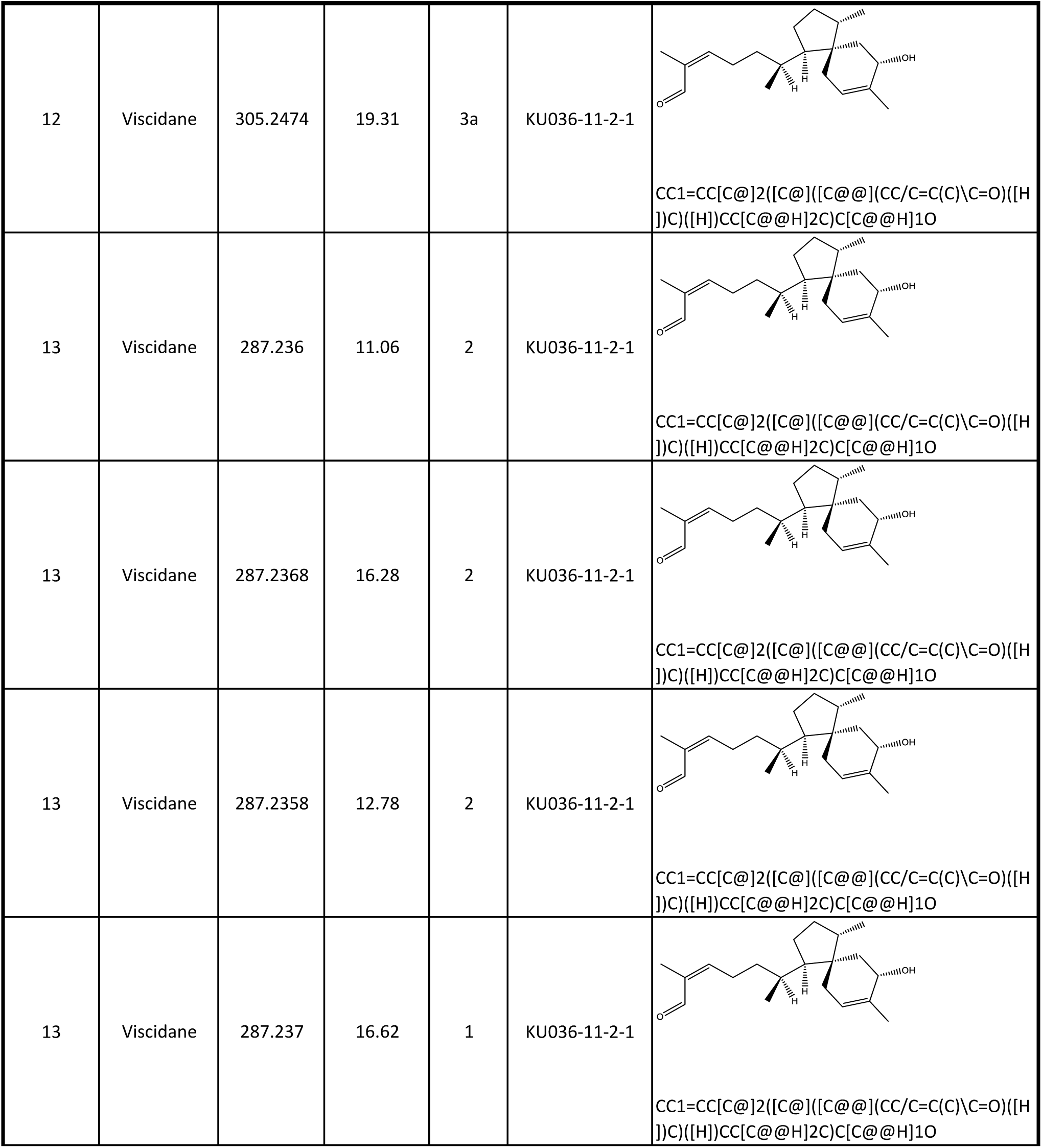
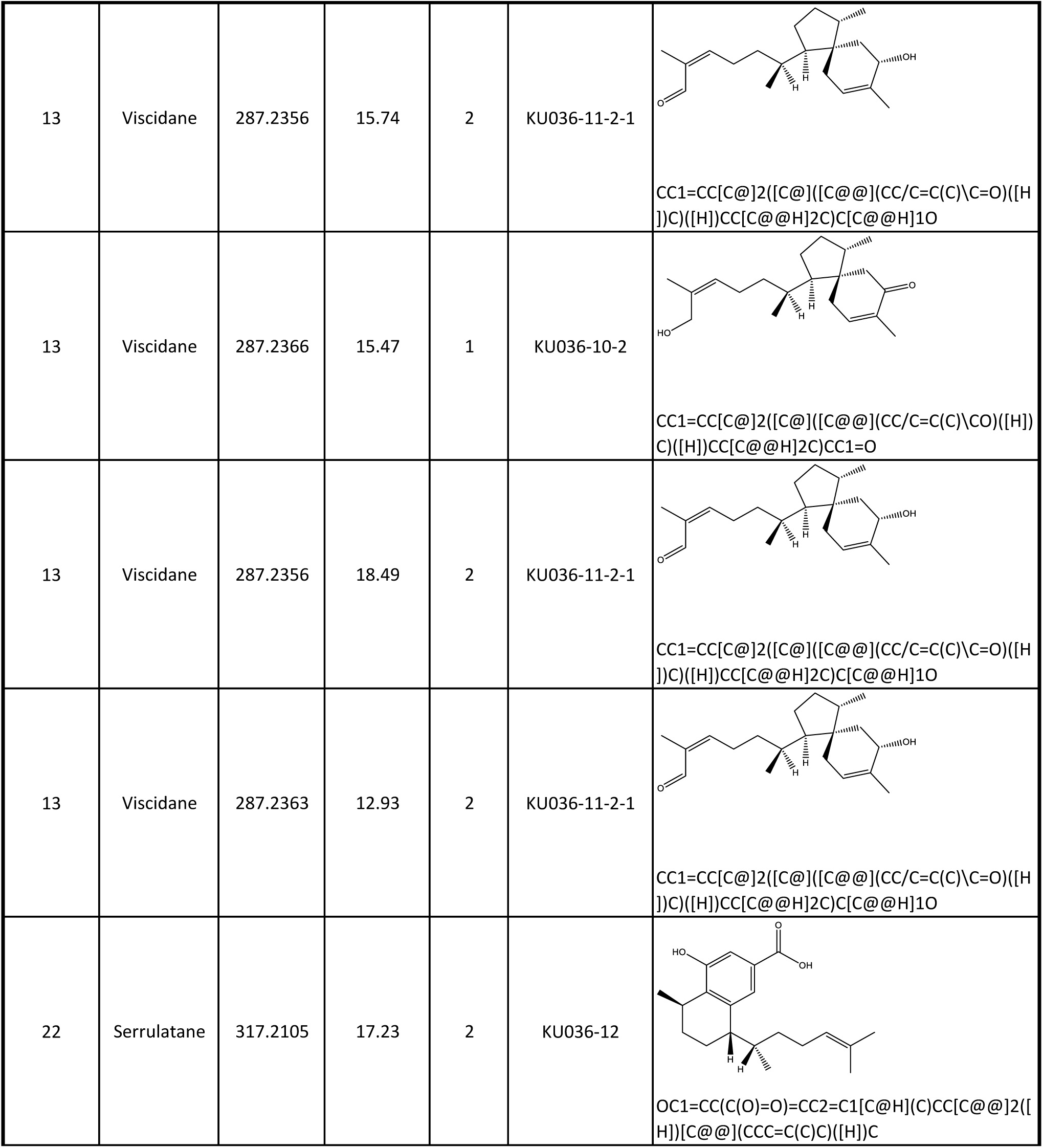
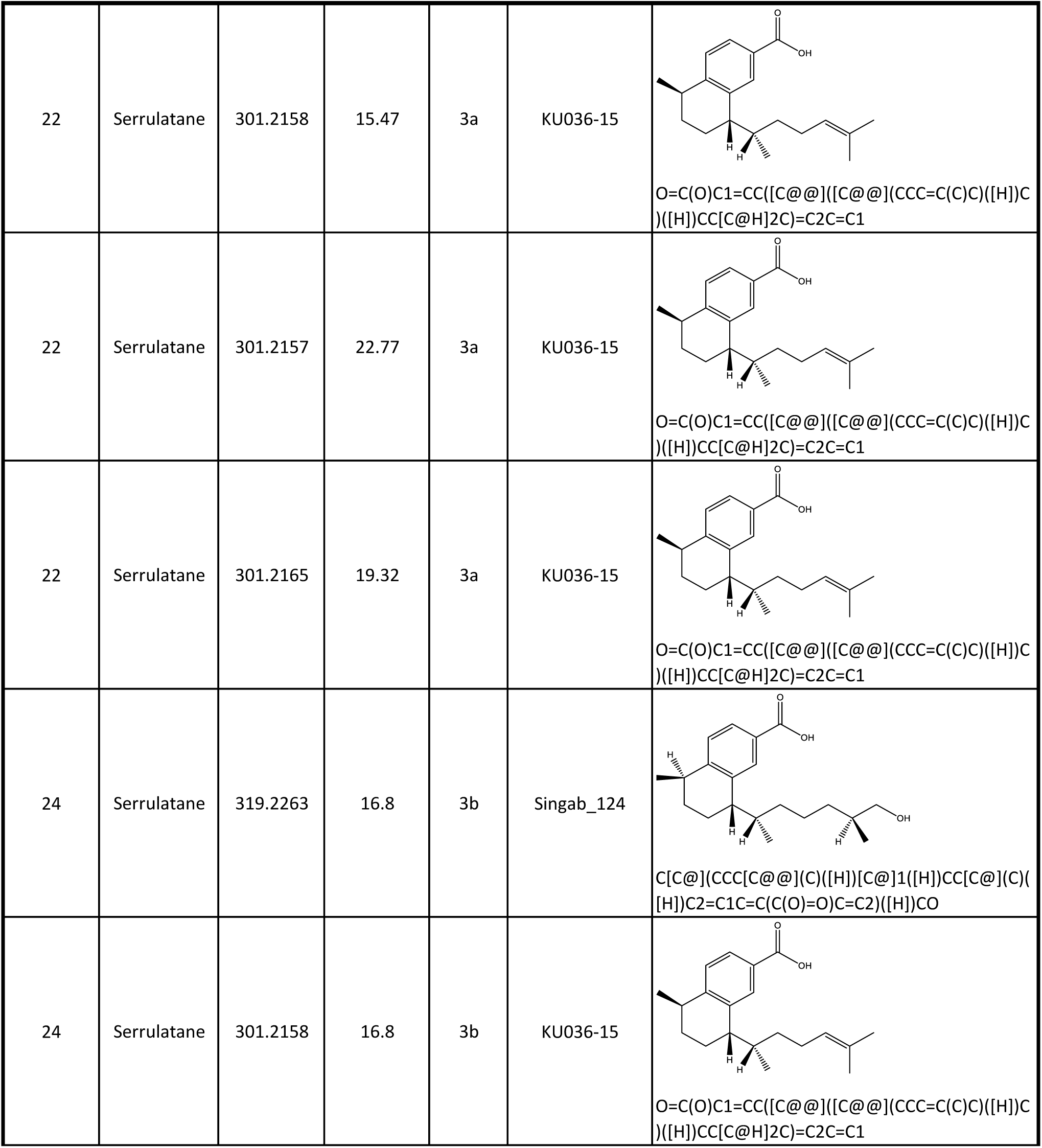
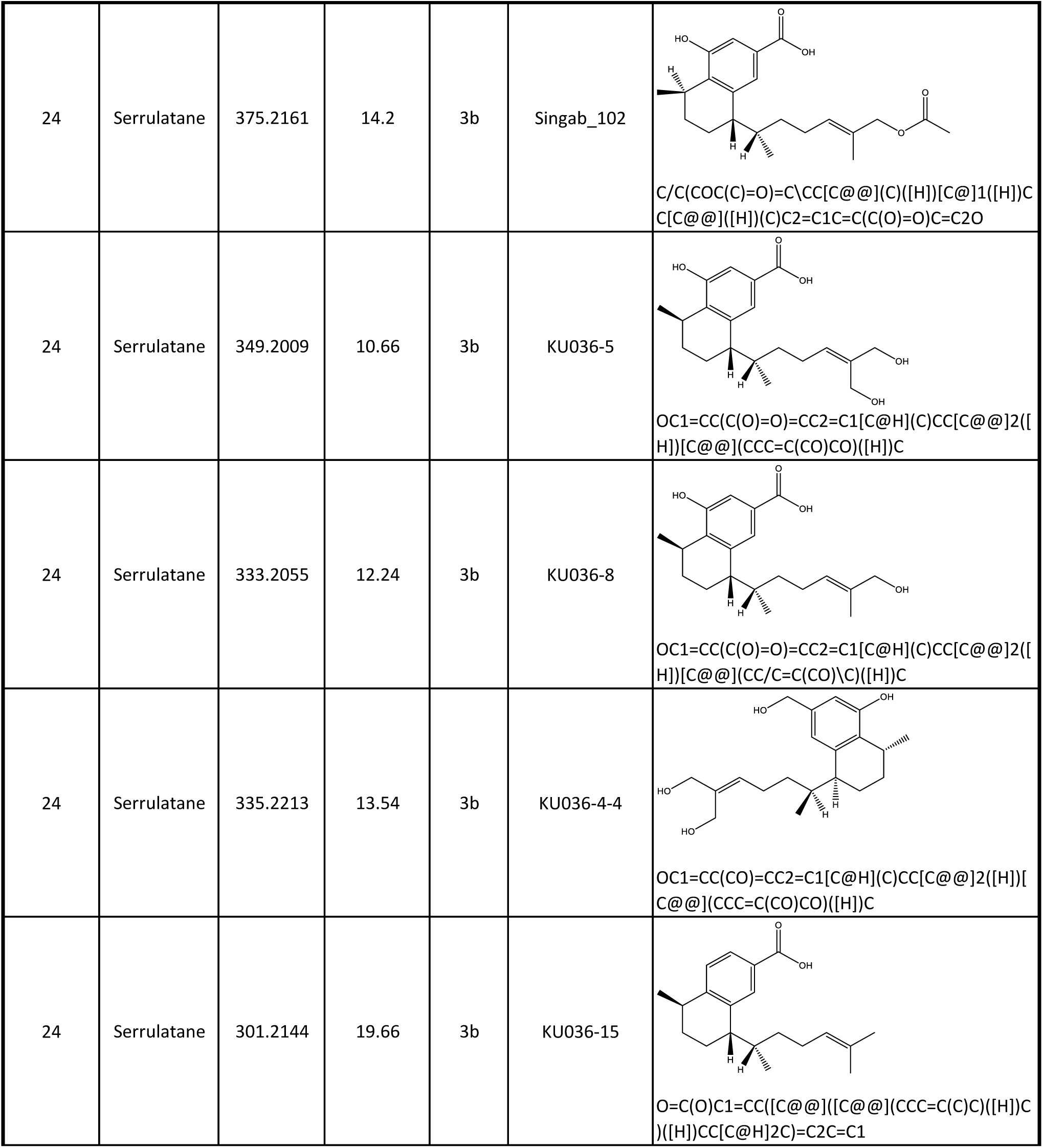
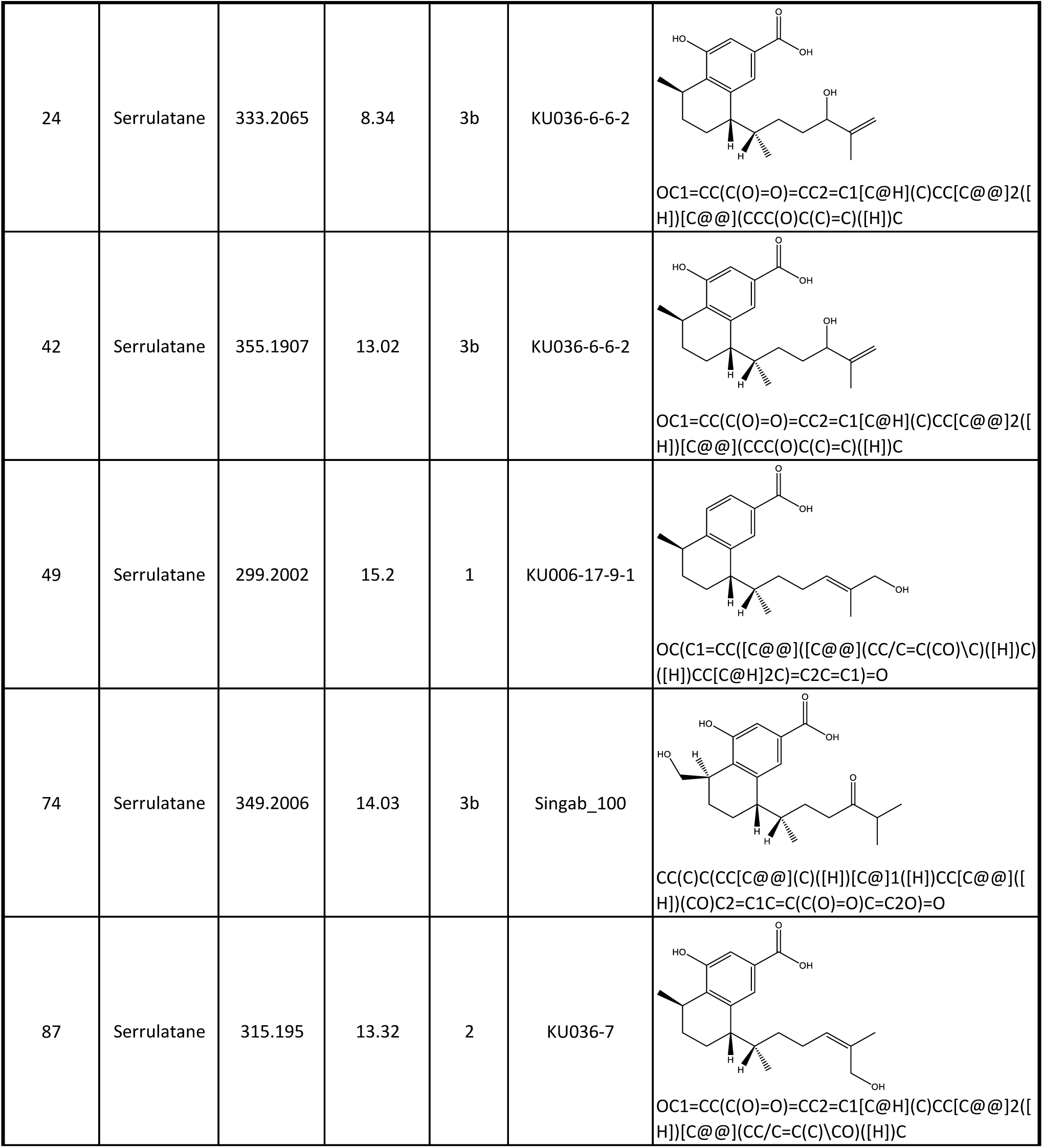
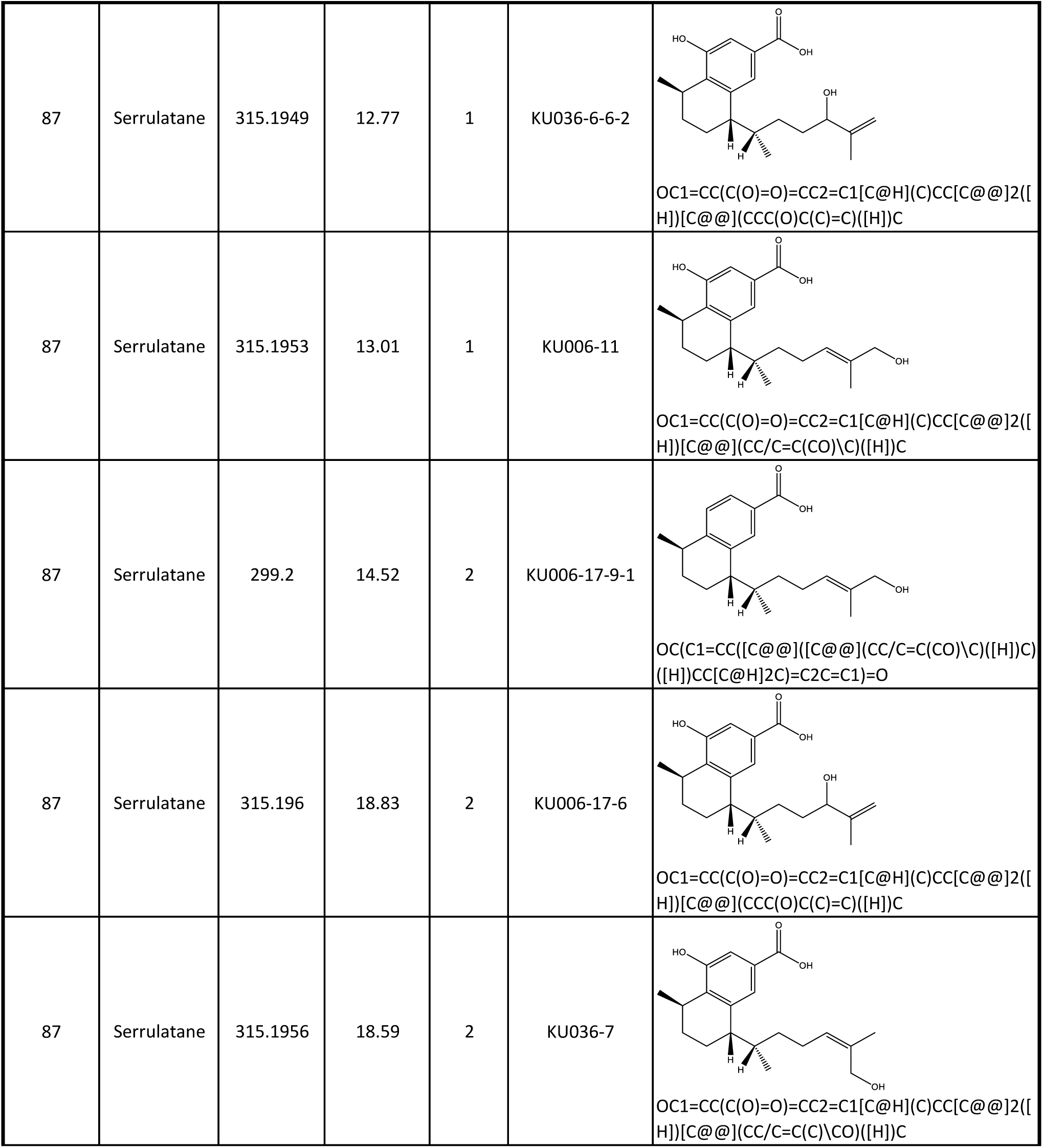
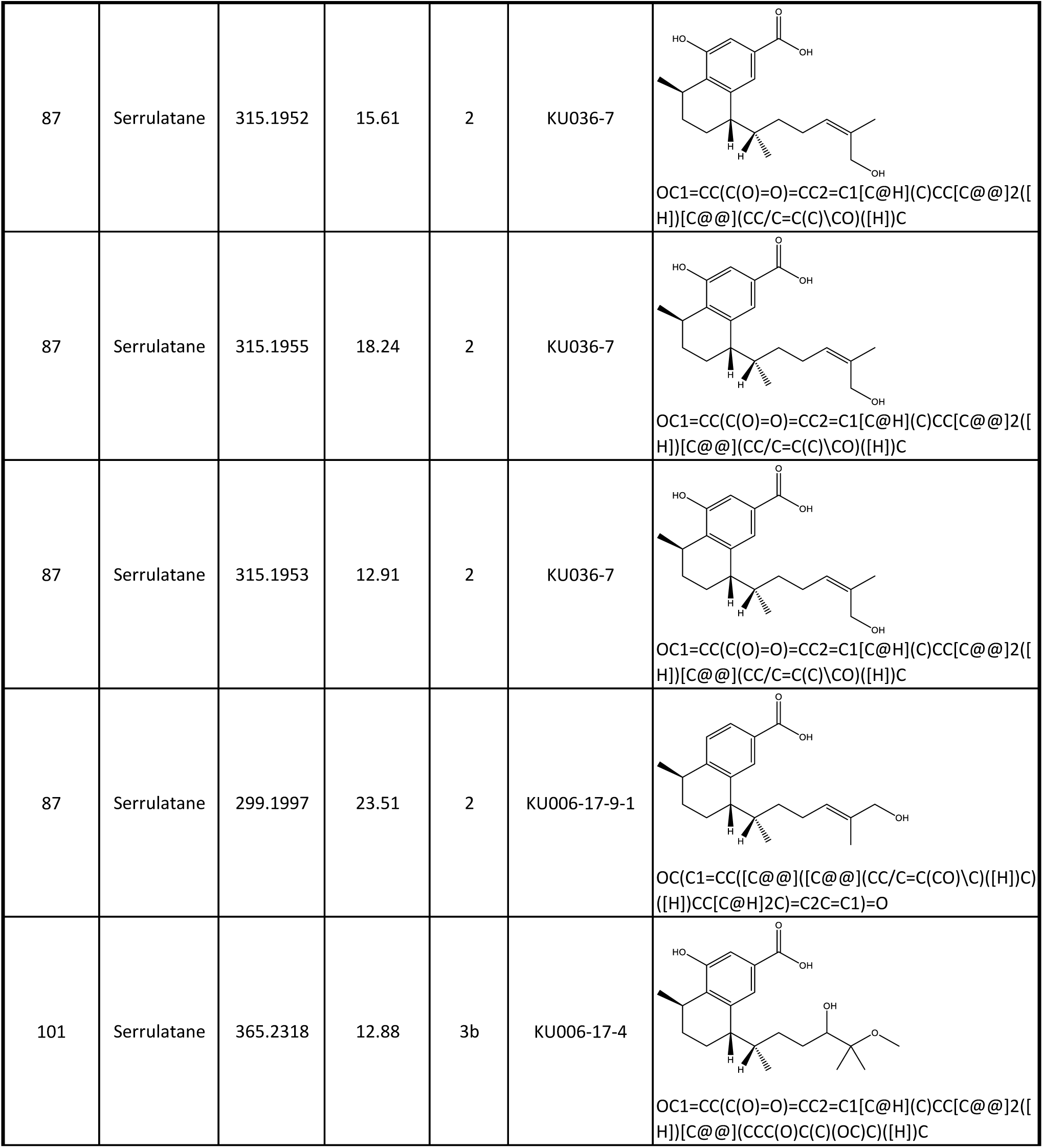
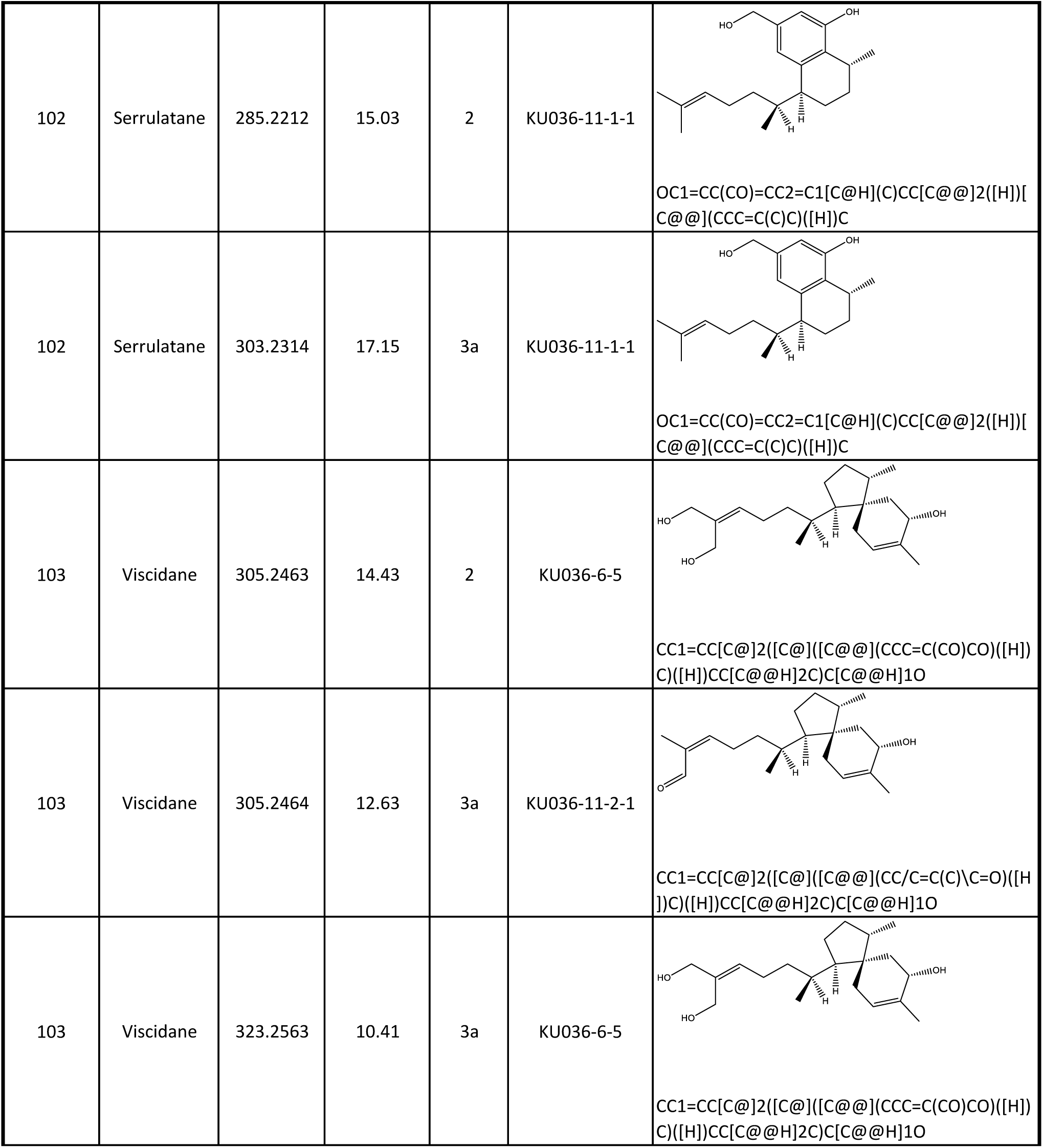
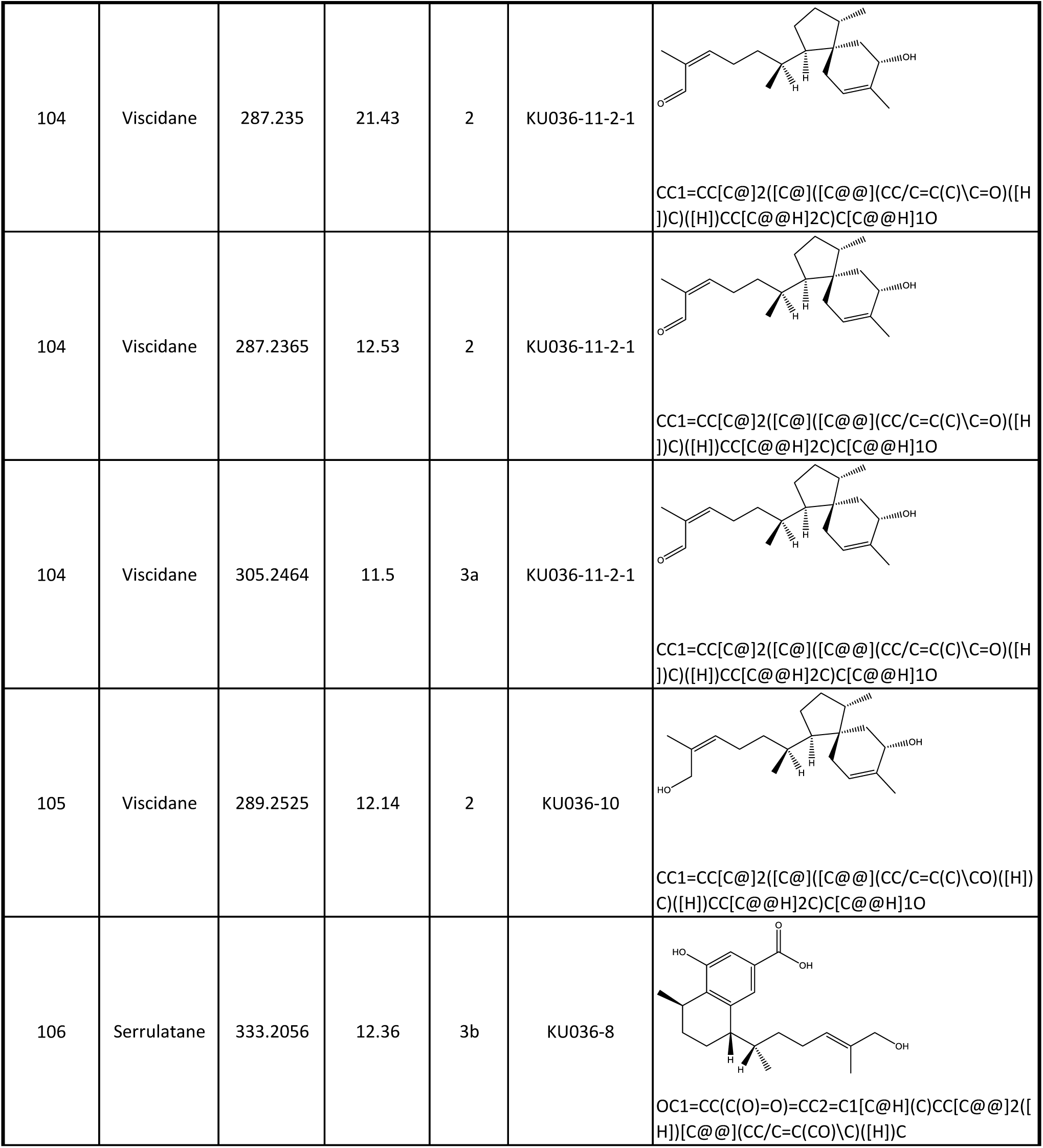
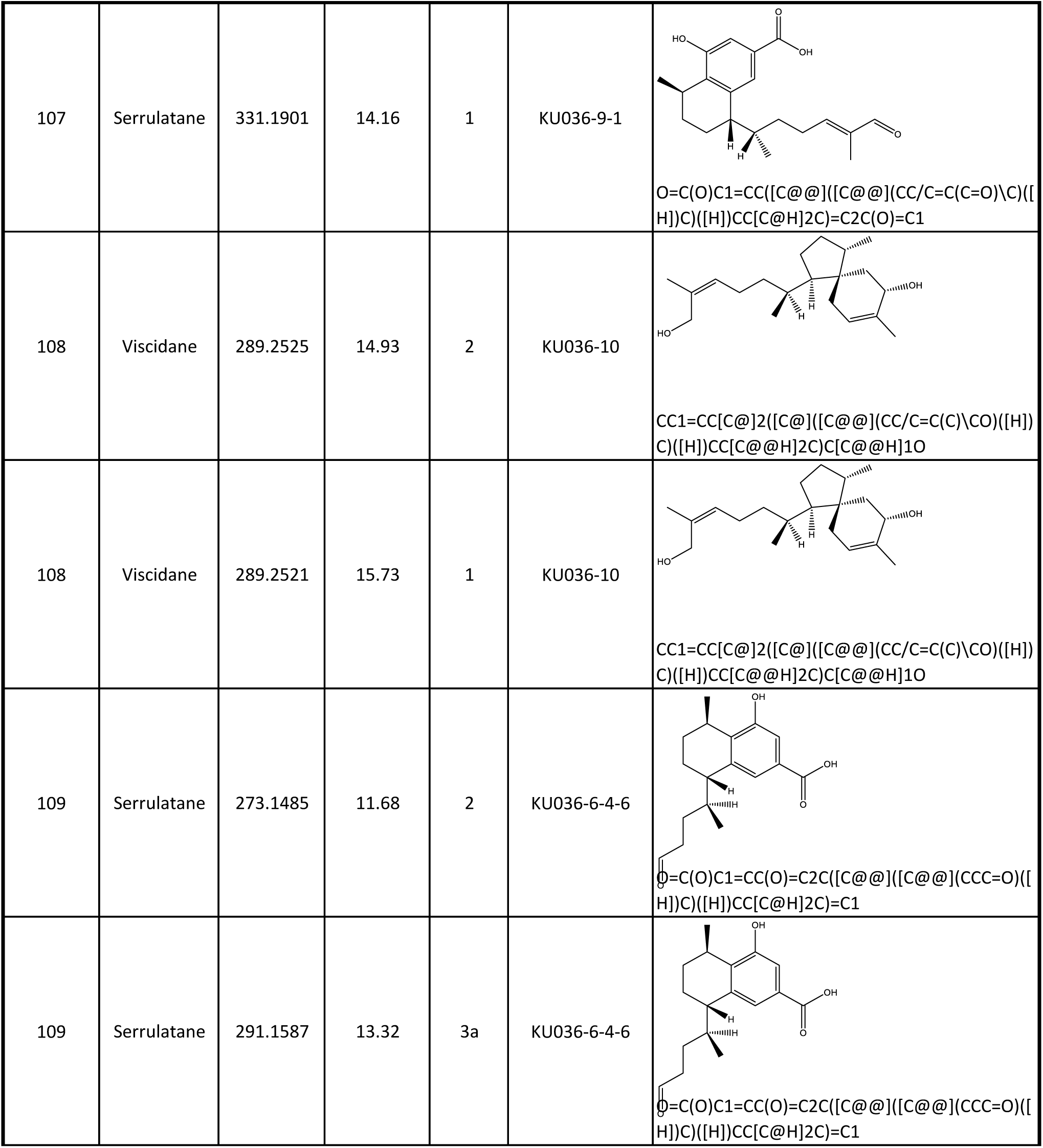
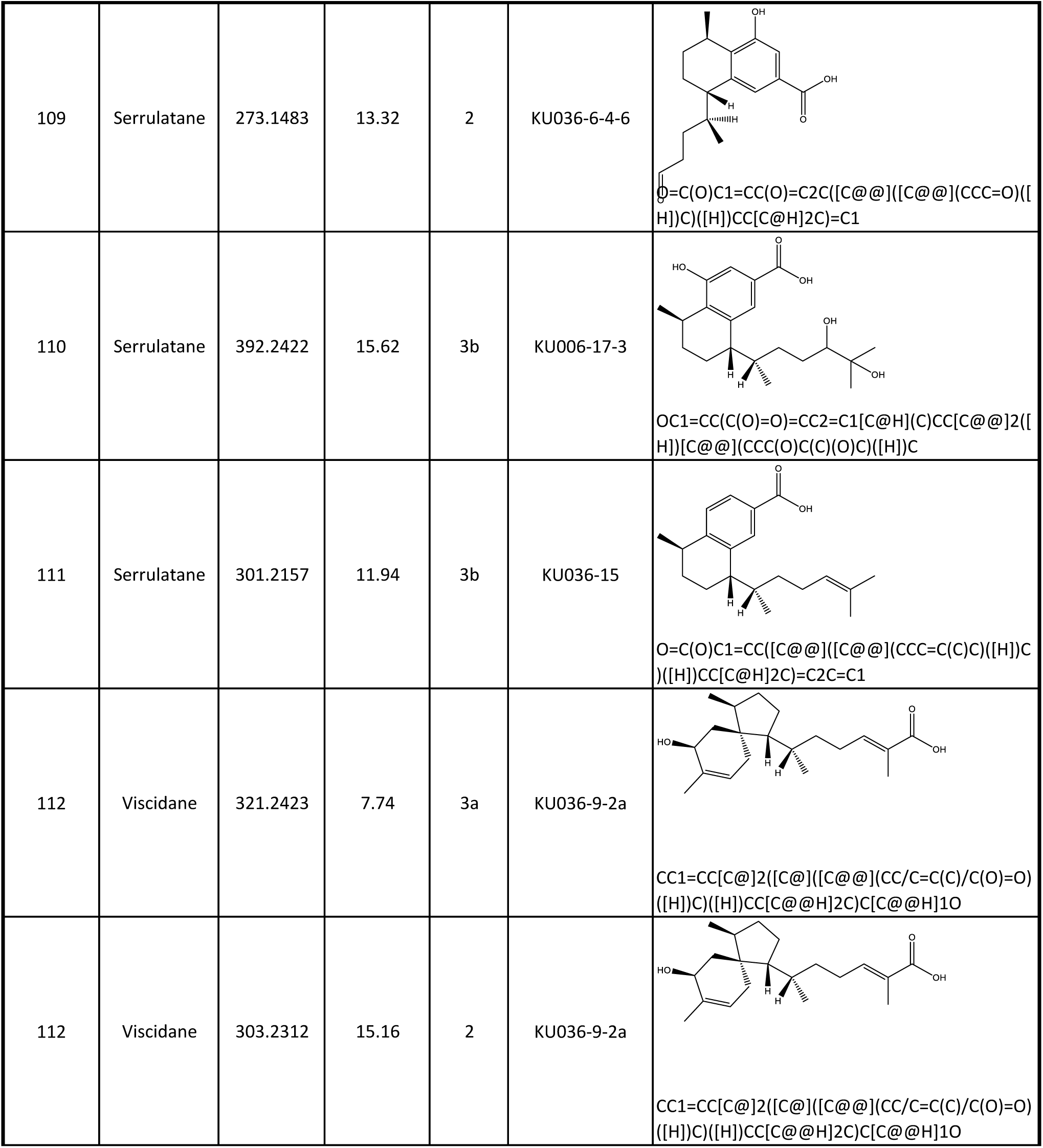
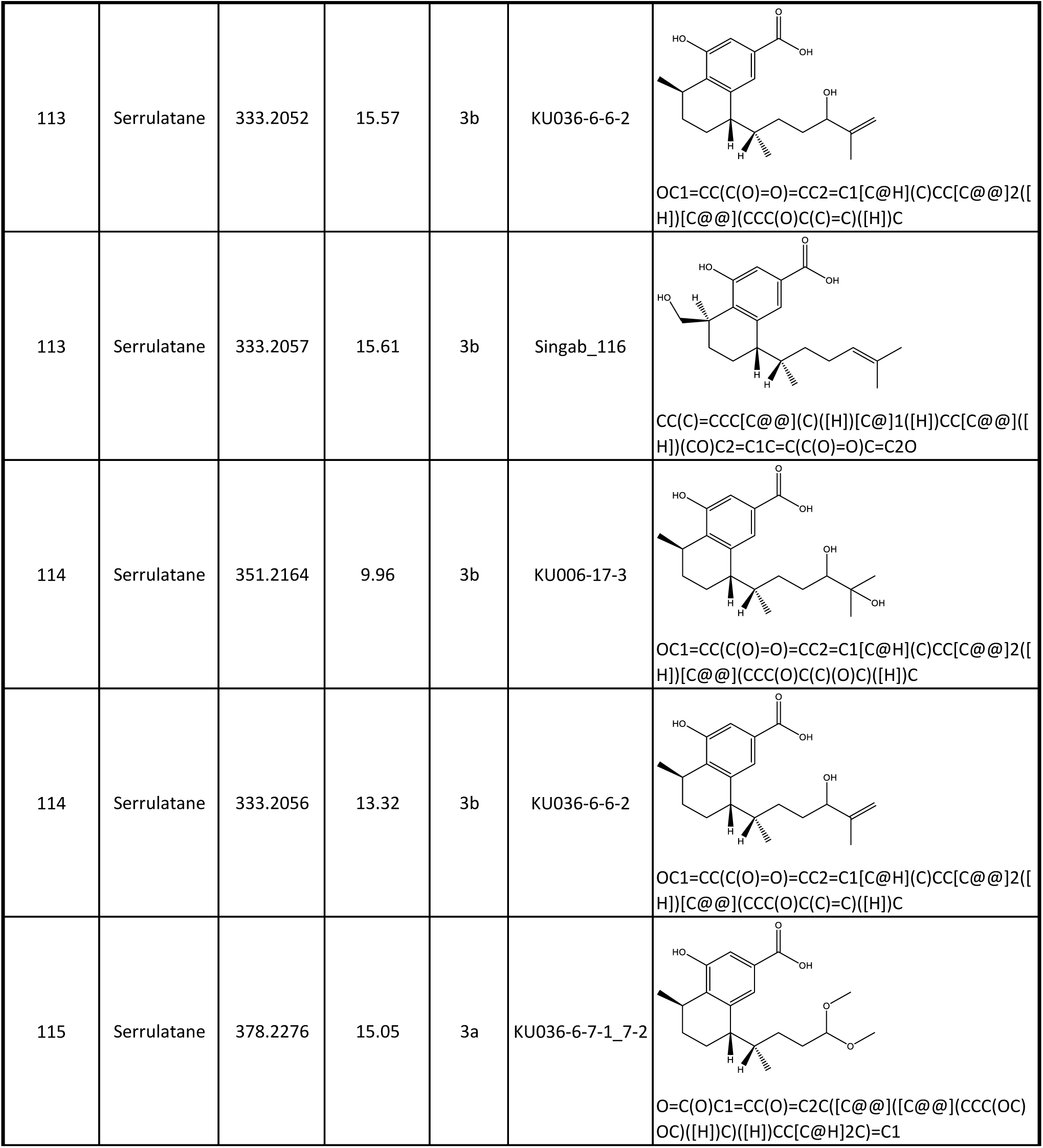
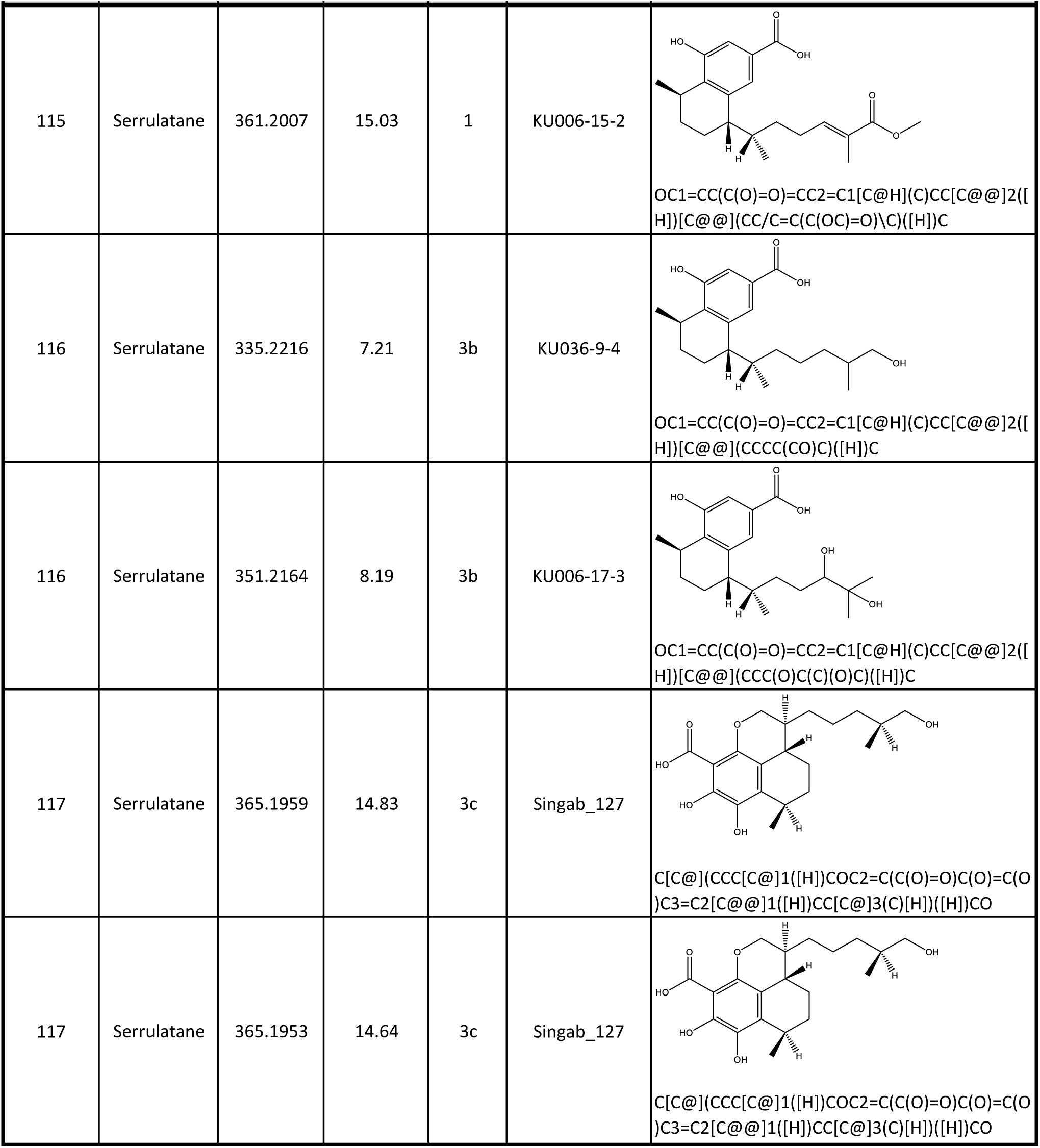
Predicted serrulatane and viscidane diterpenoid structures in chemical families of Myoporeae. Detailed spectrometric and structural information about serrulatane and viscidane diterpenoid-related identification events determined within the 30 chemical families within the Myoporeae molecular network that have been found to be associated with this specific chemistry.

### Supplementary figures

**Figure S1.**
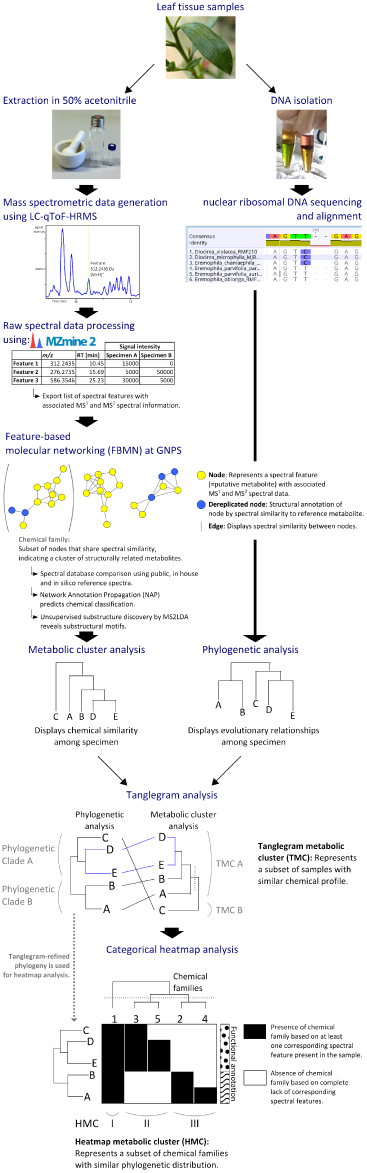
Schematic workflow of the chemo-evolutionary analysis. 291 specimen involving species and subspecies of the tribe Myoporeae were investigated regarding their chemical and phylogenetic relationships. LC-qToF-HRMS was applied on crude leaf extracts to generate mass spectrometric data including MS^1^ and MS^2^. MZmine2 was used to isolate spectral features from the raw data and align those across all samples to output a feature table that contains the relatively quantified chemical composition of the dataset. Spectral information of those isolated features is then exported and integrated into the feature-based molecular networking (FBMN) pipeline at GNPS (https://gnps.ucsd.edu). The generated molecular network is composed of nodes and edges that represent individual spectral features and shared spectral similarity, respectively. Spectral similar nodes fall into clusters of varying size that are considered as a chemical family of structurally related metabolites. The molecular network provides the foundation for an exhaustive dereplication approach involving public, in house and *in silico* libraries of reference metabolites for spectral comparison. Dereplicated nodes are structurally annotated and utilized to achieve a prediction of chemical classification for each chemical family by using the Network Annotation Propagation (NAP) method. Using the unsupervised substructure discovery approach MS2LDA, substructural motifs can be predicted within the given spectral data and annotated throughout the molecular network. To compare the chemical profiles of all 291 specimen, the presence (at least one spectral feature present of a corresponding chemical family) and absence (no feature present in the sample) information of chemical families can be used. This approach yields a binary dataset that can be compiled by a metabolic cluster analysis into a dendrogram that displays chemical similarity among the specimen. In contrast, a phylogenetic analysis shows the evolutionary relationships among those specimen and is based on nuclear ribosomal DNA sequencing of leaf tissue from the same specimens. A tanglegram analysis directly compares the analyses by rearranging branches in both dendrograms to reveal similar clustering (i.e. specimen D and E) and thus chemo-evolutionary relationships. Groups of specimen that display an evolutionary lineage are referred to as phylogenetic clades, while chemical similarity is shared within tanglegram metabolic cluster (TMC). To extend the view on the chemo-evolutionary framework, a categorical heatmap analysis is used that clusters the presence/absence of individual chemical families according to their phylogenetic distribution alongside the tanglegram-refined phylogeny. Heatmap metabolic cluster (HMC) can be derived from this approach, which represent subsets of chemical families that share a similar phylogenetic signature. Additional functional annotations can be integrated to test metadata for correlation with the chemo-evolutionary framework that is displayed by the heatmap.

**Figure S2.**
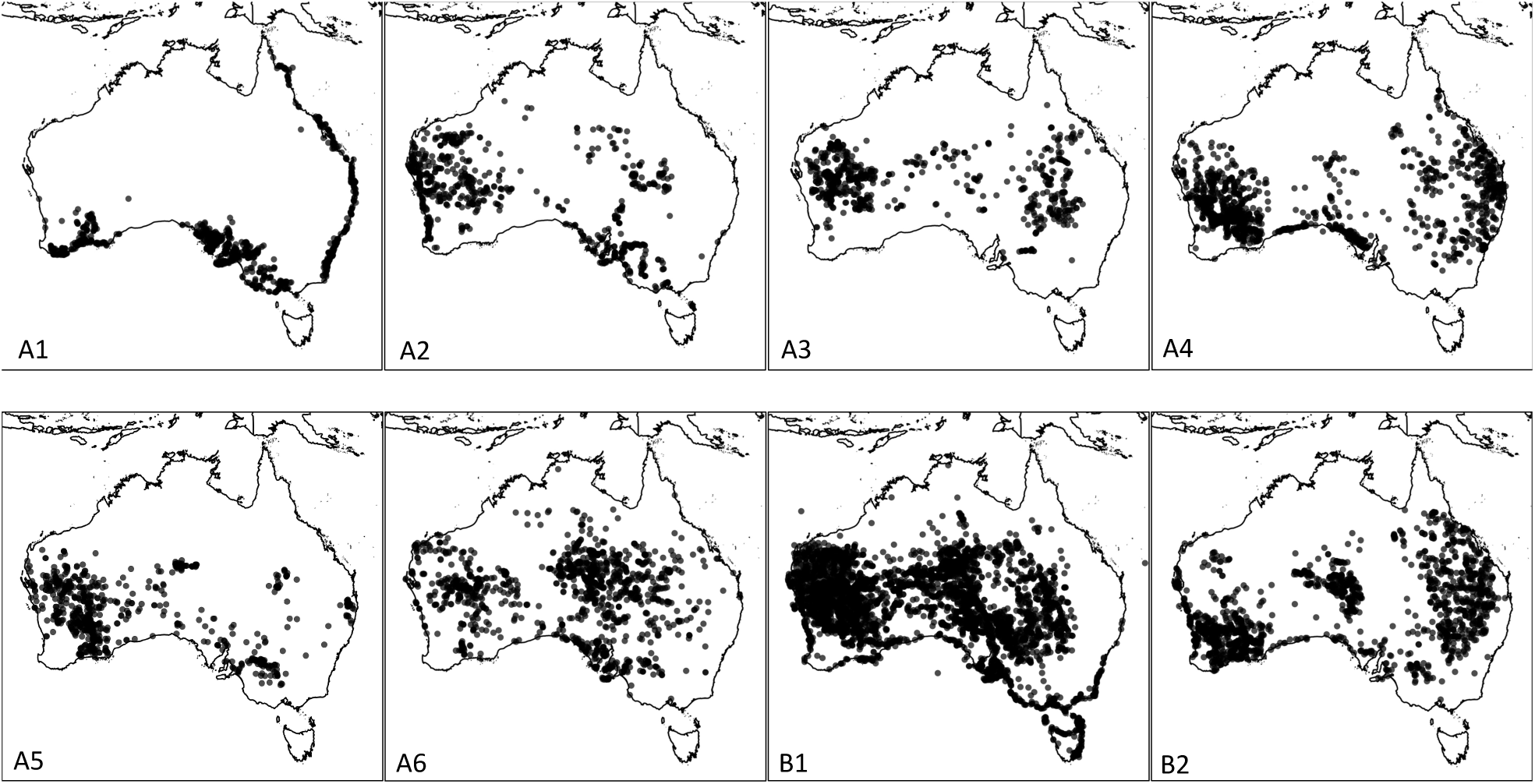
Comparison of species distribution between members of TMC A and B related chemistry. Species distributions for members of tanglegram metabolite clusters A (A1 – A6) and B (B1 – B2). Widespread species (those with broad distributions across multiple biomes) have been excluded in clusters A1 (*Eremophila bignoniiflora*, *E. alternifolia*, *E. deserti*), A2 (*Myoporum montanum*, *E. latrobei subsp. glabra*, *E. deserti*), A5 (*M. acuminatum*, *E. longifolia*) and B6 (*E. serrulata*). Data generated from the Australasian Virtual Herbarium (https://avh.chah.org.au/) as at 29/06/20.

**Figure S3.**
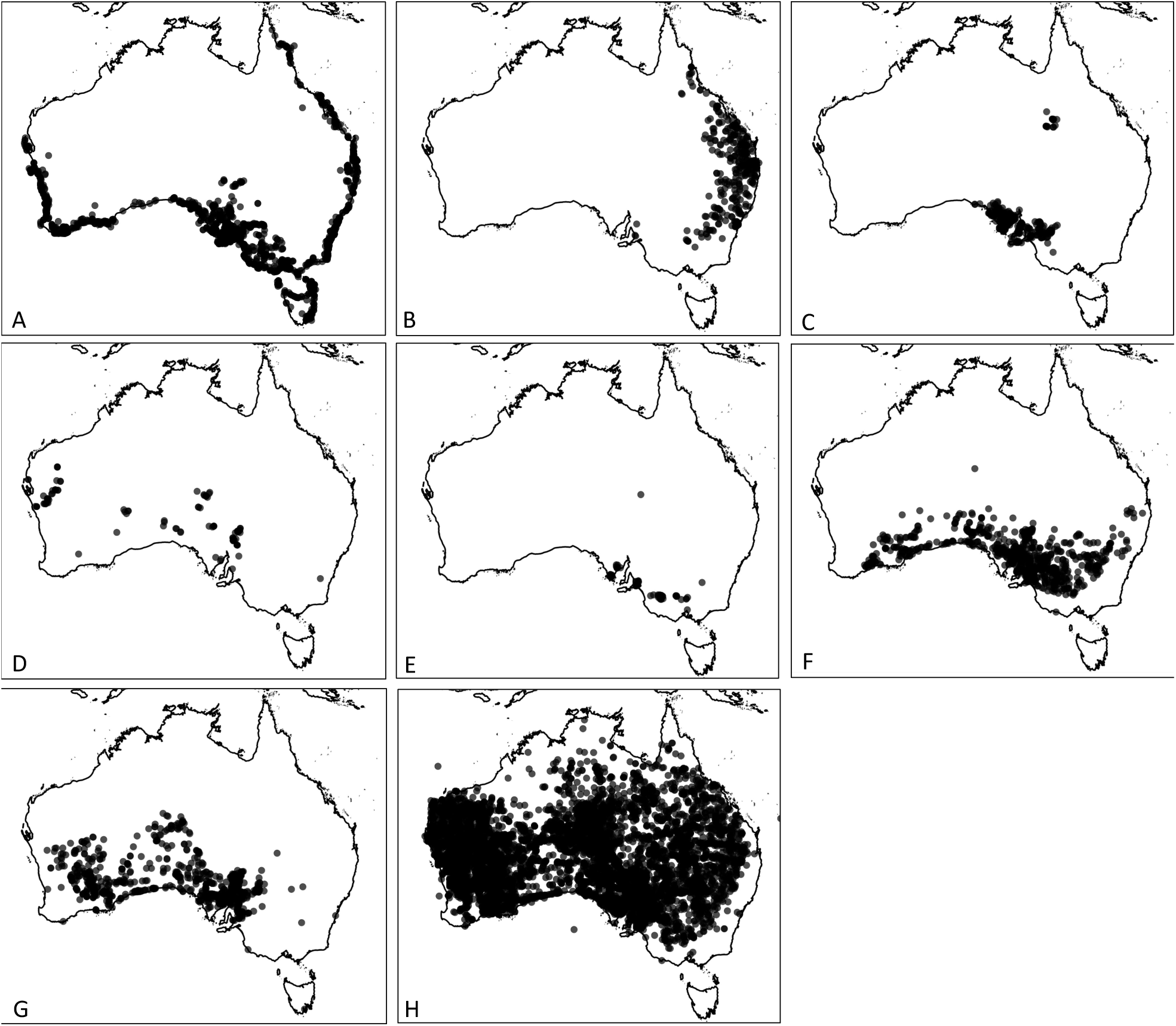
Species distributions for members of Myoporeae phylogenetic clades A–H. In this representation, widespread species (those with broad distributions across multiple biomes) have been excluded in clade A (*M. acuminatum*, *M. montanum*) and clade D (*E. bignoniiflora*, *E. deserti*, *E. polyclada*). Data generated from the Australasian Virtual Herbarium (https://avh.chah.org.au/) as at 06/04/20.

**Figure S4.**
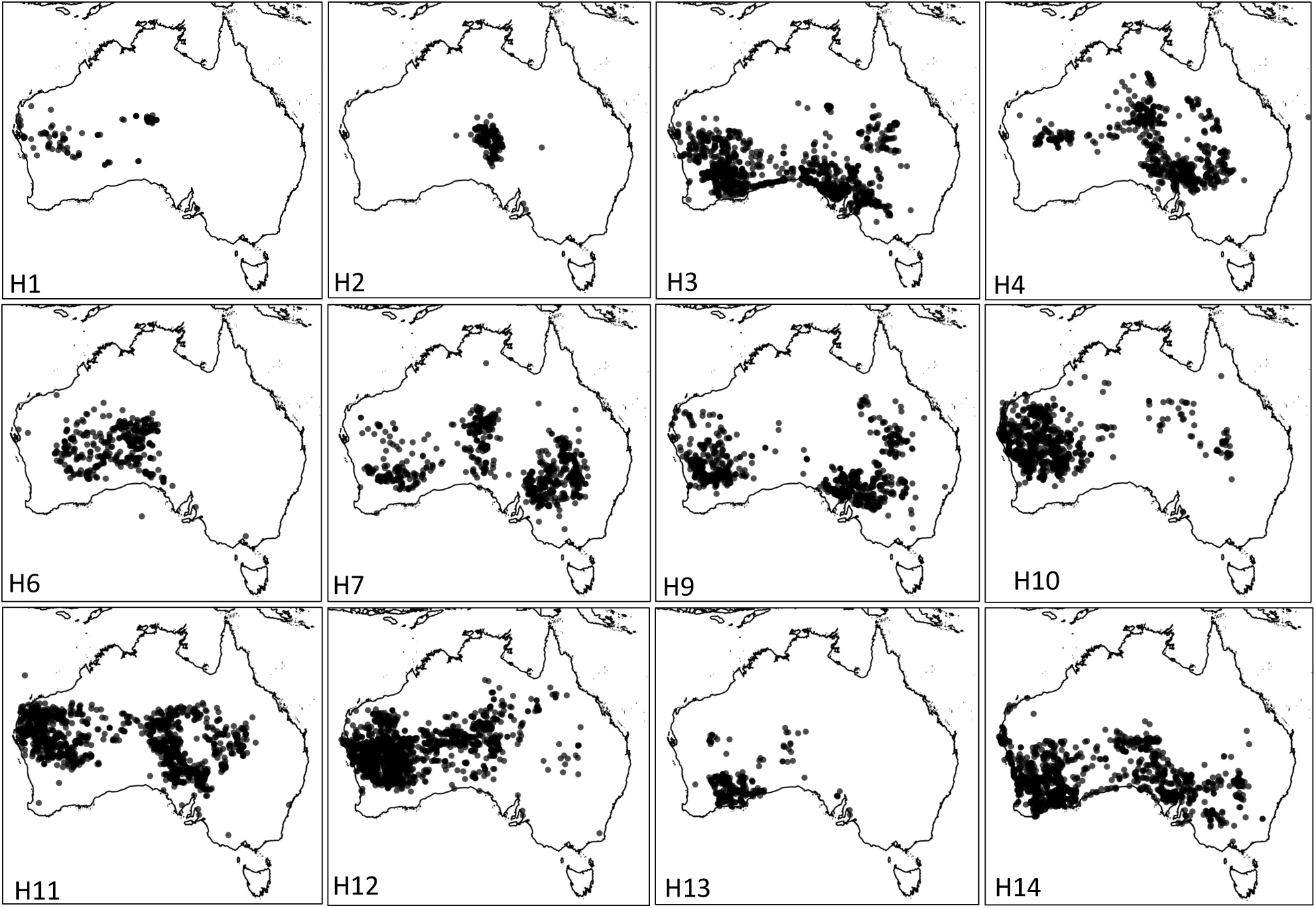
Species distributions for members of Myoporeae phylogenetic clade H, subclades H1–H14. In this representation, widespread species (those with broad distributions across multiple biomes) have been excluded in clade H2 (*E. mitchellii*), clade H5 (*E. longifolia*), clades H8 (*E. maculata*) and clade H10 (*E. latrobei*). Data generated from the Australasian Virtual Herbarium (https://avh.chah.org.au/) as at 06/04/20.

**Figure S5.**
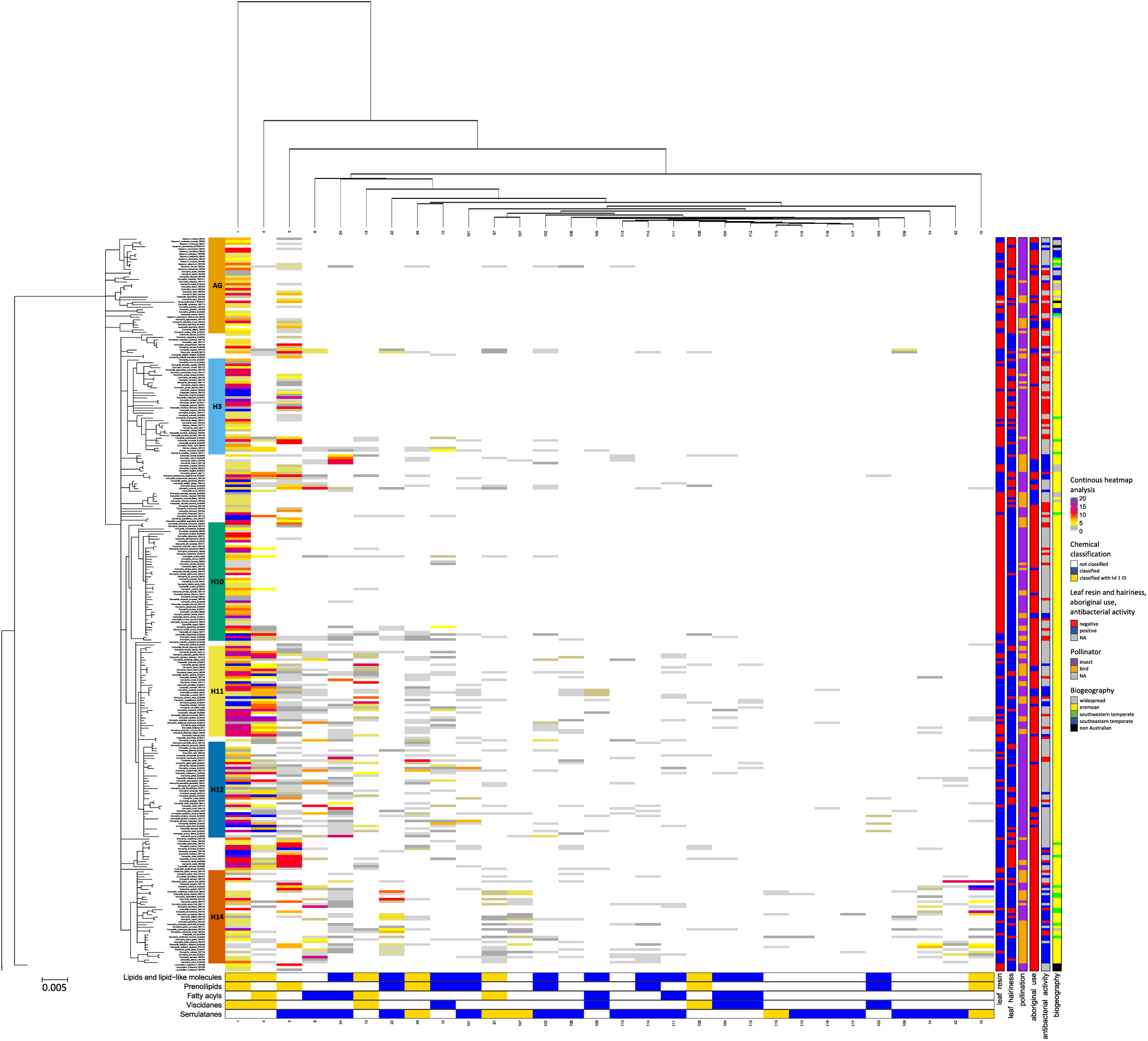
Continuous heatmap analysis of chemo-evolutionary patterns involving serrulatane and viscidane diterpenoids in Myoporeae. A heatmap analysis was used to put chemical information into an evolutionary context. For that, information about the 30 chemical families derived from the molecular network of Myoporeae, which have been found to be associated with serrulatane and viscidane diterpene chemistry was compiled. For each specimen in this study, the total amount of metabolites of a corresponding chemical family is displayed within the heatmap. *LEFT*: The nuclear ribosomal DNA phylogeny on the left side presents the major phylogenetic clades. *TOP*: As depicted on top of the heatmap, the chemical information underwent a hierarchical clustering according to the given phylogeny to reveal chemo-evolutionary patterns. *BOTTOM*: Selected chemical classification generated by NAP based dereplication are displayed below in blue, while the presence of level 1 identification (m/z, retention time and MS^2^ match) in a particular chemical family is shown in gold. *RIGHT*: Functional annotations including the presence of leaf resin and hairiness, pollination, antibacterial activity, traditional medicinal usage as well as biogeographical species distribution information are displayed on the right side. A summary of legends are also included to the right of the figure.

**Figure S6.**
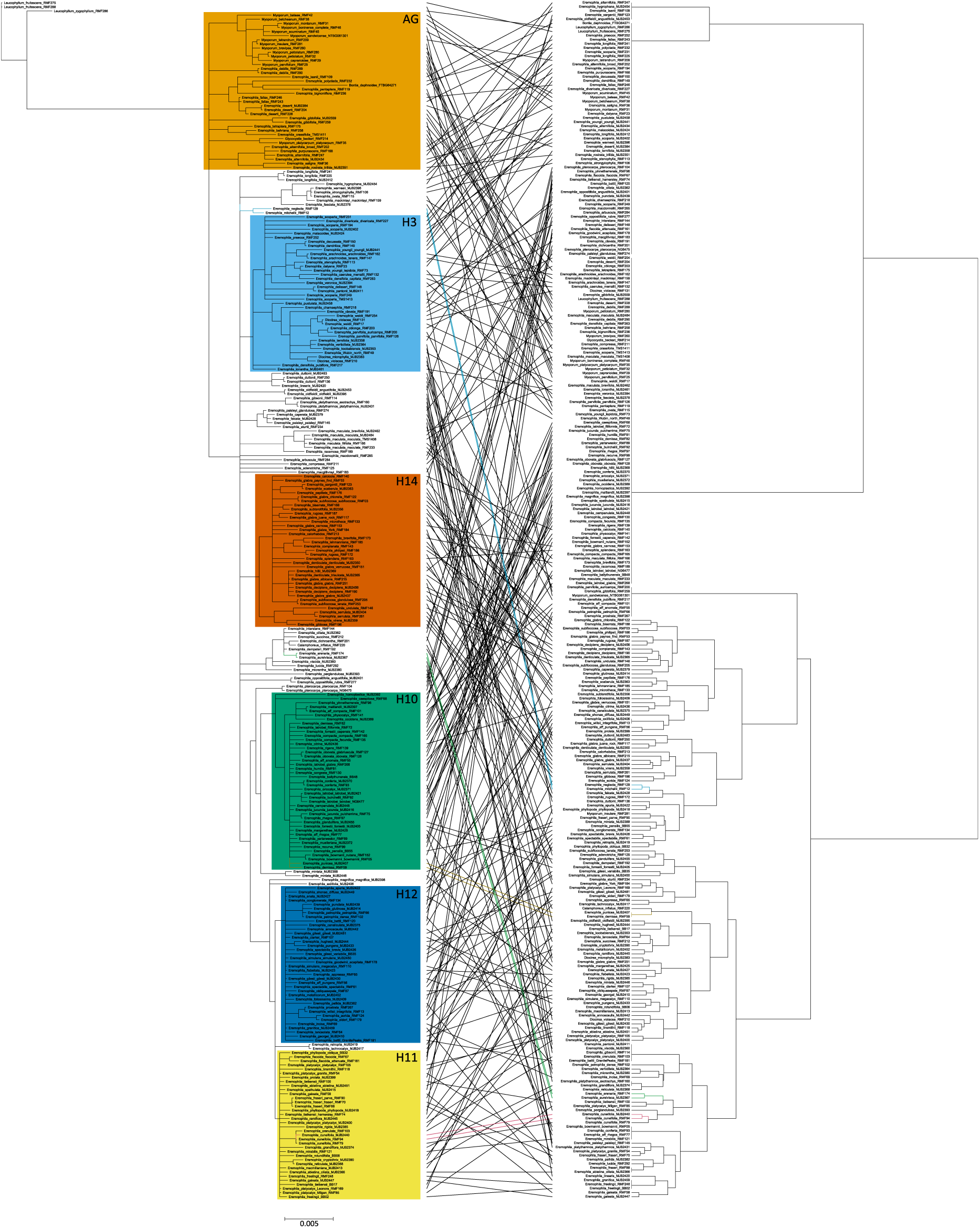
Tanglegram analysis based on serrulatane and viscidane related chemical families. Conjunction of phylogenetic and metabolic information from 291 specimen of the tribe Myoporeae visualized by a tanglegram. The nrDNA based phylogeny shows taxa with posterior probability ≥ 95%. A metabolic cluster analysis was conducted based on the presence or absence of the 30 chemical families found to be associated with serrulatane or viscidane metabolism within the generated global molecular network of Myoporeae. Phylogenetic subclades are indicated with the most prevalent ones highlighted by color code. The tanglegram analysis connects same specimens by a line, which is colored when equal branching of taxa is present in both analyses.

**Figure S7.**
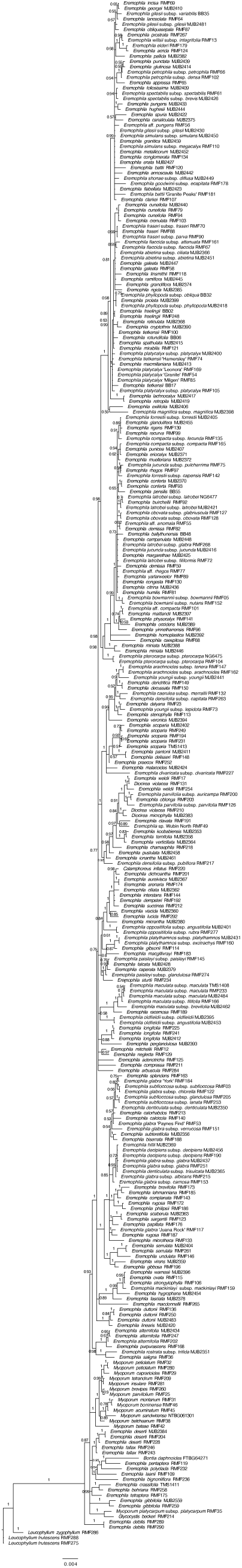
Phylogeny of tribe Myoporeae based on analysis of nuclear ribosomal DNA sequences. Bayesian inference 50% majority-rule consensus tree, showing posterior probability values on each branch. Branches with values <0.50 collapsed.

**Figure S8.**
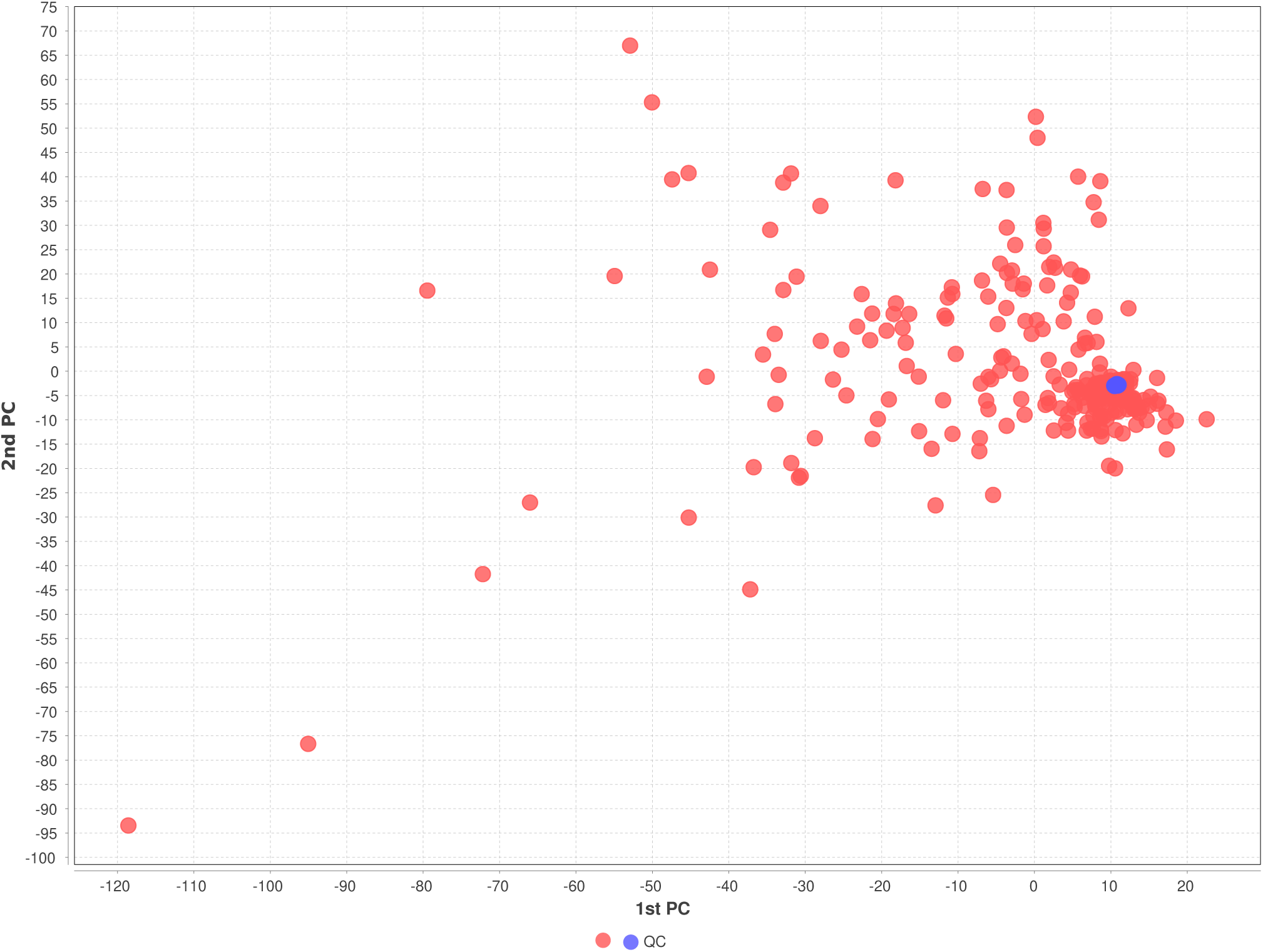
LC-qToF-HRMS data quality control. Assessing quality of the full spectral dataset used in this study by plotting all samples (red) together with the quality controls (blue) within a principal component analysis.

**Figure S9.**
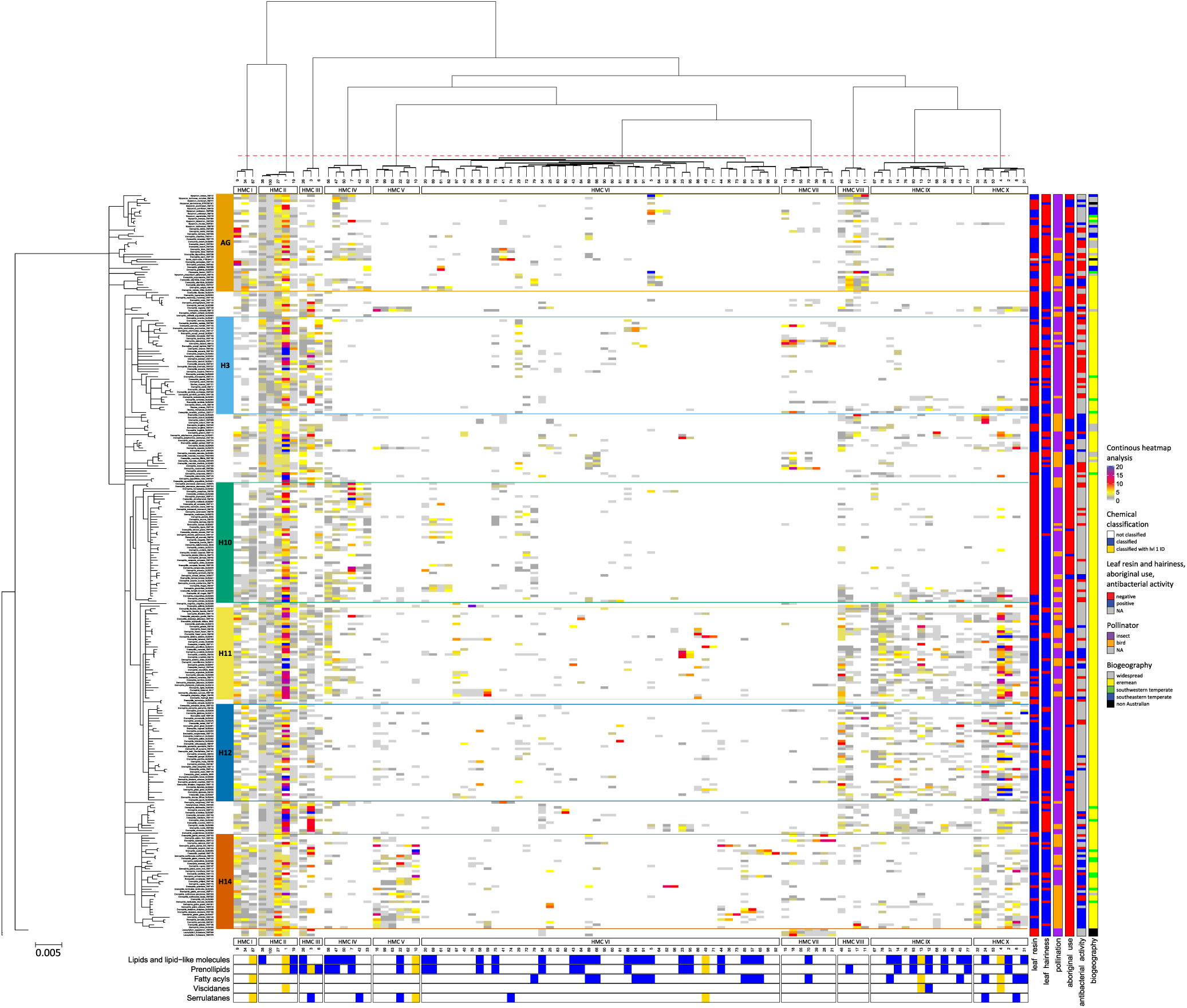
Continuous heatmap analysis showing chemo-evolutionary relationships in Myoporeae. A heatmap analysis was used to put chemical information into an evolutionary context. For that, information about the 100 largest chemical families was compiled, which derived from the molecular network of Myoporeae. For each specimen in this study, the total amount of metabolites of a corresponding chemical family is displayed within the heatmap. *LEFT*: The nuclear ribosomal DNA phylogeny on the left side presents the major phylogenetic clades. *TOP*: As depicted on top of the heatmap, the chemical information underwent a hierarchical clustering according to the given phylogeny to reveal chemo-evolutionary patterns among subsets of chemical families, defined as heatmap metabolic clusters (HMC) I – X. *BOTTOM*: Selected chemical classification generated by NAP based dereplication are displayed below in blue, while the presence of level 1 identification (m/z, retention time and MS2 match) in a particular chemical family is shown in gold. *RIGHT*: Functional annotations including the presence of leaf resin and hairiness, pollination, antibacterial activity, traditional medicinal usage as well as biogeographical species distribution information are displayed on the right side. A summary of legends are also included to the right of the figure.

**Figure S10.**
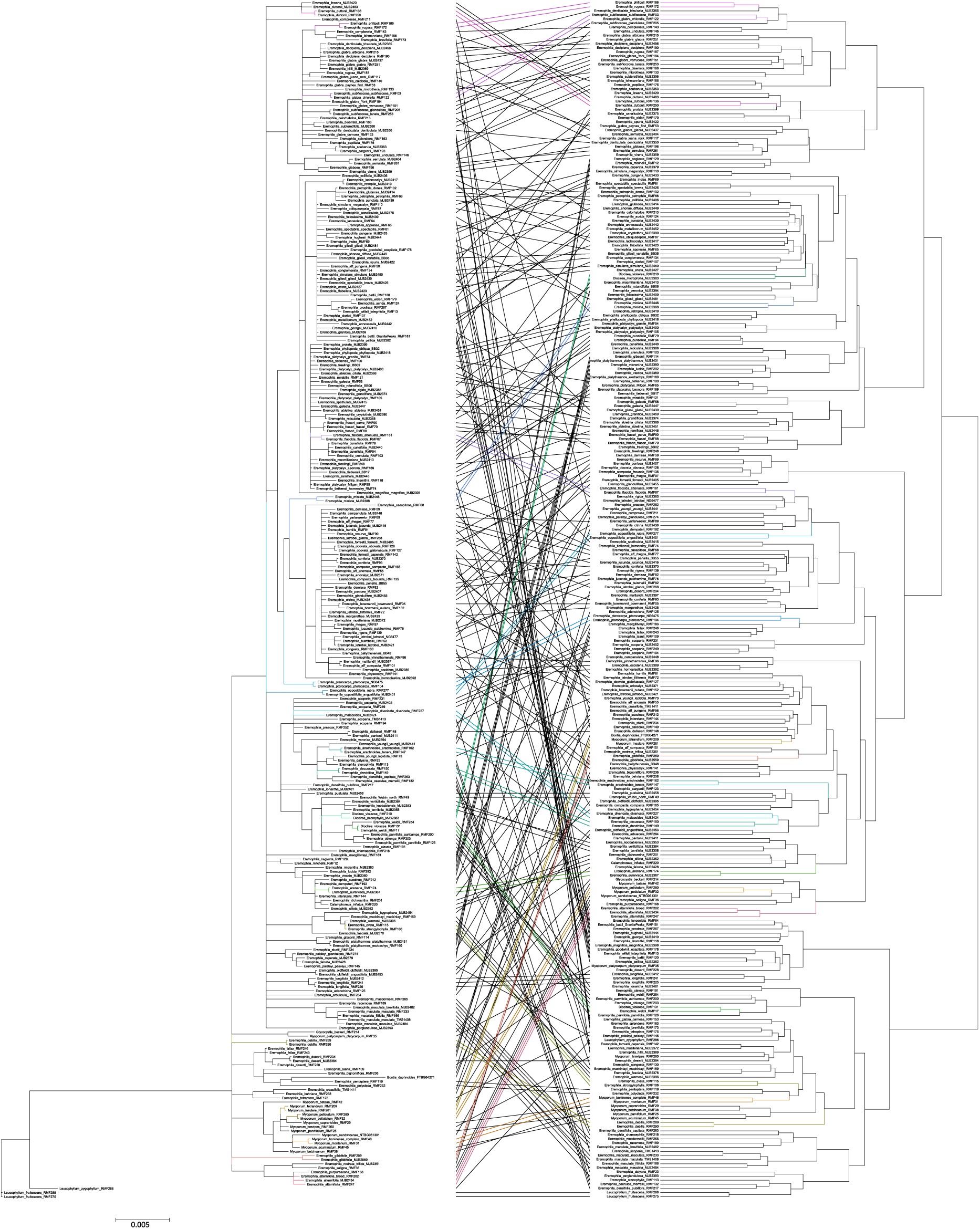
Metabolite based tanglegram analysis showing chemo-evolutionary relationships in Myoporaea. Conjunction of phylogenetic and metabolic information from 291 species of the tribe Myoporeae visualized by a tanglegram. The nrDNA based phylogeny shows taxa with posterior probability above 95%. A metabolic cluster analysis was conducted based on the presence or absence of respective metabolites after normalization of the full spectral dataset (10696 metabolites). Phylogenetic subclades are indicated with the most prevalent ones highlighted by color code. The tanglegram analysis connects same specimens by a line, which is colored when equal branching of taxa is present in both analyses.

## Notes

### Competing Interest Statement

The authors have declared no competing interest.

